# Biofoundry-assisted expression and characterisation of plant proteins

**DOI:** 10.1101/2021.03.11.434954

**Authors:** Quentin M. Dudley, Yao-Min Cai, Kalyani Kallam, Hubert Debreyne, Jose A. Carrasco Lopez, Nicola J. Patron

## Abstract

Many goals in synthetic biology, including the elucidation and refactoring of biosynthetic pathways and the engineering of regulatory circuits and networks, require knowledge of protein function. In plants, the prevalence of large gene families means it can be particularly challenging to link specific functions to individual proteins. However, protein characterisation has remained a technical bottleneck, often requiring significant effort to optimise expression and purification protocols. To leverage the ability of biofoundries to accelerate design-built-test-learn cycles, we present a workflow for automated DNA assembly and cell-free expression of plant proteins that accelerates optimisation and enables rapid progression to characterisation. First, we developed a phytobrick-compatible Golden Gate DNA assembly toolbox containing plasmid acceptors for cell-free expression using *E. coli* or wheat germ lysates as well as a set of N- and C-terminal tag parts for detection, purification, and improved expression/folding. We next optimised automated assembly of miniaturised cell-free reactions using an acoustic liquid handling platform and then compared tag configurations to identify those that increase expression. We additionally developed a luciferase-based system for rapid quantification that requires a minimal 11 aa tag and demonstrate facile removal of tags following synthesis. Finally, we show that several functional characterisation experiments can be performed with cell-free protein synthesis reactions without the need for protein purification. Together, the combination of automated assembly of DNA parts and cell-free expression reactions should significantly increase the throughput of experiments to test and understand plant protein function and enable the direct reuse of DNA parts in downstream plant engineering workflows.

## 1. Introduction

Plant synthetic biology endeavours to apply principles of abstraction, modularity, and standardisation to engineer plants for useful purposes (1, 2). In support of this goal, the plant community adopted a common syntax, commonly known as the ‘phytobrick’ standard, that defines the features of DNA parts (3, 4). For expression in plants, phytobricks can be assembled into synthetic genetic circuits using a number of plasmid toolkits including MoClo (5), Loop (6), GoldenBraid (7), and Mobius (8). These systems use Type IIS restriction endonucleases to direct one-pot digestion-ligation assembly reactions, known as Golden Gate (9), that are easily parallelised and miniaturised for automation (10, 11). Parallel DNA assembly has become an enabling technology for biofoundries, which specialise in automating the design–build–test–learn (DBTL) cycle that underpins synthetic biology (12-14).

Plants have long been exploited as sources of bioactive and high-value natural products (15) and, recently, applications of model-informed synthetic biology approaches have led to sophisticated engineering of crop traits including biomass and responses to environment (16-18). There is also growing interest and investment in the use of plants, particularly *Nicotiana benthamiana*, as photosynthetic platforms for the production of recombinant proteins and small molecules for industry and medicine (19-21), including rapid-response vaccines (22-25). For many applications, methods to rapidly characterise protein functions are essential. This particularly applies to understanding the specific functions of members of large protein families such as decorating enzymes and transcription factors. However, for plant proteins this is frequently a bottleneck, with researchers often investing considerable time and effort to identify permissive constructs and conditions to obtain useful yields before lengthy protocols to purify recombinant proteins for *in vitro* assays.

Cell-free protein synthesis (CFPS) is an established tool for rapid *in vitro* protein production that combines a DNA template, energy source, amino acids, NTPs, and excess cofactors, along with a crude lysate containing the translational machinery (26, 27). The source lysate from *Escherichia coli* is highly active, easy to manipulate, and can produce a broad range of proteins (28) including enzymes (29, 30), antibodies (31), glycoproteins (32, 33), as well as proteins containing non-canonical amino acids (34-36). It is also possible to use plant-based lysates from wheat germ (37), Arabidopsis (38), and BY-2 tobacco cells (39). Cell-free expression can minimise “build” times in the DBTL cycle (40) and has been used to prototype metabolic pathways (29, 30, 41, 42) and characterise several natural product biosynthesis pathways (43, 44) including cyanobacterial alkaloids (45) and antibiotic peptides (46). Furthermore, cell-free protein synthesis is amenable to automation and miniaturisation (47). The use of automation platforms that utilise acoustic energy to transfer reagents have been shown to reduce operator error and variability in CFPS reactions (48), facilitate active learning-guided optimisation of reaction conditions (49), and generally increase the throughput of experiments (50-55).

In this work, we have developed molecular tools and automated biofoundry workflows to enable cell-free expression of plant-proteins (**Figure 1A**). We first developed a phytobrick-compatible plasmid toolkit including (1) plasmid acceptors containing regulatory elements for T7-driven *E. coli* CFPS and the commercial TNT SP6 Coupled Wheat Germ expression system (Promega) (2) affinity tags for purification or detection, and (3) a suite of tags for improving the yields of soluble protein. We then use the toolkit to optimise the expression of a range of plant proteins including enzymes and transcription factors. Finally, we demonstrate the functional activity of the cell-free expressed proteins. Together, our tools and workflows enable the rapid optimisation of expression conditions and allow immediate progression to characterisation assays, significantly increasing the scale and throughput of experimentation.

**Figure 1.**
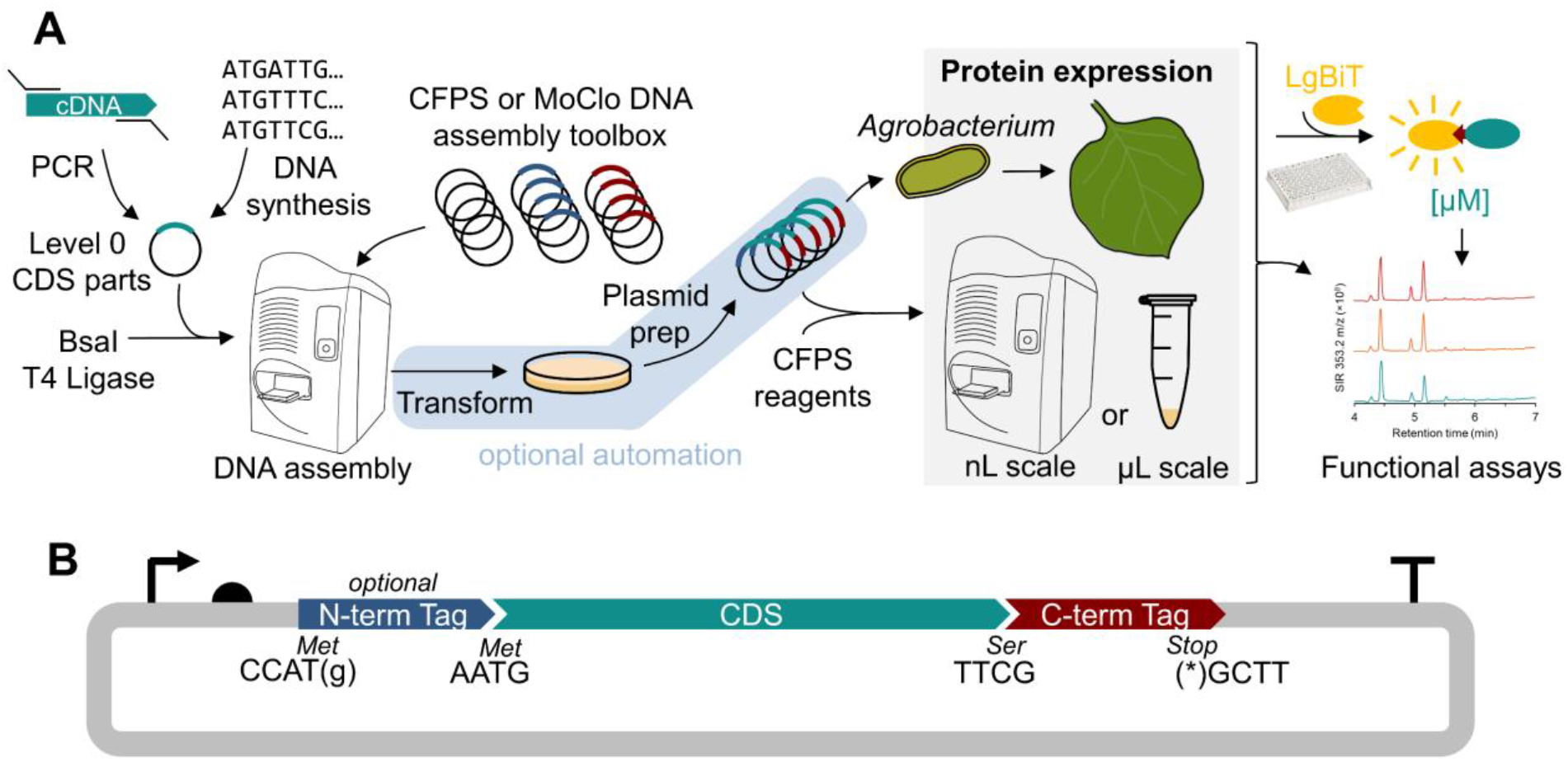
A workflow for biofoundry-assisted DNA assembly and cell-free protein synthesis (CFPS) of plant proteins. (A) Level 0 DNA parts (phytobricks) encoding the protein of interest can be assembled into functional expression plasmids for CFPS or expression *in planta*. Acoustic liquid handling enables the screening of libraries of protein variants to determine optimal expression configurations using the HiBiT-LgBiT luminescence. (B) The CFPS cloning toolbox consists of acceptor plasmids that assemble with a CDS and C-terminal tag along with an optional N-terminal tag. The cloning overhangs are compatible with existing plant DNA assembly standards

## 2. Methods

### 2.1 DNA assembly

Acceptor plasmids for CFPS (see **Table 1**) were designed as “terminal acceptors” and are not designed for subsequent assembly into plasmids containing multiple transcriptional units; each contains promoter and terminator sequences specific to the expression system being utilised. Acceptors for *E. coli* CFPS were based on pJL1 (Addgene #69496) and some include an N-terminal Expression Tag (NET) which encodes Met-Glu-Lys-Lys-Ile (MEKKI) shown to enhance expression of some proteins (29). Acceptors for wheat germ expression were based on pEU-Gm23 (Addgene #53738) which uses the pEU backbone (56) containing the E01 translational enhancer (57).

**Table 1.** A plasmid toolkit for cell-free protein expression compatible with the phytobrick assembly standard. Each fusion site lists the top strand only based on the orientation shown in **Figure 1B**.

| Plasmid code | Addgene number | Purpose | Description | Fusion sites |
| --- | --- | --- | --- | --- |
| pEPQD0KN0024 | 162281 | acceptor, *E. coli* CFPS | T7 promoter, RBS, N-term exp. tag, T7 terminator, kanR, pUC ori | AATG GCTT |
| pEPQD0KN0244 | 162282 | acceptor, *E. coli* CFPS | T7 promoter, RBS, N-term exp. tag, T7 terminator, kanR, pUC ori | CCAT GCTT |
| pEPQD0KN0025 | 162283 | acceptor, *E. coli* CFPS | T7 promoter, RBS, T7 terminator, kanR, pUC ori | AATG GCTT |
| pEPQD0KN0245 | 162284 | acceptor, *E. coli* CFPS | T7 promoter, RBS, T7 terminator, kanR, pUC ori | CCAT GCTT |
| pEPQD0CB0026 | 162285 | acceptor, TNT SP6 WG | SP6 promoter, EO1 enhancer, folA terminator, ampR, pUC ori | AATG GCTT |
| pEPQD0CB0246 | 162286 | acceptor, TNT SP6 WG | SP6 promoter, EO1 enhancer, folA terminator, ampR, pUC ori | CCAT GCTT |
| pEPQD0KN0282 | 162287 | acceptor, TNT SP6 WG | SP6 promoter, EO1 enhancer, T7 terminator, kanR, pUC ori | AATG GCTT |
| pEPQD0KN0283 | 162288 | acceptor, TNT SP6 WG | SP6 promoter, EO1 enhancer, T7 terminator, kanR, pUC ori | CCAT GCTT |
| pEPQD0KN0284 | 162289 | acceptor, TNT SP6 WG | SP6 promoter, EO1 enhancer, folA terminator, kanR, pUC ori | AATG GCTT |
| pEPQD0KN0285 | 162290 | acceptor, TNT SP6 WG | SP6 promoter, EO1 enhancer, folA terminator, kanR, pUC ori | CCAT GCTT |
| pEPMY1CB0001 | 162317 | acceptor, *E. coli* | T7 promoter, lac operator, RBS, T7 terminator, ampR, pUC ori | CCAT GCTT |
| pEPYC0CM0258 | 162291 | N-terminal tag | HiBiT | CCAT AATG |
| pEPQD0CM0541 | 162292 | N-terminal tag | GST-[thrombin cleavage site] | CCAT AATG |
| pEPQD0CM0542 | 162293 | N-terminal tag | HiBiT-GST-[thrombin cleavage site] | CCAT AATG |
| pEPQD0CM0543 | 162294 | N-terminal tag | GST-[TEV cleavage site] | CCAT AATG |
| pEPQD0CM0544 | 162295 | N-terminal tag | MBP-[Factor Xa cleavage site] | CCAT AATG |
| pEPQD0CM0545 | 162296 | N-terminal tag | HiBiT-MBP-[Factor Xa cleavage site] | CCAT AATG |
| pEPQD0CM0546 | 162297 | N-terminal tag | MBP-[TEV cleavage site] | CCAT AATG |
| pEPQD0CM0547 | 162298 | N-terminal tag | TrxA-[TEV cleavage site] | CCAT AATG |
| pEPQD0CM0548 | 162299 | N-terminal tag | HiBiT-TrxA-[TEV cleavage site] | CCAT AATG |
| pEPQD0CM0549 | 162300 | N-terminal tag | SUMO-[TEV cleavage site] | CCAT AATG |
| pEPQD0CM0550 | 162301 | N-terminal tag | HiBiT-SUMO-[TEV cleavage site] | CCAT AATG |
| pEPQD0CM0281 | 162302 | N-terminal tag | [S-tag]-[TEV cleavage site] | CCAT AATG |
| pEPQD0CM0551 | 162303 | N-terminal tag | HiBiT-[S-tag]-[TEV cleavage site] | CCAT AATG |
| pEPMY0SP0002 | 162304 | N-terminal tag | 6xHis tag-[HRV 3C cleavage site] | CCAT AATG |
| pEPQD0CM0296 | 162305 | CDS | superfolder GFP, no stop codon | AATG TTCG |
| pEPQD0CM0539 | 162306 | CDS | TEV protease (S219V), no stop codon | AATG TTCG |
| pEPYC0CM0134 | 162307 | C-terminal tag | HiBiT-stop, frame1 (NNT-TCG) | TTCG GCTT |
| pEPYC0CM0257 | 162308 | C-terminal tag | HiBiT-stop, frame2 (TTC-G) | TTCG GCTT |
| pEPQD0CM0027 | 162309 | C-terminal tag | 9 aa linker, superfolder GFP, strep tag, stop codon | TTCG GCTT |
| pEPQD0CM0028 | 162310 | C-terminal tag | 9 aa linker, strep tag, stop codon | TTCG GCTT |
| pEPQD0CM0029 | 162311 | C-terminal tag | strep tag, stop codon | TTCG GCTT |
| pEPQD0CM0030 | 162312 | C-terminal tag | stop codon | TTCG GCTT |
| pEPQDKN0248 | 162313 | reporter, *E. coli* CFPS | T7prom-RBS-sfGFP-HiBiT | n/a n/a |
| pEPQDKN0332 | 162314 | reporter, TNT SP6 WG | SP6prom-EO1-sfGFP-HiBiT | n/a n/a |
| pEPQDKN0329 | 162315 | protease, *E. coli* CFPS | T7prom-RBS-NET-TEVprotease-HiBiT | n/a n/a |
| pEPQDKN0729 | 162316 | protease, *E. coli* CFPS | T7prom-RBS-NET-TEVprotease | n/a n/a |

New level 0 DNA parts (i.e., N-terminal tags, coding sequences (CDS), C-terminal tags) were either synthesised (Twist Bioscience, San Francisco, CA) or amplified by PCR with overhangs containing BpiI (BbsI) recognition sites and assembled into pUAP1 (Addgene #63674) to create parts compatible with the phytobrick standard (3). We included a number of N-terminal tags previously shown to improve translation, solubility, and folding (58, 59): the S-tag sequence pancreatic ribonuclease A (KETAAAKFERQHMDS) typically used for quantitation/purification but anecdotally suggested to improve protein solubility (60); a small ubiquitin-related modifier (SUMO) sequence known to increase soluble expression (61) derived from pDest-Sumo (Addgene #106980) (62); a TrxA sequence from *E. coli* and derived from pET32a-TRXtag (Addgene #11516) (63); a glutathione-S-transferase (GST) (64) and thrombin sequence derived from pGEX-2T (GE Healthcare, Chicago, IL), and a maltose binding protein (MBP) (65) and Factor Xa sequence derived from pMAL-c5X vector (New England Biolabs, Ipswich, MA) with a V313A mutation to be consistent with the native *E. coli* sequence. We also included N-terminal and C-terminal HiBiT tags (Promega, Madison, WI) for quantification using luminescence, a green florescence protein (GFP) derived from pJL1 (Addgene #69496), and a C-terminal twin-strep-tag (66). Finally, we included a CDS part encoding TEV protease containing a S219V mutation reported to have less autolysis (67).

All DNA assembly reactions were performed in 20 μL (manual) or 2 µL (automated) reaction volumes as previously described (10) and were verified by sequencing.

### 2.2 Cell-free protein synthesis

All *E. coli* CFPS reactions used a modified PANOx-SP formula (68, 69) as previously described (29) with minor modifications. Lysate was generated from BL21(DE3) *E. coli* cells and S30 buffer used for lysate prep contained 2 mM dithiothreitol (DTT). Ammonium glutamate was not available and not included in the CFPS reaction. Optimal magnesium concentration was determined to be 8 mM based on expression of the plasmid pJL1-sfGFP (Addgene #69496) encoding superfolder Green Fluorescent Protein (sfGFP).

Reactions were assembled in a final volume of 2 μL using a Labcyte Echo® 550 (Beckman Coulter, Brea, CA) via a two-step strategy. Initially, 30-50 μL of each plasmid (or water) were distributed into an Echo® Qualified 384-Well Polypropylene Source Microplate (384PP, P-05525). Subsequently, 65 μL of a master mix containing all remaining CFPS reagents (with lysate added just before distributing) was aliquoted to fresh wells of the same source plate. Next, the Labcyte Echo® Plate Reformat software v1.6.4 was used to direct the distribution of 1735 nL to each well of a FrameStar® 384 Well PCR Plate (4ti-0384/C; 4titude, Wotton, UK) (i.e., the destination plate) using 384PP_AQ_SP2 as sample plate type setting. Each source well can support 20-21 reactions. Then, the Labcyte Echo® Cherry Pick software v1.6.4 (guided by a .csv file) used the 384PP_AQ_BP2 sample plate type setting to direct the distribution of 265 nL containing 26.6 ng plasmid (equivalent to a final concentration of 13.3 ng/μL) with remaining volume water to all destination wells. The destination plate was then centrifuged briefly at room temperature, covered with adhesive aluminium foil, and incubated at 30 °C for 20 hours (lid temperature 37 °C) in an Mastercycler pro384 vapo.protect thermocycler (Eppendorf, Hamberg, Germany). Scaled-up (15 μL) CFPS reactions were performed in 1.5 mL DNA LoBind® microcentrifuge tubes (0030108051; Eppendorf). To quantify soluble/total protein, reactions were centrifuged for 10 minutes at 4 °C at 21,000 *x g* to pellet insoluble protein.

All wheat germ cell-free expression used the TNT® SP6 High-Yield Wheat Germ Protein Expression System L3260/L3261 (Promega) according to manufacturers’ instructions. Plasmids were added at a final concentration of 80 ng/μL and assembled reactions were incubated at 25 °C for 20 hours.

### 2.3 Protein quantification

To compare relative protein concentration of proteins containing GFP, 2 μL of CFPS reaction were diluted with 198 μL of 10mM Tris-HCl pH 7.5 in a black 96-well plate (655076 PS medium binding; Greiner Bio-One Vilvoorde, Belgium). After a double orbital shake for 10 seconds at 300 rpm, fluorescence was measured (excitation 470 nm, emission 515 nm) using a CLARIOstar plate reader (BMG Labtech Gmbh, Ortenberg, Germany).

Absolute protein concentration of HiBiT-tagged proteins was measured using the Nano-Glo® HiBiT Extracellular Detection System N2420 (Promega). CFPS reactions were diluted 104-106 -fold using 1X PLB+PI buffer (generated by mixing 2 mL of 5x Passive Lysis Buffer E1941 (Promega) with 8 mL water plus one cOmplete™ Mini EDTA-free 11836170001 (Roche, Basel, Switzerland) protease inhibitor tablet). A standard curve of HiBiT Control Protein N3010 (Promega) was diluted in PLB+PI at concentration ranging from 0.01 to 2 nM. Luminescence reactions were mixed in a white 96-well plate (Greiner Bio-One 655075 PS medium binding) by combined 40 μL of diluted protein (CFPS reaction or standard protein) with 10 μL of luminescence reagent master mix (containing 10 μL of LgBiT protein, 20 μL of 50X Substrate, and 970 μL of HiBiT buffer). Luminescence values was recorded in a CLARIOstar plate reader every 2.33 minutes for an hour to ensure the signal is stable over time with the value at 18 minutes typically used for sample comparison. CFPS reactions assembled on the Labcyte Echo® 550 were diluted 102-105 -fold using 10mM Tris-HCl pH 7.5 and frozen at -20 °C; 4 μL was later thawed and added to 36 μL of PLB+PI along with 10 μL luminescence reagent master mix for measurement.

### 2.4 TEV protease cleavage of N-terminal tags and detection by Western blot

Plasmids pEPQDKN0329 (with C-terminal HiBiT tag) and pEPQDKN0729 (no C-terminal tag) were expressed using standard CFPS conditions and pEPQDKN0329 amount quantified via HiBiT. CFPS reactions of pEPQDKN0729 were mixed with CFPS reactions containing 2.5 μg protein of pEPQDKN0313 (MBP-sfGFP), pEPQDKN0310 (GST-sfGFP), pEPQDKN0314 (TrxA-sfGFP), pEPQDKN0316 (SUMO-sfGFP) and pEPQDKN0318 (S-tag-sfGFP) at a w/w ratio of 1:130. CFPS reaction incubated without a plasmid filled remaining volume to 5 μL. Cleavage reactions were incubated at 30 °C for 16 hours, diluted 1:200 in 10 mM Tris-HCl pH 7.5 and 9 μL loaded (along with 1 μL 1 M DTT and 10 μL 2X Laemmli Buffer) into an Any kD™ Mini-PROTEAN® TGX™ Precast Protein Gel (Bio-rad). For Western blot, proteins were transferred to a PVDF membrane, incubated with α-GFP-HRP (Santa Cruz Biotech GFP, sc-9996 HRP), and imaged using SuperSignal West Pico chemiluminescent substrate (ThermoFisher, Waltham, MA).

### 2.5 Purification of His-tagged UGT73C5

Plasmid pEPQDCB0093 was transformed into BL21(DE3) *E. coli* cells. A starter culture of 50 mL of LB media containing 100 µg/mL carbenicillin was inoculated with 2 mL of saturated overnight culture. Cells were grown at 37 °C (250 rpm) until OD_600_ ∼ 1.5 to 3 and used to inoculate 1000 mL of 2YT media containing 100 µg/mL carbenicillin to a calculated initial OD_600_ of 0.037. Cells were further grown at 37 °C (250 rpm) until OD_600_ = 0.6 and transferred to 18 °C shaking incubator and allowed to cool. After 1 hour at 18 °C (250 rpm), 0.5 mM IPTG was added to induce protein expression and cells incubated at 18 °C (250 rpm) overnight. Cells were pelleted by centrifugation at 8,000 *x g* for 15 minutes at 4 °C and resuspended in 1X PBS. Cells were centrifuged again at 4,000 *x g* for 15 minutes at 4 °C, supernatant poured off, and flash frozen on liquid nitrogen for storage at -80 °C. After thawing, cells were resuspended in 50 mL Buffer A (50 mM Tris-HCl pH 8, 50 mM glycine, 500 mM NaCl, 5% (v/v) glycerol, 20 mM imidazole) supplemented with 10 mg lysozyme (Sigma, 62971) and 1 tablet of cOmplete™ EDTA-free protease inhibitor (Roche, 11873580001). Cell were lysed by a single run through a Cell Disruption System CF1 (Constant Systems Limited, Daventry, UK) cell disruptor at 26 kpsi and centrifuged at 40,000 *x g* for 30 minutes at 4 °C to remove debris. Protein was purified using an AKTA Pure HPLC (GE Healthcare) fitted with a HisTrap HP 5-ml column (GE Healthcare) equilibrated with Buffer A. Samples were step-eluted using Buffer B (50 mM Tris-HCl pH 8, 50 mM glycine, 500 mM NaCl, 5% (v/v) glycerol, 500mM imidazole). Eluted protein was further purified on a HiLoadTM 16/600 SuperdexTM 200 column (GE Healthcare) and eluted with Buffer A4 (20 mM HEPES, 150 mM NaCl, pH 7.5). Protein was concentrated using a Vivaspin 500 column (Sartorius, Goettingen, Germany) column following the manufacturer’s instructions.

### 2.6 Enzymatic production of glycosides and detection by liquid-chromatography mass-spectrometry (LC-MS)

*In vitro* reactions were performed using both purified enzymes and CFPS reactions enriched in the glycosyltransferase. For purified enzymes, the 100 μL reaction contained 100 mM Tris-HCl (pH 8.0), 1 mM UDP-glucose, 0.5 mM substrate (geraniol or *cis-trans-*nepetalactol) suspended in methanol, and 1 μM purified AtUGT73C5. Reactions were incubated at 30 °C for one hour. For CFPS-derived enzymes, the 30 μL reaction contained 100 mM Tris-HCl (pH 8.0), 2 mM UDP-glucose, 2 mM substrate (geraniol or *cis-trans-*nepetalactol) suspended in methanol, and 15 μL of CFPS reaction. Geraniol (163333) and UDP-glucose (94335) were purchased from Sigma (St. Louis, MO) and *cis-trans*-nepetalactol from Santa Cruz Biotech (sc-506178). Reactions were quenched using 1X volume of methanol and vortexed for 20 seconds. Reactions were centrifuged at 21,000 x g for 5 minutes at room temperature and 50 μL of supernatant filtered through a 0.22 µm nylon Corning® Costar® Spin-X® filter CLS8169 (Sigma).

For detection of geraniol and nepetalactol glycosides, 2 μL of quenched *in vitro* reaction were injected and separated on a Waters UPLC with an Acquity BEH C18, 1.7 µm (2.1 x 50 mm) column (40 °C) at a flow rate of 0.6 mL/min. Mobile phase A was 0.1 % formic acid and mobile phase B was acetonitrile. A linear gradient from 5% B to 60 % B in 5.5 min and 60% to 100 % B in 0.5 min was applied for compound separation followed by 100 % B for 1 min. The flow was returned to 5% B for 2.5 min to re-equilibrate prior to the next injection. Eluting compounds were subjected to positive ESI and analysed on a Waters Xevo TQ-s (QqQ) using optimised source conditions: cone voltage 30 eV, Capillary voltage, 3.0 kV; source temperature, 150 °C; desolvation temperature, 450 °C; cone gas, 150 L/h; and desolvation gas, 800 L/h. MRM transitions monitored for analysis included geraniol-glucoside-H (317.2 m/z), geraniol glucoside-Na (339.2 m/z), geraniol glucoside-NH_4_ (334.2 m/z), nepetalactol-glucoside-H (331.2 m/z), nepetalactol-glucoside-Na (353.2 m/z), and nepetalactol-glucoside-NH_4_ (348.2 m/z).

### 2.7 Enzymatic production of chrysanthemol and detection by gas-chromatography mass-spectrometry (GC-MS)

CFPS reactions expressing pEPKK1KN0203 encode chrysanthemol diphosphate synthase (CcCPPase) from *Chrysanthemum cinerariaefolium* (P0C565.2) (70). Enzymatic reactions, based on (70, 71), consisted of 13 μL of CFPS reaction (containing 3.30 ± 0.22 μM soluble protein) along with added 35 mM HEPES pH 7.6, 5 mM MgCl_2_, 0.5 mM DTT, 2 mM DMAPP to a final volume of 100 μL. The reaction was incubated overnight at 30°C and then heated at 95°C for 2 min. Glycine (500 mM pH 10.5), 5mM MnCl_2_, and 20 units of calf alkaline phosphatase (NEB) were added to the cooled solution and incubated at 37°C for 1 hour. Approximately 0.1 g of NaCl was added and the terpenoids extracted by addition of 500 μL tert-butyl methyl ether. Compounds were analysed using an HP 6890 gas chromatograph with a 5973 MSD (Hewlett Packard / Agilent, Santa Clara, CA) using 1 μL injection, a Zebron™ ZB-5HT Inferno™ capillary column (30 m × 0.25 mm × 0.10 μm + 5 m Guardian) with split vent, and helium carrier gas at constant flow of 1.0 mL/min. The inlet temperature was 200 °C and initial column temperature held at 40 °C for 0.5 min, increased at 25 °C/min to 70 °C, increased at 3 °C/min to 120 °C, and finally increased at 50 °C/min to 200 °C, which was maintained for 3 more minutes. Mass spectra of relevant peaks were compared with NIST database standards to identify chrysanthemol and lavandulol.

### 2.8 Electrophoretic mobility shift assay

CPFS reactions expressing AtTGA2 (pEPQDKN0742) were buffer exchanged with protein dilution buffer (20 mM Tris-HCl, 50 mM KCl, pH 7.5) using an Amicon Ultra-0.5 centrifugal filter (UFC501024; Merck, Kenilworth, NJ). DNA probes were synthesised as oligos (TGA TFBS 5’-gacccctattgcagctatttcacCTGACGTAAGGGATGACGCACAggccatcacgcagta, Random control 5’-gacccctattgcagctatttcacacataccaacgcttagcgcaatggccatcacgcagta). A primer labelled with Alexa-488 (5’-Alexa488-tactgcgtgatggcc) was designed to anneal to the 3’ end of the probe oligos. A full-length, double-stranded Alexa-488 labelled probe was produced using DNA Polymerase I, Large (Klenow) Fragment (NEB, M0210) following the manufacturers’ protocol. 500 pmol protein and 5.5 pmol probe were mixed with EMSA reaction mixture (20 mM Tris-HCl pH 8.0, 50 mM KCl, 2 mM DTT, 1 mM EDTA, 500 ng poly(dI-dC), 5% glycerol, 0.05% IGEPAL CA630, 0.1 mg/mL BSA). The mixture was incubated at room temperature for 1 hour. Bound and unbound probes were separate on a 6% TBE acrylamide gel (Invitrogen, EC6265BOX) and visualised using a ChemiDocTM Touch imaging system (Bio-Rad, Hercules, CA).

## 3. Results

### 3.1 A modular type IIs cloning toolbox for cell free protein synthesis

To enable a pipeline for cell-free expression of plant proteins, we first built a suite of plasmid vectors amenable to miniaturisation and automation and able to utilise DNA parts in the phytobrick standard (**Figure 1A**). We built plasmid toolkits for two cell-free systems: the *E. coli-*based PANOx-SP (29, 68, 69) (driven by a T7 promoter) and the Promega TNT® SP6 High-Yield Wheat Germ Protein Expression System (driven by the SP6 promoter) (37). Using high copy number plasmid backbones, we constructed a suite of 10 different acceptor plasmids with minimal regulatory sequences and strong terminators for both systems each capable of utilising Level 0 phytobrick N-tag, CDS, and C-tag parts (**Figure 1B, Table 1**). All plasmids have been deposited in the Addgene plasmid repository.

To expand the utility of the DNA assembly toolbox, we made a library of parts encoding N-terminal and C-terminal tags for purification, detection, and improved expression. Facile detection of proteins is important for high-throughput CFPS applications and although superfolder GFP (sfGFP) was optimised to reduce interference with the folding of its fusion partners, bulky fluorescent protein tags are not always tolerated. To accurately measure protein expression without a GFP fusion or using expensive radiolabelled amino acids, we adapted the HiBiT system sold by Promega. This requires only a minimal 11 amino acid tag to be genetically encoded. Following protein expression, an 18 kDa engineered polypeptide with high-affinity for the tag is added resulting in a complex with luciferase activity proportional to the level of HiBiT-tagged protein. To assess the ability of HiBiT to quantify protein in CFPS reactions, we measured *E. coli* cell-free expression of superfolder GFP (sfGFP) fused with a C-terminal HiBiT tag (pEPQDKN0248) to be 41.8 ± 3.0 μM (1140 ± 84 μg/mL) (**Figure 2A**, second bar from top), which is consistent with other PANOx-SP systems using BL21(DE3) (69, 72). The HiBiT tag works for quantification on both the N-and C-terminus of a target protein, however, we found that inclusion on the N-terminus reduced cell-free protein yield when measuring relative GFP fluorescence (**Figure 2A**).

**Figure 2.**
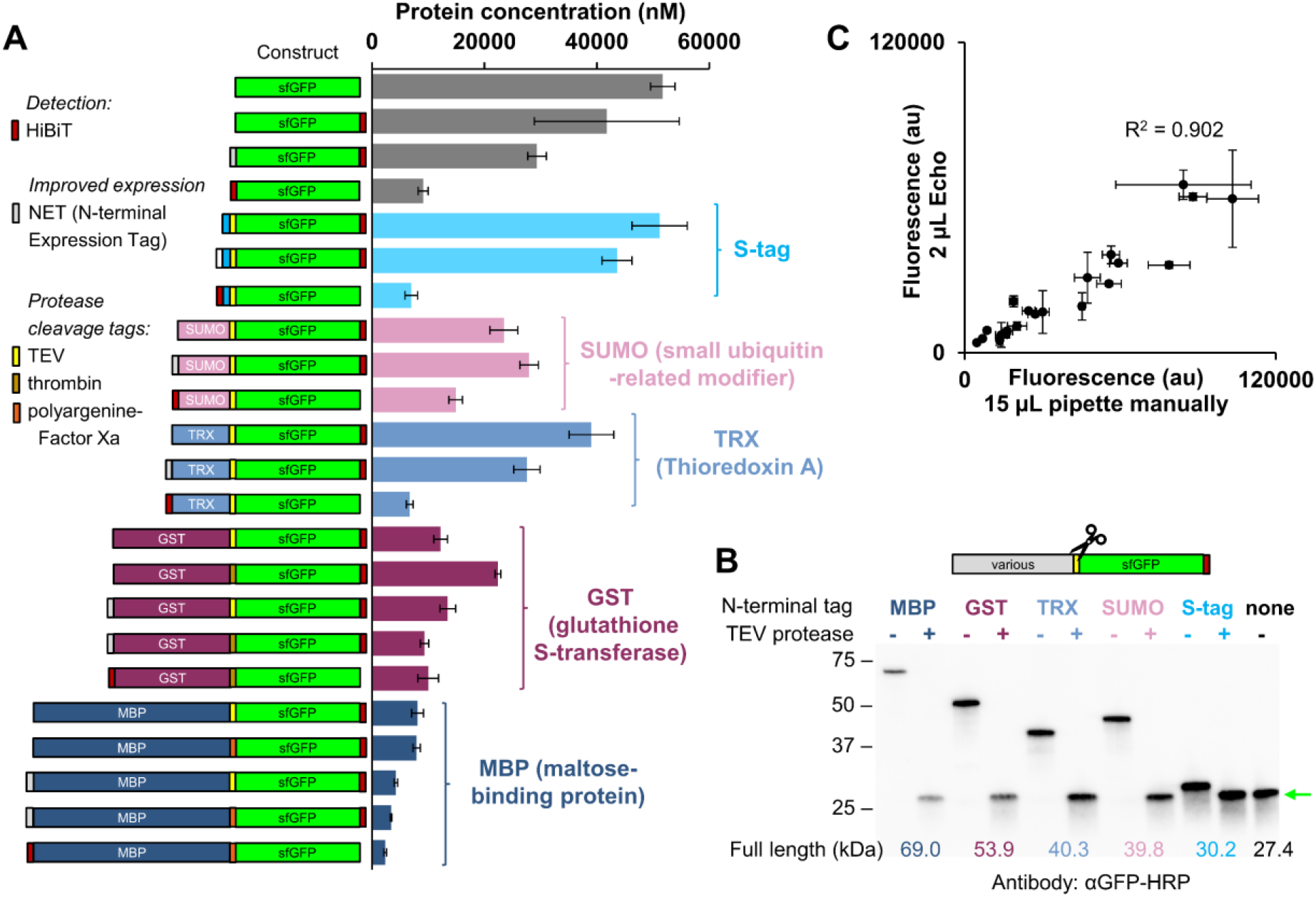
(A) Cell-free expression of sfGFP fused to a variety of N-and C-terminal tags. CFPS reactions were run at 15 µL scale and incubated for 20 h at 30 °C. Relative protein expression was measured by GFP fluorescence and normalised to pEPQDKN0248 quantified by HiBiT. (B) Removal of N-terminal tags by mixing a CFPS reaction expressing TEV protease (1:130 w/w ratio of TEV protease to substrate protein) and incubating at 16 h at 30 °C. Protein visualised by Western Blot with an α-GFP-HRP antibody. (C) Expression of GFP variants from panel A with reactions assembled manually using handheld pipettes (15 μL reactions) or automated using an Echo 550 acoustic liquid handler (2 μL reactions). Values in panel A and C represent averages (n=3) and error bars represent 1 standard deviation.

Several fusion tags have been shown to improve translation, solubility, and folding of recombinant proteins, however, it is rarely obvious which expression tag is optimal for a given protein. We therefore included a number of different tags in the toolkit. We initially assessed the functionality of S-tag, SUMO, thioredoxin (TRX), glutathione S-transferase (GST), and maltose-binding protein (MBP) tags by assembling them with the sfGFP CDS and doing *E. coli* CFPS reactions (**Figure 2A**). We found that many N-terminal tags appeared to express well, though the larger tags (such as MBP) reduced expression. As expected, they did not increase the expression of sfGFP, for which the CFPS reaction conditions have been optimised (34, 69).

As many downstream protein applications require removal of tags that may interfere with, for example, enzyme functionality, we developed a low-cost method for tag cleavage. First, we expressed the TEV protease S219V using CFPS. Subsequently, this reaction was mixed with the reaction containing the target protein containing the cleavage site (**Supplementary Figure S1**). Mixing the two reactions overnight at a 1:130 w/w ratio of protease:target (∼1:2 v/v ratio of cell-free reactions) showed efficient cleavage when assayed by Western Blot using an anti-GFP antibody (**Figure 2B**).

### 3.2 Sequential improvement of automated cell-free protein synthesis

To enable the progression of high throughput combinatorial experiments, for example, to select tags that enable the best yields for individual proteins, we established an automated workflow for low-volume CFPS. Our aim was to achieve consistent results across replicates and to ensure that the yields obtained from low-volume reactions correlated with those from large-scale reactions, thus demonstrating that conditions established using biofoundries are transferable. To maximise consistency, we pre-mixed all cell-free reaction components into a master mix, which was distributed to all wells of a 384-well plate to which the plasmid template was subsequently added. We first found that calibration of plate type settings was required to enable nanovolumes of master mix to be accurately transferred by the acoustic droplet technology of the Labcyte Echo (**Supplementary Figure S2A and B**). We then optimised the number of destination wells to which reaction mix was transferred from each source well to reduce the number of failed reactions (**Supplementary Figure S2C**). Finally, to reduce the time taken for 384 CFPS reactions to be assembled while maximising the consistency and yield of protein between reactions, we compared four different reaction volumes (**Supplementary Figure S2D**) as well software protocols that monitor reagent transfer (**Supplementary Figure S2E**).

In summary, we found that a single specific plate setting (384PP_AQ_SP2) was essential for consistent distribution, and that expression levels obtained from 2000 nL reactions were the most consistent and produced the largest amount of protein. Expression of GFP fusion protein was found to be consistent across all wells of a 384-well plate and the yields obtained correlated well (R2 = 0.902) with assembly by manual pipette (15 μL) (**Figure 2C**). This suggests that automated reaction assembly is an appropriate method for quantifying and comparing expression from different DNA templates.

### 3.3 Selecting optimal configurations for expression of plant proteins

To test our DNA and cell-free assembly workflow on non-model coding sequences, we first selected a range of plant UDP glycosyltransferases (UGTs). UGTs are a superfamily of enzymes that catalyse the addition of glycosyl groups, including to many plant specialised metabolites. Although UGTs can easily be identified from their primary sequence, functional assays are typically required to identify which family members are active on a specific substrate. We included UGT73C5 from *Arabidopsis thaliana*, together with ten uncharacterised UGTs from *N. benthamiana*. We constructed eight different versions of eleven plant UGTs (88 total plasmids), assembled 2000 nL cell-free reactions, and measured total protein level using the HiBiT assay (**Figure 3A**). In general, we observed that SUMO and S-tags improved the expression of most UGTs, however, several did not express well in any context and may require further optimisation or an alternate expression strategy. The GST fusion produced high protein levels for only a handful of UGTs (UGT74T6). Finally, we manually assembled 15 μL reactions for a dozen different UGT-encoding plasmids and again obtained a strong correlation with the low-volume automated reaction (**Supplementary Figure S3**).

**Figure 3.**
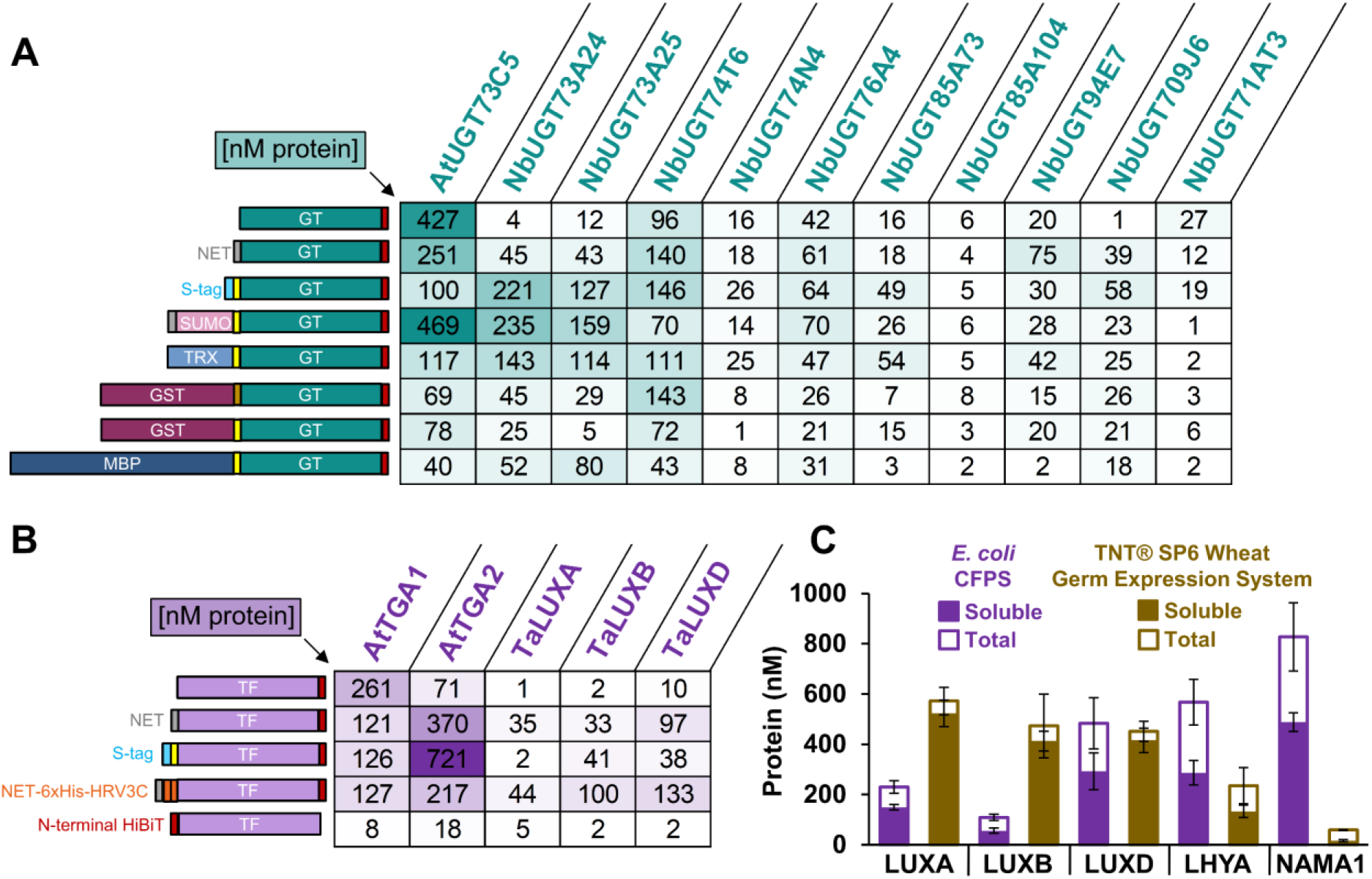
(A-B) HiBiT quantification of plant proteins with various N-terminal tags with cell-free protein synthesis reactions assembled using acoustic liquid handling platforms. (A) Expression of eleven plant UGTs, each with eight different N-terminal tags (B) Expression of five plant transcription factors, each with five different N-terminal tags. (C) Comparison of *E. coli* CFPS (with NET tag) and the TNT® SP6 High-Yield Wheat Germ Protein Expression System (Promega) for expressing wheat transcription factors at 15 μL scale. Values represent averages (n=3) and error bars represent 1 standard deviation.

After demonstrating that biofoundry protocols can be used to select optimal construct configurations, we proceeded to test the expression of different classes of proteins. We first tested additional enzymes. CFPS has previously been demonstrated for monoterpene synthases that catalyse a regular head-to-tail 1-4 linkage, for example, geranyl diphosphate synthase or farnesyl diphosphate synthase (29). We successfully progressed CFPS expression of chrysanthemol diphosphate synthase (CcCPPase) from *Chrysanthemum cinerariaefolium* (**Supplementary Figure S4**) that catalyses an irregular, non-head-to-tail 1-2 linkage between molecules of dimethylallyl diphosphate (DMAPP) and also hydrolyses the diphosphate moiety to produce chrysanthemol which contains a cyclopropane ring and is precursor to pyrethrins, a widely used class of plant pesticides (73).

We then tested expression of plant transcription factor (TF) proteins, which are of great interest to plant synthetic biologists aiming to engineer complex gene regulatory networks known to control important agricultural traits or to link the synthetic circuits to endogenous processes. Most plants contain large TF families and many of their cognate DNA sequences remain unknown. We selected two TGA-family TFs from *Arabidopsis thaliana* (TGA) and three TFs from *Triticum aestivum* (bread wheat). All five transcription factors were expressed using *E. coli* CFPS with five different N-terminal configurations (**Figure 3B**). The NET and NET-6xHis tags were the most consistent across multiple TFs, but the S-tag proved remarkably helpful in expressing TGA2. To further assess cell-free expression of transcription factors from wheat, we wanted to compare *E. coli* CFPS with expression using TNT SP6 High-Yield Wheat Germ Protein Expression System (Promega). As a first test, we first assembled GFP into three different plasmid acceptors containing different combinations of terminator and antibiotic resistance sequences and found that folA terminator from the pEU vector outperformed the T7 terminator (**Supplementary Figure S5**). We then cloned the three LUX protein coding sequences (along with LHYA and NAMA1) into the optimal wheat germ acceptor and measured soluble and total protein expression using the HiBiT assay. While both systems produced folded protein, the wheat germ kit generally produced higher levels of soluble protein with the exception of NAMA1, which only expressed in the *E. coli* system (**Figure 3C**).

### 3.4 Cell-free expressed proteins are functionally active

Having identified conditions from which we could obtain good yield of several proteins, we wanted to test if the proteins obtained were functionally active and whether assays could be conducted without extensive purification protocols. To test UGT enzymatic activity, we expressed UGT73C5 in 15μL reactions and incubated the reactions with UDP-glucose along with geraniol or *cis-trans*-nepetalactol. Reactions were quenched at one hour (**Supplementary Figure S6**) and then analysed by LC-MS, which monitored possible glucoside-substrate adducts (**Supplementary Figures S7-S8**). Cell-free reactions expressing UGT73C5 in three different tag configurations produced glucosides of both substrates with peaks matching the retention time of glucosides produced by purified UGT73C5 and not present in the GFP negative control (**Figure 4A**). The cell-free reactions, including GFP negative control, contain additional peaks likely derived from unspecified compounds in the *E. coli* lysate. We were also able to detect synthesis of chrysanthemol by GC-MS after incubating cell-free reactions expressing CcCPPase with DMAPP (**Figure 4B**), obtaining the expected fragmentation pattern (**Supplementary Figure S9**).

**Figure 4.**
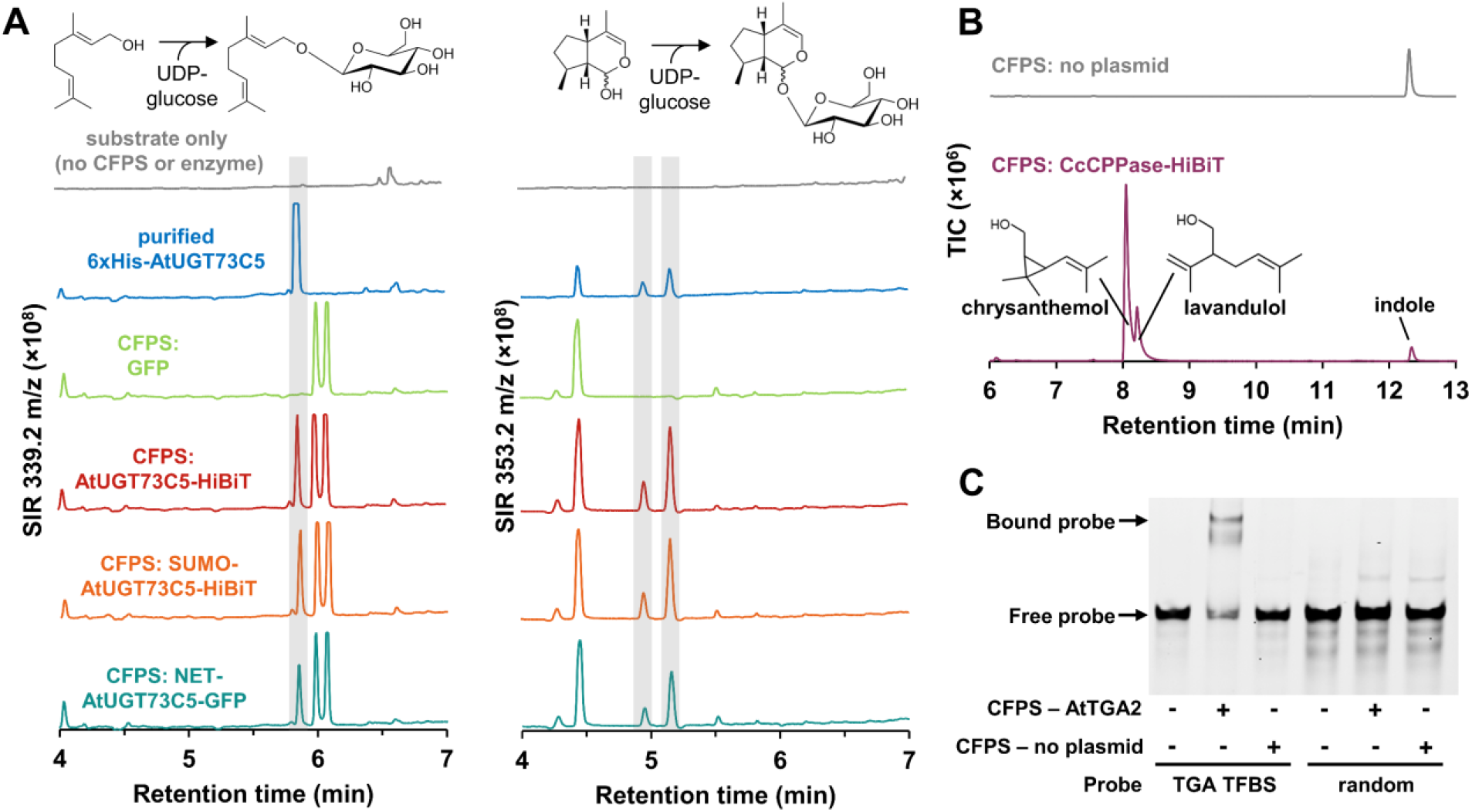
Plant proteins expressing using *E. coli* cell-free protein synthesis are functionally active. (A) UDP-glycosyltransferase UGT73C5 from *Arabidopsis thaliana* can glycosylate geraniol or nepetalactol when purified (blue trace) or when expressed using CFPS (red, orange, teal trace) (B) chrysanthemol diphosphate synthase (CcCPPase) from *Chrysanthemum cinerariaefolium* converts substrate DMAPP to chrysanthemol and lavandulol. (C) Cell-free expressed AtTGA2 binds a 60 bp DNA probe containing its cognate binding site but does not impede mobility of a probe with randomised sequence.

To test the binding ability of cell-free expressed protein to DNA, CFPS-derived AtTGA2 was incubated with a DNA sequence probe containing a known binding site from the Cauliflower Mosaic Virus (CaMV) 35s promoter (74) and DNA-protein interactions visualised by electrophoretic mobility shift assay (EMSA). Cell-free expressed TGA2 successfully bound to the probe containing its binding site but not the random control sequence (**Figure 4C**), suggesting that the DNA-binding domain of the cell-free expressed TGA2 was correctly folded and functional.

## Discussion

Biofoundries provide a powerful combination of automation platforms and synthetic biology approaches including the application of engineering principles and computational modelling and analysis to aid the design and evaluation of large datasets. Consequently, they can significantly increase the scale and throughput for experiments testing a given biological problem or question (12-14). Building on earlier work in which we established automation-compliant DNA assembly tools and protocols for the assembly of constructs for engineering plants systems (3, 5, 10), we have developed a toolbox for cell-free protein synthesis compatible with DNA parts used by the plant community **(Figure 1, Table 1)**. This enabled us to obtain useful yields of both enzymes and regulatory proteins, progressing directly to characterisation experiments. The resulting toolbox contains 37 plasmids including acceptors for T7-driven *E. coli* CFPS as well as commercial wheat germ protein expression. While other Golden Gate assembly systems could potentially be used for *E. coli* cell-free expression, they are customised for specific applications (75-77) and are incompatible with DNA parts in the Phytobrick standard. While, for some proteins, optimal expression levels might be obtained by system-specific codon-optimisation, optimal expression is generally unnecessary for rapid characterisation and is outweighed by the ability to reuse large number of parts in prototyping experiments.

Over the past few decades CFPS has been successfully used to express and prototype bacterial proteins including DNA regulatory elements (50, 77-79), metabolic sensors (80, 81), and ribosomal peptides (82). It has also been applied to metabolic pathways from a range of organisms, mainly to prototype biosynthesis for production in microbial chassis (29, 30, 41, 45, 83, 84). More recently, the advantages of this technology have been applied to enabling the expression of proteins that can be challenging to express, such as cytotoxic proteins and glycoproteins (26, 28, 32, 33, 85, 86). Automated liquid handling has the general advantage of increasing throughput, reducing reaction volumes, and improving repeatability. These advantages have already been applied to assembling the components of CFPS reactions (47-54). In this study we coupled automated nanoscale DNA assembly workflows to CFPS to progress high-throughput experiments that enable the selection of optimal configurations for the expression of multiple plant proteins. We show how software settings, reaction volumes and assembly time are important for consistent cell-free expression and optimise multiple parameters to obtain consistent, repeatable yields from low-volume reactions (**Figure 2, Supplementary Figure S2**). Our workflows were aided by the use of the high-affinity, 11 amino acid HiBiT tag (87), which is able to complex with a 18 kDa engineered NanoLuc polypeptide (88) and luminesce proportional to the level of HiBiT-tagged protein. We found that this worked well as a C-terminal tag but reduced yields when fused to the N-terminus of sfGFP (**Figure 2**), perhaps because the HiBiT amino acid sequence is not optimal for early translational elongation (89). Although sfGFP has been optimised to improve folding (90), the minimal size of the HiBiT tag may be less likely to inhibit protein function and accessibility. However, alternatives may be required for proteins in which the C-terminus needs to be folded inside the protein or must be freely available for activity or post-translational modifications. To assist this, we demonstrate that tags can be removed after synthesis by simply incubating with a parallel reaction expressing TEV protease (**Figure 2C**).

We then applied the twin capabilities of automated high-throughput DNA assembly and low-volume cell-free protein synthesis to rapidly select the tag configurations that resulted in useful levels of expression of different plant proteins. Although we did not expect to find a single configuration that was generally applicable to multiple types of proteins, there was surprisingly little consistency between even closely related proteins. Tags that significantly improved the yields of some family members resulted in little or no expression of others (**Figure 3A**). This demonstrates the utility of automated high-throughput and low-volume screens to select optimal configurations.

Cell-free expression has yet to be widely applied in plant science but has had impact in the development of next-generation sequencing techniques such as DNA affinity purification (DAP) sequencing. In this technique, wheat germ-based cell-free expression systems have been used to produce affinity tagged TFs that are used to capture genomic DNA enabling the identification of TF binding sites (91, 92). However, when applied to genome-scale collections of TFs, insufficient expression was obtained for several hundred proteins (91). We expressed TGA TFs, known to regulate expression of defence related genes (93-97) as well as the CaMV 35s promoter (74, 98-102). We also expressed wheat homologues of TFs that regulate circadian rhythms in Arabidopsis including homeologues of LUX ARRHYTHMO (LUX) from each of the three wheat sub-genomes (103). The *E. coli* lysate system provides substantial cost-benefits and, although better yields were obtained with the wheat-germ system for three TFs of wheat origin, the *E. coli* system proved beneficial two additional TFs (LHYA and NAMA1) (**Figure 3C**). We do not, however, expect success with all protein classes and did not, for example, attempt the synthesis of plant proteins that are widely known to express poorly in non-eukaryotes such as cytochrome P450s and oxidosqualine cyclases, which in their native cells are localised to lipid droplets or embedded into the endoplasmic reticulum (104, 105).

Plants synthesise a diverse array of metabolites that contribute to adaptation to ecological niches, serving as attractants for beneficial organisms and providing defence against biotic and abiotic agents (106). Metabolic profiling has been widely applied to assess this diversity, identifying a number of molecules that have been leveraged for use in industry and medicine. However, the genetic basis of biosynthesis for many molecules remains unknown. With many genomes now sequenced and many more underway, a major challenge is to assign function to sequence, which is particularly challenging for large enzyme families. Sequence similarity allows the classification of enzymes into large and complex superfamilies, but the highly specific reactions, substrates and products that enzymes catalyse remain difficult to predict. For example, there is interest in characterising the substrate specificity of plant glycosyltransferases as it has been observed that some metabolites are over-glycosylated by native enzymes present *N. benthamiana* hindering yield (107-110). A significant advantage of cell-free expression was the ability to progress directly to characterisation without time-consuming cell-disruption and purification protocols. We were able to express and demonstrate the expected activity an Arabidopsis UGT that has previously been leveraged for overproduction of triterpene saponins (111-113).

As these approaches are applied to a wider range and number of diverse proteins, high-throughput, low-cost experimental pipelines will be useful both for directly assigning function to novel sequences but also for the creation of large datasets can train machine learning algorithms to predict the characteristics such as specificity from primary sequence (114, 115). Further, automated reaction assembly may be a helpful approach for adjusting cell-free reaction conditions unique to a given protein of interest (49, 85).

## Supplementary data

Supplementary Data are available at SYNBIO Online

Supplementary Figure S1. Optimisation of tag cleavage by TEV protease

Supplementary Figure S2. Sequential improvement of Echo assembly protocol for consistent cell-free protein synthesis.

Supplementary Figure S3. Correlation of 2 µL vs 15 µL expression of various UDP-glycosyltransferases.

Supplementary Figure S4. Quantification of CcCPPase-HiBiT expression.

Supplementary Figure S5. Comparison of plasmid architecture for expressing sfGFP using the TNT® SP6 High-Yield Wheat Germ Protein Expression System

Supplementary Figure S6. Reaction of purified AtUGT73C5 with 0.5 mM geraniol and 1 mM UDP-glucose

Supplementary Figure S7. Monitoring of six ions for geraniol-glucoside production. Supplementary Figure S8. Monitoring of six ions for cis-trans-nepetalatcol-glucoside production.

Supplementary Figure S9. GC-MS Total Ion Chromatogram of CcCPPase-HiBiT reaction with DMAPP. Supplemental Table S1. Description of cell-free expression plasmids assembled from acceptor plasmids and Level 0 DNA parts (phytobricks).

Supplemental Table S2. Nucleotide sequences of Level 0 DNA parts (phytobricks) encoding plant proteins

## Author Contributions

Q.M.D. and N.J.P. conceived the study. Q.M.D and H.D. developed the acceptor parts. Q.M.D. expressed tagged GFP variants, ran Echo optimisation reactions, and tested UGTs using LC-MS. J.A.C.L assisted with development of automation protocols. Y.C. expressed TGA transcription factors and performed EMSA, K.K. expressed CPPase and performed GC-MS. Q.M.D. and N.J.P. wrote the manuscript. N.J.P obtained funding and provided supervision.

## Acknowledgements

We thank Susan Duncan and Hannah Rees for provision of coding sequences for wheat transcription factors. We also thank Mark Youles for contributing help with plasmid development. pJL1 was a gift from Michael Jewett (Addgene plasmid # 69496; http://n2t.net/addgene:69496; RRID:Addgene_69496). pEU-Gm23 was a gift from Gabor Igloi (Addgene plasmid # 53738; http://n2t.net/addgene:53738; RRID:Addgene_53738)

## Data Availability

Plasmids are available from Addgene (#162281-162317). Nucleotide sequences and corresponding accession numbers are provided in the supplementary data.

## Funding

The authors gratefully acknowledge the support of the Biotechnology and Biological Sciences Research Council (BBSRC) and the Engineering and Physical Sciences Research Council (EPSRC); this research was funded by the BBSRC Core Strategic Programme Grant (Genomes to Food Security) BB/CSP1720/1; the National Capability Grant BBS/E/T/000PR9815 (Earlham Biofoundry); BB/P010490/1 an industrial partnership award with Leaf Expression Systems; BB/R021554; and the BBSRC/EPSRC OpenPlant Synthetic Biology Research Centre Grant (BB/L014130/1). The funders had no role in study design, data collection and analysis, decision to publish, or preparation of the manuscript.

## Conflict of interest statement

None declared

## Notes

### Competing Interest Statement

The authors have declared no competing interest.

