## Supplemental Information for "Biofoundry-assisted expression and characterisation of plant proteins"

#### Supplementary Material

Supplementary Figures S1-9
Supplementary Tables S1-2

**Supplementary Figure S1. Optimisation of tag cleavage by TEV protease** (A) Comparison of cell-free expression levels for two plasmids encoding TEV protease (S219V). Protein concentration was measured by HiBiT; bar chart values represent averages (n=3) and error bars represent 1 standard deviation. (B) Protease cleavage of an MBP-sfGFP fusion protein at varying ratios of TEV protease (pEPQDKN0729) to target protein. The pEPQDKN0729 plasmid contains an N-terminal expression tag (similar to pEPQDKN0329) but has no C-terminal HiBiT tag.

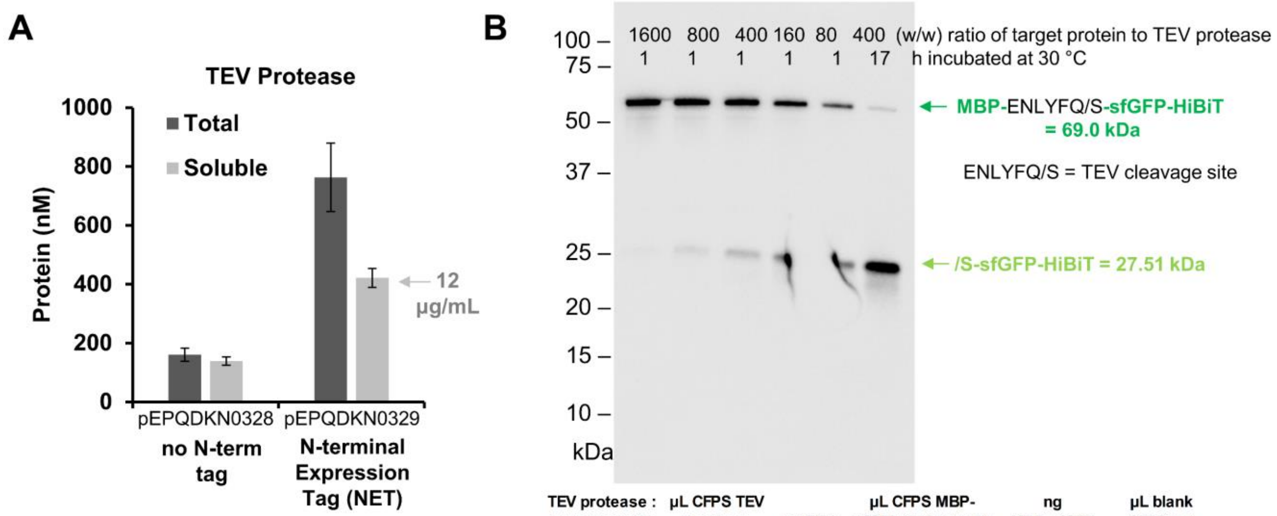


| TEV protease : target protein (w/w) | µL CFPS TEV protease (12 µg/mL) | ng TEV protease | µL CFPS MBP-sfGFP (559 µg/mL) target protein | ng MBP-sfGFP target protein | µL blank CFPS rxn (no plasmid) |
|---|---|---|---|---|---|
| 1:1600 | 0.14 | 1.7 | 5.00 | 2800 | 4.86 |
| 1:800 | 0.28 | 3.5 | 5.00 | 2800 | 4.72 |
| 1:400 | 0.57 | 6.9 | 5.00 | 2800 | 4.43 |
| 1:160 | 1.41 | 17.3 | 5.00 | 2800 | 3.59 |
| 1:80 | 2.83 | 34.6 | 5.00 | 2800 | 2.17 |

**Supplementary Figure S2. Sequential improvement of Echo assembly protocol for consistent cell-free protein synthesis.** All CFPS reactions are expressing pJL1-sfGFP (Addgene #69496) (A) The 2000 nL CFPS reaction consists of 265 nL of plasmid/water plus 1735 nL of a master mix containing the remaining reagents. The high viscosity of the master mix suggested that different sample plate type settings would transfer at different efficiency/consistency. A "droplet" test to transfer from the source plate to a foil-covered destination plate show that 384PP_AQ_GP2 and 384PP_DMSO2 were unable to transfer the master mix. (B) The sample plate type settings that successfully transferred master mix were compared in their ability to assemble CFPS reactions expressing sfGFP. 384PP_AQ_SP2 performed the most consistently, however, there is sufficient volume in the master mix source well to complete only 23 of 24 planned transfers. (C) Reducing the number of destination wells per source well avoided missed transfers, however, the long run time means the reactions assembled last (columns 23 and 24) have lower fluorescence. (D) To reduce the run time, we compared four different reaction volumes. 2000 nL is more consistent than 500 nL and produces the largest amount of protein. Pipetting master mix and *then* plasmid also minimized variability across the plate. (E) To further reduce the transfer time, the software program Echo Plate Reformat was used in place of Echo Cherry Pick to decrease the time taken for well scanning during master mix transfer.

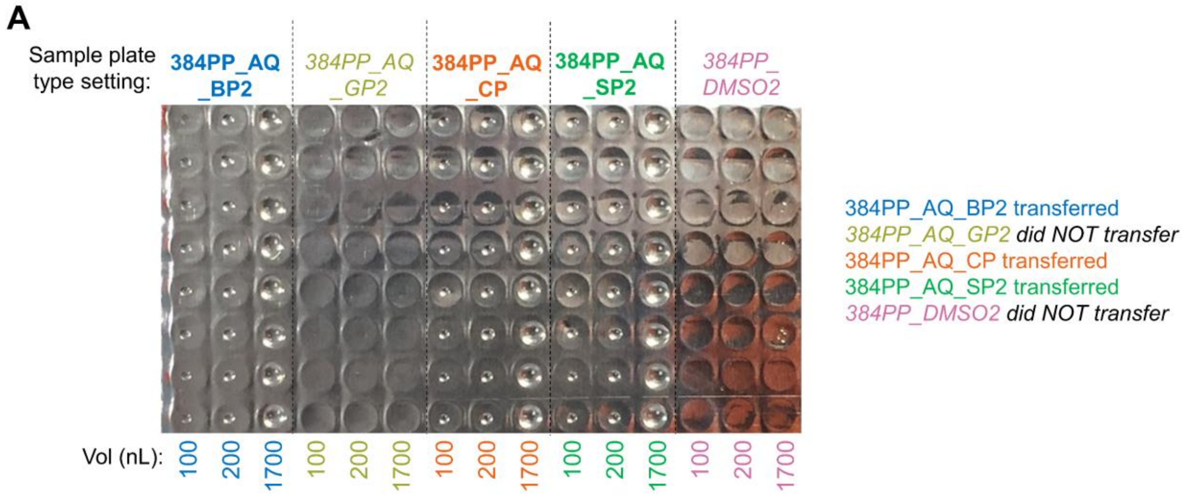


**B**

Sample plate type setting for CFPS master mix: variable (384PP_AQ_BP2, 384PP_AQ_SP2, 384PP_AQ_CP)

Destination wells per master mix source well: 24

Reaction volume: 2000 nL

Assembly order using Echo Cherry pick in single run: Plasmid/water first, CFPS master mix second (53 min total time)

384PP_AQ_BP2 (rows A–H, columns 1–12); 384PP_AQ_CP (rows I–P, columns 1–12)

| | 1 | 2 | 3 | 4 | 5 | 6 | 7 | 8 | 9 | 10 | 11 | 12 |
|---|---|---|---|---|---|---|---|---|---|---|---|---|
| A | 58944 | 26314 | 27 | 59011 | 5238 | 98060 | 53000 | 19099 | 57640 | 58982 | 90492 | 30 |
| B | 55316 | 66896 | 22886 | 69035 | 19712 | 88292 | 73092 | 35139 | 47912 | 80668 | 76445 | 27 |
| C | 56904 | 64259 | 75021 | 65718 | 47222 | 100536 | 70750 | 36221 | 51758 | 63207 | 70950 | 27 |
| D | 55137 | 66250 | 83575 | 60767 | 72703 | 63987 | 64289 | 36704 | 64428 | 53378 | 99570 | 83104 |
| E | 49013 | 81757 | 27 | 65678 | 81399 | 85760 | 66395 | 90597 | 64533 | 63135 | 91258 | 30 |
| F | 68811 | 95904 | 28 | 12677 | 66187 | 65241 | 29034 | 31824 | 89956 | 90867 | 93061 | 28 |
| G | 23231 | 61685 | 30 | 18508 | 98853 | 28 | 20379 | 32405 | 90631 | 85559 | 98965 | 28 |
| H | 47974 | 62309 | 27 | 7541 | 39049 | 28 | 24080 | 70881 | 82486 | 96588 | 83511 | 27 |
| I | 59858 | 59722 | 44093 | 63568 | 61350 | 28 | 62666 | 64243 | 28 | 75606 | 59806 | 27 |
| J | 58054 | 60893 | 54127 | 60751 | 58973 | 27 | 64158 | 68425 | 27 | 58027 | 62222 | 27 |
| K | 56935 | 60465 | 56909 | 53879 | 60460 | 40218 | 54041 | 65023 | 38875 | 47158 | 59279 | 28 |
| L | 54523 | 64638 | 56225 | 53485 | 57673 | 63027 | 51620 | 64222 | 59825 | 54214 | 65125 | 27 |
| M | 55572 | 56098 | 26 | 58502 | 57251 | 29 | 55429 | 57542 | 27 | 58938 | 55830 | 28 |
| N | 51094 | 62681 | 26 | 63274 | 55724 | 28 | 57305 | 63845 | 28 | 69765 | 59932 | 27 |
| O | 54854 | 62367 | 27 | 57352 | 57352 | 27 | 55648 | 55061 | 27 | 59263 | 59878 | 27 |
| P | 56129 | 54429 | 27 | 60074 | 55215 | 27 | 59865 | 55815 | 27 | 62536 | 60532 | 27 |

384PP_AQ_SP2 (rows A–H, columns 13–24); 384PP_AQ_CP (rows I–P, columns 13–24)

| | 13 | 14 | 15 | 16 | 17 | 18 | 19 | 20 | 21 | 22 | 23 | 24 |
|---|---|---|---|---|---|---|---|---|---|---|---|---|
| A | 56594 | 58644 | 54596 | 51751 | 49543 | 46545 | 46303 | 40050 | 42215 | 39559 | 42149 | 39474 |
| B | 62917 | 61739 | 53473 | 66338 | 53451 | 52369 | 53323 | 56443 | 41405 | 45294 | 41536 | 39087 |
| C | 62708 | 58907 | 60203 | 58871 | 56325 | 51321 | 53800 | 50138 | 45984 | 43271 | 42100 | 40853 |
| D | 65230 | 58073 | 55994 | 58901 | 53598 | 55079 | 65994 | 56543 | 50706 | 51258 | 40368 | 49871 |
| E | 60571 | 59675 | 60219 | 58757 | 56614 | 58333 | 56086 | 54610 | 51636 | 50038 | 43343 | 40992 |
| F | 63219 | 66241 | 71226 | 72323 | 57835 | 58290 | 57135 | 61080 | 50809 | 52699 | 40889 | 41897 |
| G | 63092 | 58523 | 38522 | 61057 | 54463 | 48604 | 57022 | 54665 | 35364 | 50179 | 43926 | 36203 |
| H | 60016 | 56125 | 27 | 59612 | 59959 | 28 | 28255 | 61813 | 25 | 59566 | 44623 | 27 |
| I | 71968 | 63655 | 62493 | 53165 | 59628 | 27 | 67104 | 51898 | 52265 | 44026 | 49458 | 49795 |
| J | 59480 | 70343 | 62775 | 27 | 69642 | 27 | 70633 | 45385 | 49921 | 46665 | 51415 | 48854 |
| K | 56607 | 68796 | 59806 | 27 | 77096 | 50220 | 59063 | 43059 | 47898 | 48917 | 49791 | 52834 |
| L | 64219 | 56611 | 72819 | 28 | 72817 | 46747 | 45062 | 40701 | 47731 | 53035 | 51588 | 56662 |
| M | 64581 | 27 | 80571 | 49269 | 64893 | 53893 | 62814 | 54390 | 45953 | 51114 | 42328 | 54199 |
| N | 80616 | 28 | 69818 | 59768 | 58019 | 49051 | 57050 | 52904 | 58320 | 53079 | 53353 | 53649 |
| O | 64669 | 54981 | 63359 | 56863 | 56540 | 46907 | 26 | 53892 | 44736 | 47177 | 51985 | 48308 |
| P | 65441 | 58404 | 61752 | 53594 | 25 | 46183 | 27 | 46243 | 52070 | 44163 | 51389 | 51592 |

**C**

Sample plate type setting for CFPS master mix: 384PP_AQ_SP2

Destination wells per master mix source well: 20 or 21

Reaction volume: 2000 nL

Assembly order using Echo Cherry pick in single run: Plasmid/water first, CFPS master mix second (42 min total time)

| | 1 | 2 | 3 | 4 | 5 | 6 | 7 | 8 | 9 | 10 | 11 | 12 | 13 | 14 | 15 | 16 | 17 | 18 | 19 | 20 | 21 | 22 | 23 | 24 |
|---|---|---|---|---|---|---|---|---|---|---|---|---|---|---|---|---|---|---|---|---|---|---|---|---|
| A | 53148 | 71035 | 69352 | 72650 | 63443 | 66158 | 67962 | 69568 | 64508 | 66835 | 70181 | 67796 | 62197 | 65769 | 64202 | 55656 | 63907 | 67745 | 59772 | 60747 | 55808 | 58202 | 47639 | 38856 |
| B | 66367 | 66951 | 70940 | 70257 | 55516 | 67087 | 76722 | 69943 | 61704 | 69907 | 67937 | 70048 | 65702 | 69534 | 66471 | 52922 | 71516 | 67624 | 65427 | 65932 | 68514 | 59188 | 56142 | 42943 |
| C | 64807 | 68537 | 62777 | 69967 | 64454 | 68671 | 73318 | 69684 | 32850 | 69810 | 66317 | 66556 | 65138 | 66014 | 64069 | 62212 | 66162 | 67919 | 63600 | 65821 | 64162 | 65876 | 55865 | 40766 |
| D | 62110 | 67478 | 69819 | 71840 | 69278 | 70225 | 77662 | 74611 | 58605 | 71845 | 70700 | 65650 | 58968 | 69272 | 70201 | 61370 | 71426 | 65238 | 66571 | 53729 | 67318 | 64735 | 60420 | 42127 |
| E | 73794 | 67679 | 64835 | 68729 | | | | | | | | | | | | 70074 | 65804 | 68164 | 65705 | 64352 | 57283 | 64751 | 58957 | 40774 |
| F | 66532 | 67874 | 70872 | 67884 | | | | | | | | | | | | 72969 | 71415 | 67845 | 63494 | 69733 | 66304 | 66310 | 62676 | 41952 |
| G | 65590 | 59932 | 67371 | 69380 | | | | | | | | | | | | 63455 | 65930 | 71565 | 63930 | 66506 | 57535 | 59468 | 59201 | 43697 |
| H | 69297 | 56439 | 69837 | 69875 | | | | | | | | | | | | 70355 | 68489 | 68460 | 64608 | 70173 | 66350 | 66162 | 56149 | 40488 |
| I | 67949 | 62882 | 65510 | 66991 | | | | | | | | | | | | 70964 | 59733 | 67445 | 62266 | 64055 | 38487 | 60664 | 58776 | 43969 |
| J | 70416 | 67342 | 68176 | 61509 | | | | | | | | | | | | 71398 | 69322 | 67234 | 64660 | 63780 | 28 | 60277 | 57258 | 42404 |
| K | 68145 | 68691 | 65616 | 71264 | | | | | | | | | | | | 69871 | 62651 | 63853 | 65787 | 66055 | 57070 | 61627 | 55276 | 39567 |
| L | 71040 | 68957 | 42025 | 66672 | | | | | | | | | | | | 69077 | 71427 | 65652 | 62077 | 69002 | 60653 | 63186 | 57109 | 41212 |
| M | 64833 | 66619 | 68733 | 66968 | 65248 | 69298 | 69986 | 67633 | 65467 | 73627 | 64211 | 66288 | 65696 | 59895 | 59429 | 66454 | 64662 | 64738 | 64000 | 62456 | 62497 | 57034 | 55252 | 40039 |
| N | 70808 | 74139 | 64950 | 65589 | 70179 | 65992 | 57279 | 67061 | 64646 | 71565 | 63704 | 70005 | 58256 | 57921 | 68593 | 68000 | 69375 | 52604 | 65771 | 69929 | 61071 | 59364 | 50709 | 50307 |
| O | 63202 | 68582 | 65602 | 66640 | 66190 | 65464 | 61890 | 64387 | 63163 | 68573 | 61905 | 64853 | 59834 | 61290 | 60600 | 64372 | 63162 | 48448 | 58214 | 60603 | 63435 | 53352 | 54471 | 55055 |
| P | 68881 | 69742 | 61267 | 65938 | 67969 | 69173 | 68037 | 65813 | 47712 | 69699 | 55059 | 63753 | 61610 | 64374 | 67623 | 62660 | 64825 | 57244 | 57605 | 61807 | 51646 | 43921 | 44355 | 50387 |

| Average | Standard deviation | std dev/ avg |
|---|---|---|
| 63183 | 8526 | 13% |

*Lower fluorescence for wells with master mix added last* →

**D**

Sample plate type setting for CFPS master mix: 384PP_AQ_SP2

Destination wells per master mix source well: variable

Reaction volume: 500, 1000, 1500, 2000 nL

Assembly order in sequence: CFPS master mix *first* using Echo Cherry Pick (19 min), plasmid/water *second* using Echo Cherry Pick (4 min)

| | Average | Standard deviation | std dev/ avg |
|---|---|---|---|
| 2000 nL | 61864 | 2874 | 5% |
| 1000 nL | 40111 | 1639 | 4% |
| 500 nL | 21896 | 2143 | 10% |
| 1500 nL | 51608 | 2448 | 5% |

2000 nL (rows A–H, columns 1–12); 1000 nL (rows A–H, columns 13–24); 500 nL (rows I–P, columns 1–12); 1500 nL (rows I–P, columns 13–24)

| | 1 | 2 | 3 | 4 | 5 | 6 | 7 | 8 | 9 | 10 | 11 | 12 | 13 | 14 | 15 | 16 | 17 | 18 | 19 | 20 | 21 | 22 | 23 | 24 |
|---|---|---|---|---|---|---|---|---|---|---|---|---|---|---|---|---|---|---|---|---|---|---|---|---|
| A | 59269 | 61377 | 64036 | 61864 | 61664 | 63095 | 56817 | 62053 | 57794 | 58300 | 59934 | 63032 | 38008 | 39667 | 39520 | 39320 | 38371 | 36878 | 38486 | 39683 | 38986 | 39403 | 39122 | 38595 |
| B | 62763 | 59202 | 61434 | 59252 | 59609 | 60923 | 62356 | 64109 | 61219 | 60308 | 62926 | 61047 | 40258 | 38298 | 38885 | 39282 | 39699 | 37306 | 37767 | 40094 | 40062 | 39990 | 39564 | 41017 |
| C | 60983 | 61073 | 61657 | 58195 | 61355 | 64396 | 57828 | 61503 | 63300 | 67068 | 59778 | 62518 | 38023 | 39122 | 40149 | 39211 | 38738 | 38531 | 39776 | 40770 | 39701 | 39806 | 37941 | 38197 |
| D | 66479 | 56564 | 61725 | 54919 | 59240 | 57963 | 56915 | 59025 | 60694 | 61103 | 63711 | 61300 | 39404 | 36507 | 40264 | 40620 | 40219 | 39370 | 40581 | 39311 | 42714 | 41394 | 38187 | 40927 |
| E | 61710 | 61900 | 59201 | 61566 | 63058 | 65870 | 59799 | 61220 | 70214 | 64845 | 58014 | 63732 | 38926 | 41415 | 41344 | 40426 | 38651 | 40073 | 39150 | 39996 | 40170 | 39207 | 39018 | 38214 |
| F | 65896 | 61908 | 61881 | 61535 | 64345 | 63594 | 68346 | 61541 | 63579 | 61454 | 64478 | 61884 | 41003 | 40094 | 41803 | 40267 | 42374 | 42278 | 40388 | 41580 | 40673 | 40084 | 38564 | 43374 |
| G | 65795 | 54122 | 60917 | 62694 | 65671 | 65210 | 60331 | 62322 | 62443 | 63094 | 58795 | 58905 | 38523 | 42144 | 41916 | 42932 | 39721 | 44531 | 39355 | 40148 | 40633 | 41364 | 39764 | 38503 |
| H | 66003 | 60830 | 62377 | 61020 | 62277 | 60198 | 63385 | 63617 | 68424 | 64736 | 62114 | 68430 | 42384 | 38374 | 45143 | 43263 | 41692 | 41085 | 42349 | 44020 | 41126 | 41483 | 41529 | 41906 |
| I | 23162 | 23125 | 21043 | 24079 | 21677 | 22631 | 21494 | 22759 | 22324 | 21659 | 22224 | 22915 | 48670 | 53292 | 50768 | 52408 | 49426 | 53842 | 47390 | 46728 | 49397 | 51239 | 47764 | 56711 |
| J | 22618 | 21381 | 23820 | 22113 | 22253 | 22736 | 22764 | 22747 | 23245 | 22192 | 24112 | 23831 | 51695 | 53703 | 52159 | 50290 | 53025 | 53271 | 51479 | 50944 | 52439 | 51491 | 50760 | 60133 |
| K | 22871 | 22709 | 21357 | 24235 | 21669 | 23352 | 22313 | 21823 | 23200 | 22748 | 22109 | 22848 | 51341 | 48964 | 49144 | 53237 | 52945 | 51288 | 50716 | 51052 | 52613 | 52551 | 49266 | 51969 |
| L | 23504 | 21520 | 24600 | 20806 | 22090 | 21524 | 23180 | 22073 | 23398 | 21659 | 24274 | 22682 | 54267 | 53029 | 54718 | 53479 | 54530 | 53142 | 55671 | 52228 | 55599 | 52085 | 51620 | 53914 |
| M | 23546 | 23796 | 20619 | 22042 | 21275 | 20858 | 21629 | 20567 | 23235 | 22455 | 22230 | 20740 | 50880 | 52150 | 51082 | 51456 | 51149 | 52635 | 52809 | 48916 | 52563 | 50594 | 48959 | 46251 |
| N | 23236 | 21297 | 22764 | 21781 | 22001 | 21388 | 16282 | 20859 | 22591 | 21898 | 23750 | 22343 | 53377 | 52555 | 54305 | 55435 | 54680 | 53846 | 53800 | 52466 | 54386 | 51450 | 52738 | 51559 |
| O | 22437 | 21159 | 21711 | 21452 | 22281 | 6287 | 21075 | 20094 | 22194 | 20584 | 21528 | 21147 | 50015 | 48682 | 49915 | 51327 | 50270 | 50069 | 48840 | 45237 | 46819 | 51225 | 48931 | 48237 |
| P | 22711 | 21342 | 22568 | 21023 | 21253 | 20727 | 14343 | 20846 | 21508 | 20074 | 22695 | 22328 | 55569 | 49120 | 53406 | 49809 | 52401 | 48709 | 52283 | 52789 | 47016 | 50429 | 51741 | 53096 |

**E**

Sample plate type setting for CFPS master mix: 384PP_AQ_SP2

Destination wells per master mix source well: 20 or 21

Reaction volume: 2000 nL

Assembly order in sequence : CFPS master mix *first* using Echo Plate Reformat (18 min), plasmid/water *second* using Echo Cherry Pick (7 min)

| | 1 | 2 | 3 | 4 | 5 | 6 | 7 | 8 | 9 | 10 | 11 | 12 | 13 | 14 | 15 | 16 | 17 | 18 | 19 | 20 | 21 | 22 | 23 | 24 |
|---|---|---|---|---|---|---|---|---|---|---|---|---|---|---|---|---|---|---|---|---|---|---|---|---|
| A | 58135 | 55662 | 59342 | 59409 | 61627 | 63875 | 62682 | 57120 | 60549 | 61440 | 63034 | 58848 | 62176 | 59200 | 66355 | 59891 | 61511 | 62395 | 60107 | 62666 | 58323 | 65433 | 61556 | 58198 |
| B | 33470 | 56097 | 62717 | 61844 | 59836 | 57013 | 61253 | 57835 | 60937 | 62867 | 60026 | 62173 | 57298 | 59496 | 59242 | 60749 | 59983 | 60565 | 58952 | 56642 | 64092 | 54669 | 57793 | 61010 |
| C | 50325 | 62450 | 60018 | 61107 | 61231 | 66045 | 63806 | 58925 | 64318 | 63461 | 64515 | 61832 | 63492 | 62629 | 62814 | 59497 | 62826 | 63281 | 62646 | 65083 | 64727 | 62700 | 65331 | 60873 |
| D | 62029 | 64641 | 62886 | 64686 | 61241 | 62277 | 64116 | 58924 | 61666 | 63914 | 63120 | 62634 | 61740 | 60227 | 62617 | 62390 | 61466 | 60022 | 60850 | 59314 | 63615 | 59123 | 58664 | 60273 |
| E | 50683 | 66872 | 61258 | 64703 | | | | | | | | | | 65279 | 65132 | 59772 | 64289 | 70980 | 62529 | 59070 | 60998 | 61440 | 59570 | 61561 |
| F | 56935 | 60378 | 72331 | 65253 | | | | | | | | | | 64554 | 69249 | 65554 | 66458 | 65152 | 62396 | 59330 | 66389 | 62201 | 59963 | 62701 |
| G | 55784 | 60281 | 58599 | 59576 | | | | | | | | | | 64274 | 65918 | 61348 | 62566 | 62501 | 61117 | 61145 | 62541 | 60409 | 60386 | 67244 |
| H | 66777 | 59013 | 65048 | 63333 | | | | | | | | | | 65246 | 62844 | 65618 | 57448 | 59154 | 56768 | 60745 | 63502 | 56233 | 60015 | 61803 |
| I | 61707 | 57451 | 57388 | 59761 | | | | | | | | | 4089 | 63963 | 64827 | 62546 | 57950 | 62585 | 53994 | 62146 | 62588 | 58626 | 59946 | 59586 |
| J | 61335 | 64195 | 68989 | 63497 | | | | | | | | | 1979 | 66603 | 59196 | 60155 | 51515 | 56044 | 55647 | 58189 | 67035 | 62137 | 58779 | 67838 |
| K | 61708 | 66750 | 60636 | 61253 | | | | | | | | | 2613 | 59854 | 57721 | 55448 | 55552 | 54196 | 57583 | 54947 | 59894 | 59659 | 57336 | 58232 |
| L | 57409 | 59766 | 68967 | 63133 | | | | | | | | | 68559 | 66455 | 56935 | 59975 | 54683 | 54102 | 54776 | 57210 | 65294 | 55824 | 52811 | 59090 |
| M | 60610 | 62489 | 57394 | 63004 | 60604 | 61265 | 61516 | 60245 | 58774 | 61756 | 52311 | 56373 | 62289 | 57827 | 56080 | 52152 | 56156 | 54941 | 56089 | 58819 | 60730 | 63885 | 60413 | 58176 |
| N | 51910 | 56106 | 64889 | 62087 | 61273 | 60388 | 59428 | 49879 | 52325 | 55963 | 54235 | 56334 | 55298 | 62727 | 54252 | 54673 | 52878 | 52961 | 54151 | 54183 | 54123 | 51184 | 55627 | 53903 |
| O | 52424 | 66070 | 59936 | 55169 | 61333 | 57547 | 58177 | 56266 | 53297 | 52949 | 55108 | 54741 | 57359 | 52147 | 55695 | 55036 | 53796 | 51679 | 54746 | 57135 | 54490 | 55177 | 55809 | 67795 |
| P | 54639 | 57226 | 61064 | 59816 | 57342 | 61469 | 58204 | 52512 | 55187 | 56880 | 52252 | 60497 | 58193 | 55586 | 45107 | 55505 | 48699 | 50592 | 54309 | 52497 | 55403 | 57609 | 55806 | 66977 |

| Average | Standard deviation | std dev/ avg |
|---|---|---|
| 59763 | 4484 | 8% |

**Supplementary Figure S3. Correlation of 2 µL vs 15 µL expression of various UDP-glycosyltransferases**. (A-B) A subset of reactions from Figure 3A were assembled manually using handheld pipettes (15 µL reactions) or automated using an Echo 550 acoustic liquid handler (2 µL reactions). Total protein was measured by HiBiT. Values represent averages (n=3) and error bars represent 1 standard deviation.

**A**

| | | AtUGT73C5 | NbUGT73A24 | NbUGT73A25 | NbUGT74T6 | NbUGT74N4 | NbUGT76A4 | NbUGT85A73 | NbUGT85A104 | NbUGT94E7 | NbUGT709J6 | NbUGT71AT3 |
|---|---|---|---|---|---|---|---|---|---|---|---|---|
| **Echo 2 µL** | no tag | 427 | | | | | | | | | | 27 |
| | NET | 251 | | | | | | | | 75 | | |
| | S-tag-TEV | 100 | | | 146 | 26 | | 49 | | | 58 | |
| | NET-SUMO-TEV | 469 | 235 | 159 | | | 70 | | 6 | | | |
| **Manual 15 µL** | no tag | 459 | | | | | | | | | | 33 |
| | NET | 288 | | | | | | | | 136 | | |
| | S-tag-TEV | 176 | | | 289 | 39 | | 106 | | | 155 | |
| | NET-SUMO-TEV | 473 | 222 | 212 | | | 212 | | 12 | | | |

**B**

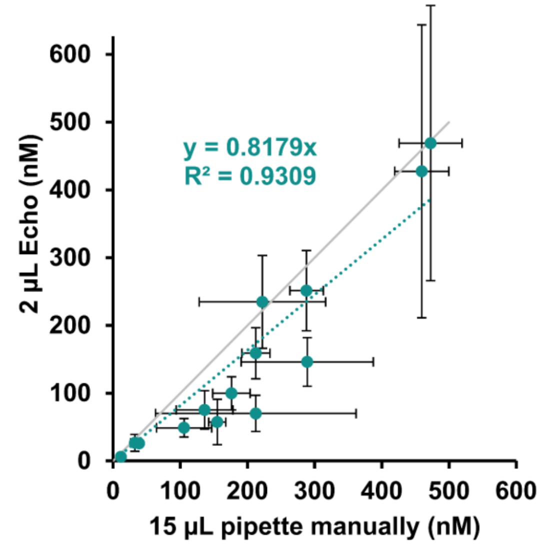

**Supplementary Figure S4. Quantification of CcCPPase-HiBiT expression.** Protein concentration was measured by HiBiT; bar chart values represent averages (n=3) and error bars represent 1 standard deviation.

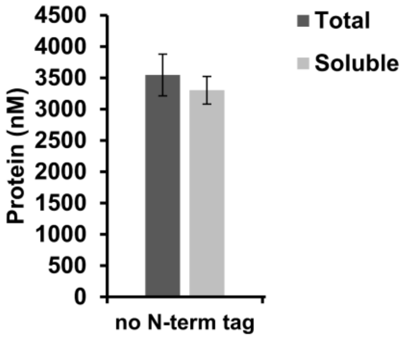

**Supplementary Figure S5. Comparison of plasmid architecture for expressing sfGFP using the TNT® SP6 High-Yield Wheat Germ Protein Expression System** L3260/L3261 (Promega) at 2 µL (A) and 15 µL (B) reaction volumes. The pEU expression plasmid contains a 331 fragment from the *E. coli* MG1655 genome which is positioned 3' of the gene of interest coding region. This genome fragment includes the terminator of the MG1655 folA and apaH coding regions. Using the "folA terminator" (architecture 1 or 2) produced more protein compared to using the T7 terminator (architecture 3). Values in panel A represent averages (n=4) and error bars represent 1 standard deviation.

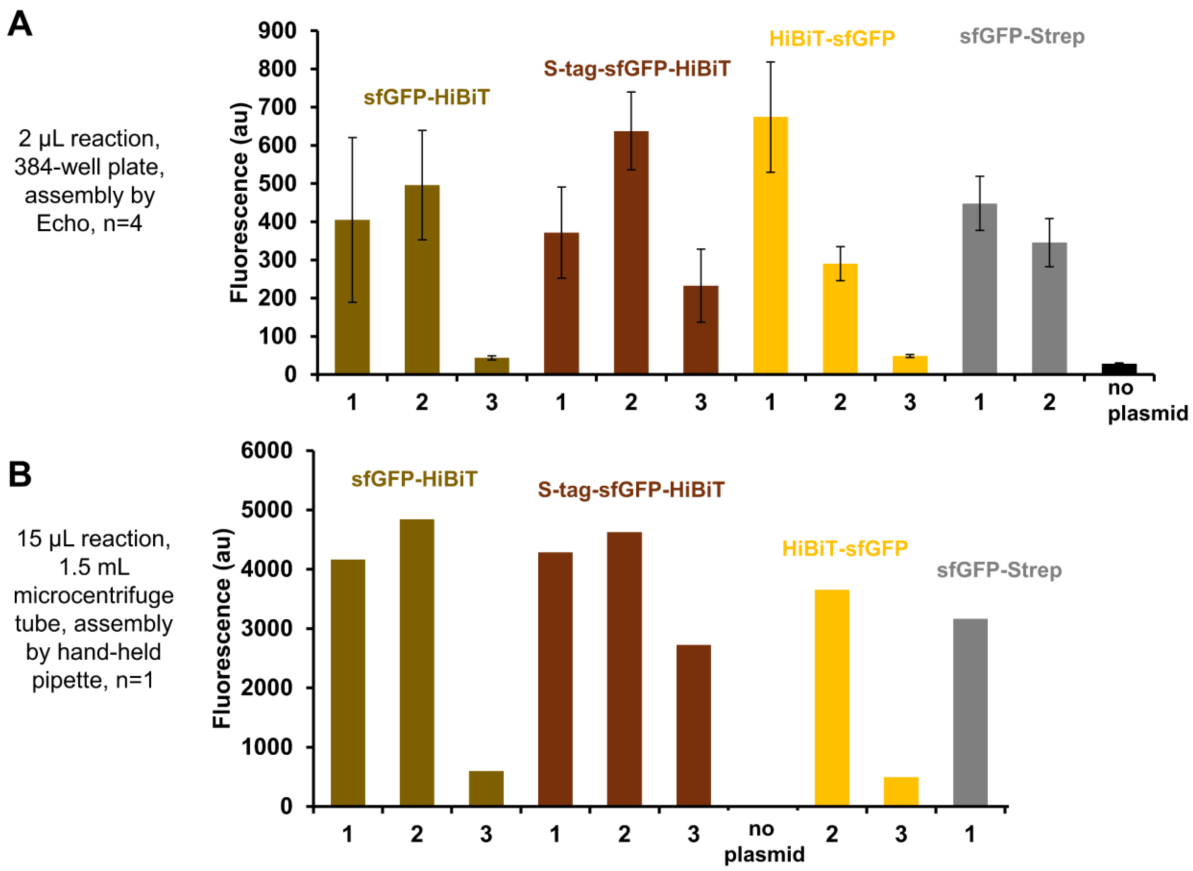


**1: pEU-based**, SP6 promoter, EO1 enhancer, folA terminator, carbR, pUC ori (pEPQD0CB0026, pEPQD0CB0246
**2: pEU/pJL1 hybrid**, SP6 promoter, EO1 enhancer, folA terminator, kanR, pUC ori (pEPQD0KN0284, pEPQD0KN0285)
**3: pJL1-based**, SP6 promoter, EO1 enhancer, T7 terminator, kanR, pUC ori (pEPQD0KN0282, pEPQD0KN0283)

**Supplementary Figure S6. Measurement of geraniol glucoside over time.** The reaction of purified AtUGT73C5 with 0.5 mM geraniol and 1 mM UDP-glucose is mostly complete after one hour.

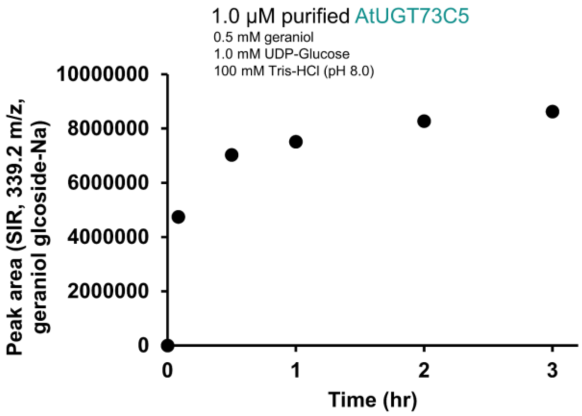

**Supplementary Figure S7. Monitoring of six ions for geraniol-glucoside production.** (A) The sodium (Na) adduct produces the strongest signal. (B) Geraniol-glucoside appears at a retention time of 5.87 minutes in the presence of AtUGT73C5

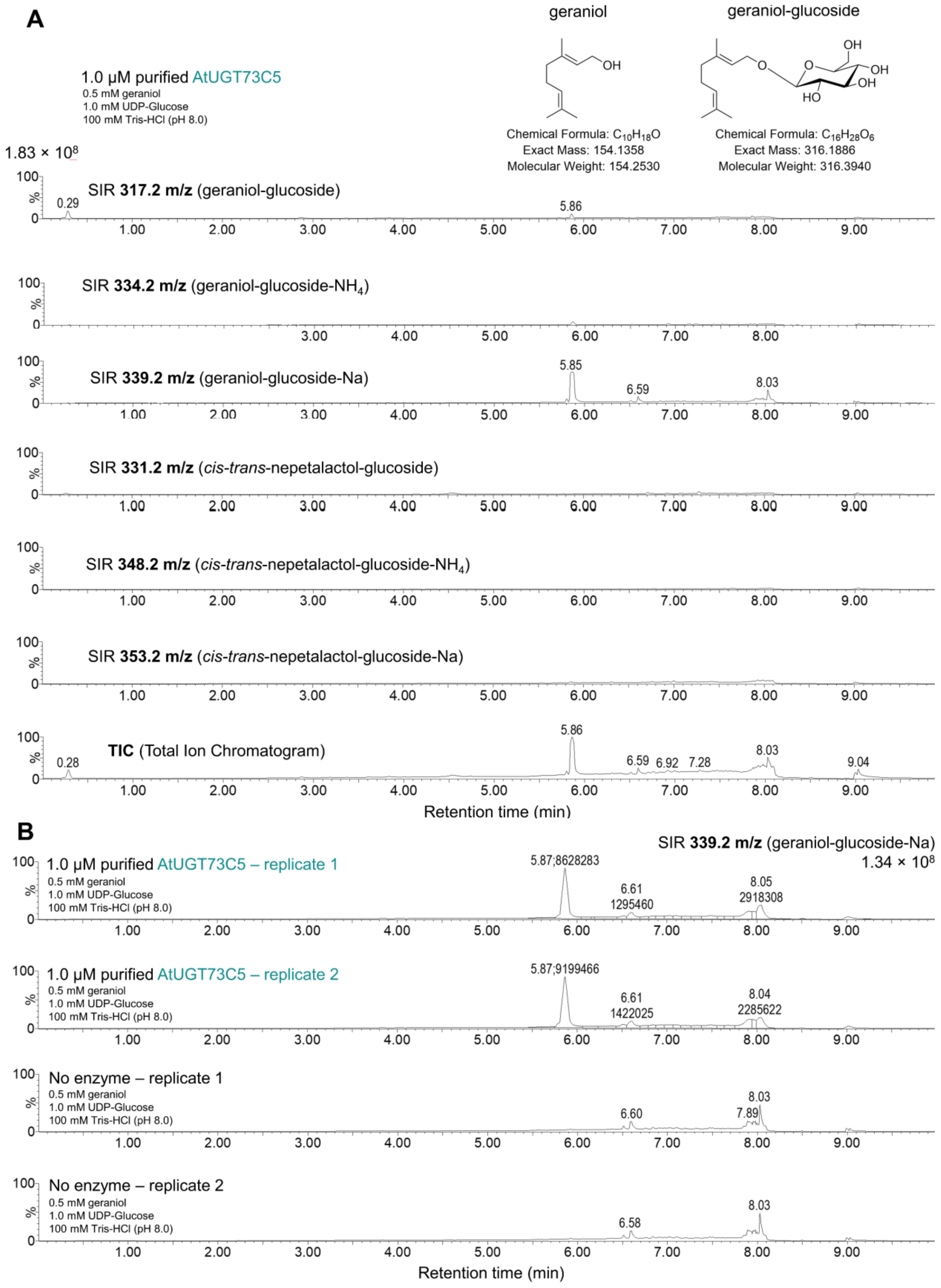

**Supplementary Figure S8. Monitoring of six ions for *cis-trans*-nepetalatcol-glucoside production.** (A) The sodium (Na) adduct producing the strongest signal. (B) *cis-trans*-nepetalctol-glucoside appears at a retention time of 5.15 minutes in the presence of AtUGT73C5

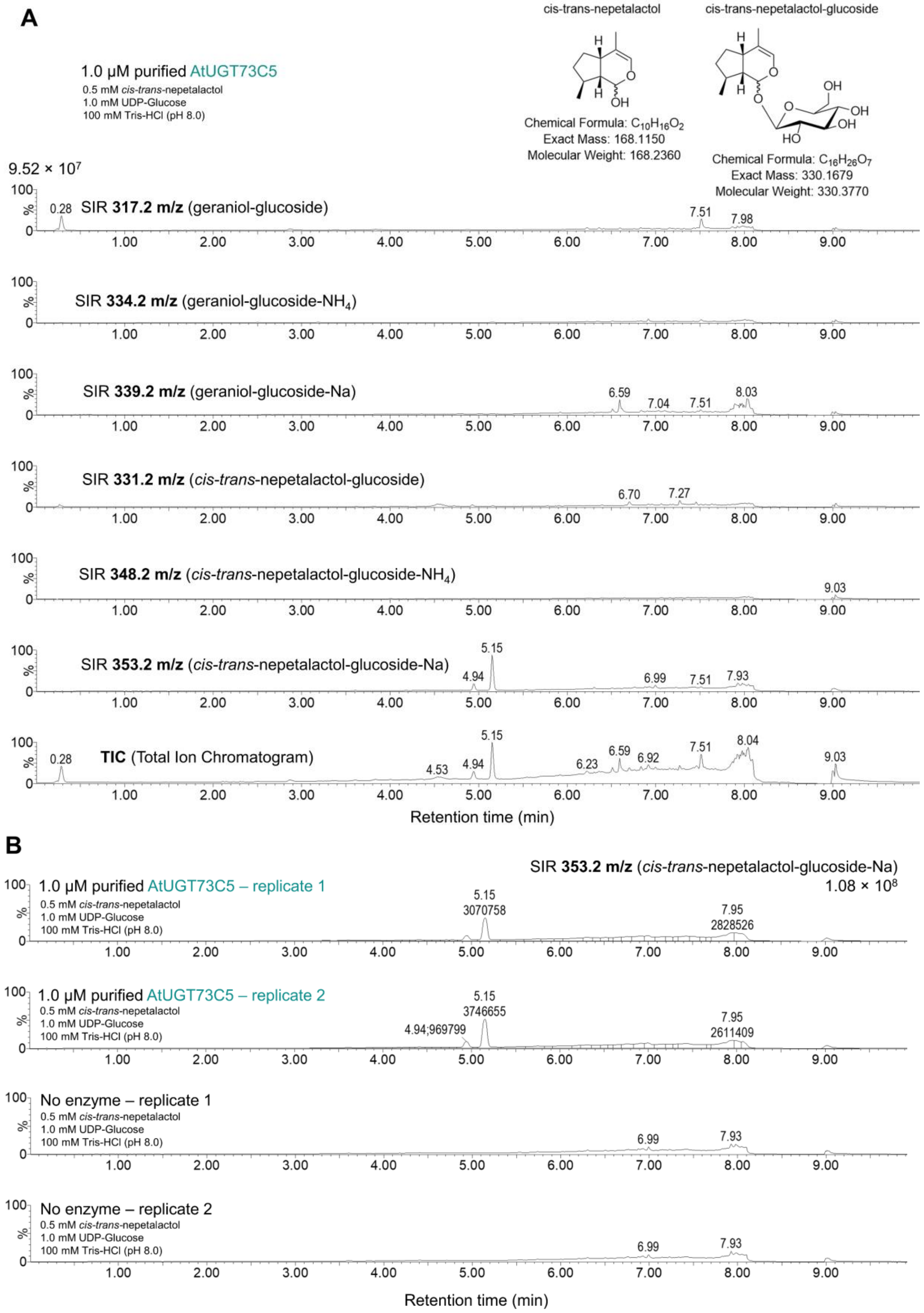

**Supplementary Figure S9**. **GC-MS Total Ion Chromatogram of CcCPPase-HiBiT reaction with DMAPP.** Ion scans of the TIC depicted in **Figure 4B** are compared to the NIST GC Method / Retention Index Library. (A) the peak at 8.08 minutes closely matches chrysanthemol B) the peak at 8.27 minutes closely matches lavandulol

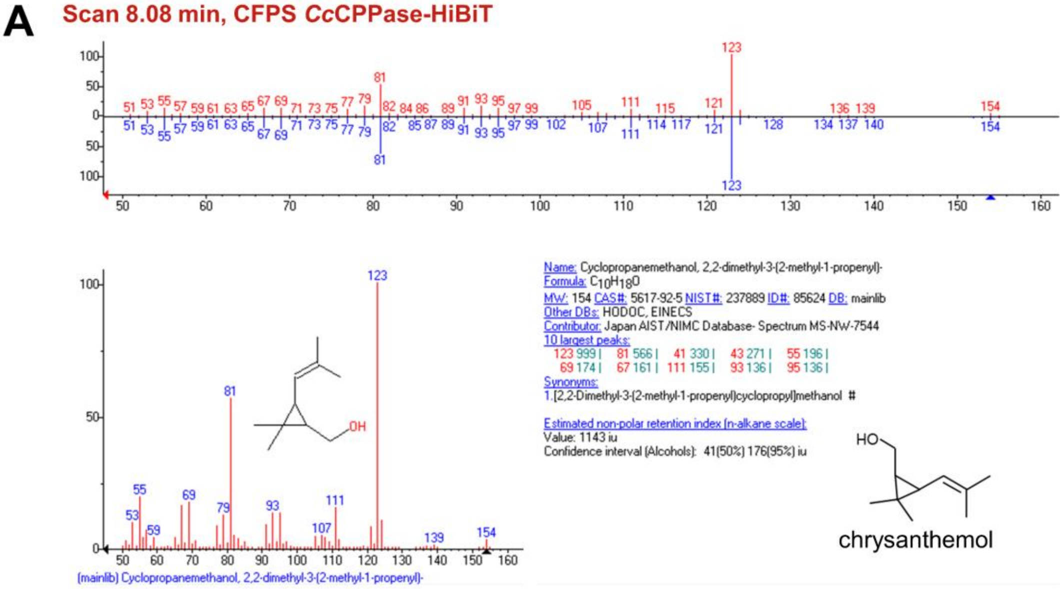


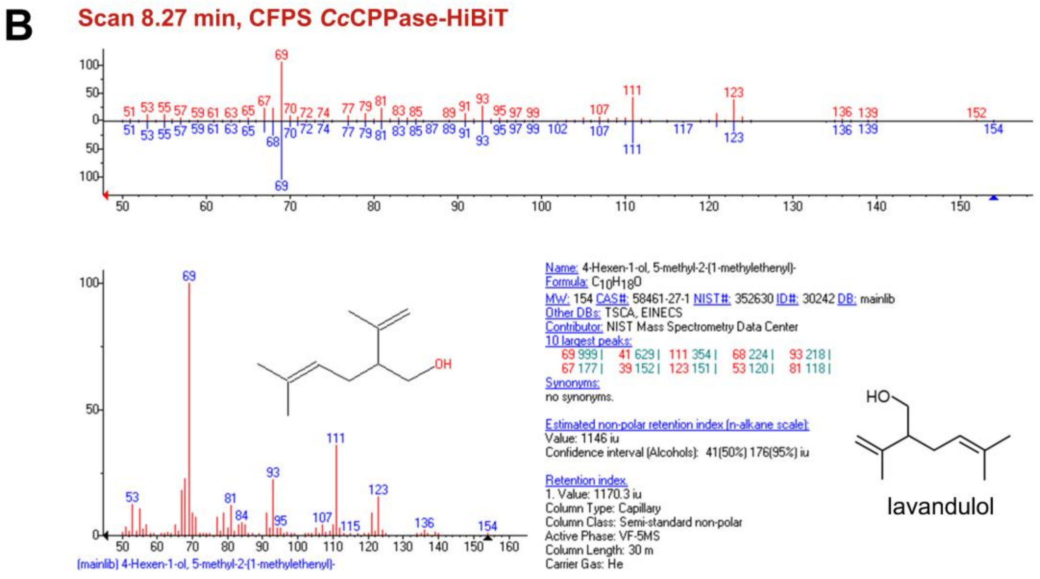

### **Supplemental Table S1**. Description of cell-free expression plasmids assembled from acceptor plasmids and Level 0 DNA parts (phytobricks).

| | Plasmid name | Description | Acceptor | N-term tag | CDS | C-term tag |
|---|---|---|---|---|---|---|
| **Figure 2A and 2C**: All plamids contain T7 promoter and T7 terminator needed for *E. coli* CFPS. **Figure 2B** uses CFPS reactions expressing pEPQDKN0313, pEPQDKN0310, pEPQDKN0314, pEPQDKN0316, pEPQDKN0318, and pEPQDKN0248 along with pEPQDKN0729 (TEV protease) | pJL1-sfGFP | Addgene #69496 | | | | |
| | pEPQDKN0248 | sfGFP-HiBiT | pEPQD0KN0025 | - | pEPQD0CM0296 | pEPYC0CM0134 |
| | pEPQDKN0308 | GST-thrombin-sfGFP-HiBiT | pEPQD0KN0245 | pEPQD0CM0541 | pEPQD0CM0296 | pEPYC0CM0134 |
| | pEPQDKN0309 | HiBiT-GST-thrombin-sfGFP | pEPQD0KN0245 | pEPQD0CM0542 | pEPQD0CM0296 | pEPQD0CM0030 |
| | pEPQDKN0310 | GST-TEV-sfGFP-HiBiT | pEPQD0KN0245 | pEPQD0CM0543 | pEPQD0CM0296 | pEPYC0CM0134 |
| | pEPQDKN0311 | MBP-polyargenine-Factor Xa-sfGFP-HiBiT | pEPQD0KN0245 | pEPQD0CM0544 | pEPQD0CM0296 | pEPYC0CM0134 |
| | pEPQDKN0312 | HiBiT-MBP-polyargenine-Factor Xa-sfGFP | pEPQD0KN0245 | pEPQD0CM0545 | pEPQD0CM0296 | pEPQD0CM0030 |
| | pEPQDKN0313 | MBP-TEV-sfGFP-HiBiT | pEPQD0KN0245 | pEPQD0CM0546 | pEPQD0CM0296 | pEPYC0CM0134 |
| | pEPQDKN0314 | TrxA-TEV-sfGFP-HiBiT | pEPQD0KN0245 | pEPQD0CM0547 | pEPQD0CM0296 | pEPYC0CM0134 |
| | pEPQDKN0315 | HiBiT-TrxA-TEV-sfGFP | pEPQD0KN0245 | pEPQD0CM0548 | pEPQD0CM0296 | pEPQD0CM0030 |
| | pEPQDKN0316 | SUMO-TEV-sfGFP-HiBiT | pEPQD0KN0245 | pEPQD0CM0549 | pEPQD0CM0296 | pEPYC0CM0134 |
| | pEPQDKN0317 | HiBiT-SUMO-TEV-sfGFP | pEPQD0KN0245 | pEPQD0CM0550 | pEPQD0CM0296 | pEPQD0CM0030 |
| | pEPQDKN0318 | S-tag-TEV-sfGFP-HiBiT | pEPQD0KN0245 | pEPQD0CM0281 | pEPQD0CM0296 | pEPYC0CM0134 |
| | pEPQDKN0319 | HiBiT-S-tag-TEV-sfGFP | pEPQD0KN0245 | pEPQD0CM0551 | pEPQD0CM0296 | pEPQD0CM0030 |
| | pEPQDKN0320 | NET-GST-thrombin-sfGFP-HiBiT | pEPQD0KN0244 | pEPQD0CM0541 | pEPQD0CM0296 | pEPYC0CM0134 |
| | pEPQDKN0321 | NET-GST-TEV-sfGFP-HiBiT | pEPQD0KN0244 | pEPQD0CM0543 | pEPQD0CM0296 | pEPYC0CM0134 |
| | pEPQDKN0322 | NET-MBP-polyargenine-Factor Xa-sfGFP-HiBiT | pEPQD0KN0244 | pEPQD0CM0544 | pEPQD0CM0296 | pEPYC0CM0134 |
| | pEPQDKN0323 | NET-MBP-TEV-sfGFP-HiBiT | pEPQD0KN0244 | pEPQD0CM0546 | pEPQD0CM0296 | pEPYC0CM0134 |
| | pEPQDKN0324 | NET-TrxA-TEV-sfGFP-HiBiT | pEPQD0KN0244 | pEPQD0CM0547 | pEPQD0CM0296 | pEPYC0CM0134 |
| | pEPQDKN0325 | NET-SUMO-TEV-sfGFP-HiBiT | pEPQD0KN0244 | pEPQD0CM0549 | pEPQD0CM0296 | pEPYC0CM0134 |
| | pEPQDKN0326 | NET-S-tag-TEV-sfGFP-HiBiT | pEPQD0KN0244 | pEPQD0CM0281 | pEPQD0CM0296 | pEPYC0CM0134 |
| | pEPQDKN0327 | NET-sfGFP-HiBiT | pEPQD0KN0024 | - | pEPQD0CM0296 | pEPYC0CM0134 |
| | pEPQDKN0348 | HiBiT-sfGFP | pEPQD0KN0245 | pEPYC0CM0258 | pEPQD0CM0296 | pEPQD0CM0030 |
| | pEPQDKN0729 | NET-TEVprotease | pEPQD0KN0024 | - | pEPQD0CM0539 | pEPQD0CM0030 |
| **Figure3A**: All plamids contain T7 promoter and T7 terminator needed for *E. coli* CFPS. All UGT coding sequenes were optimized for *E. coli* codon usage | pEPQDKN0437 | AtUGT73C5-HiBiT | pEPQD0KN0025 | - | pEPQD0CM0540 | pEPYC0CM0134 |
| | pEPQDKN0438 | NET-AtUGT73C5-HiBiT | pEPQD0KN0024 | - | pEPQD0CM0540 | pEPYC0CM0134 |
| | pEPQDKN0439 | S-tag-TEV-AtUGT73C5-HiBiT | pEPQD0KN0245 | pEPQD0CM0281 | pEPQD0CM0540 | pEPYC0CM0134 |
| | pEPQDKN0440 | NET-SUMO-TEV-AtUGT73C5-HiBiT | pEPQD0KN0244 | pEPQD0CM0549 | pEPQD0CM0540 | pEPYC0CM0134 |
| | pEPQDKN0441 | TrxA-TEV-AtUGT73C5-HiBiT | pEPQD0KN0245 | pEPQD0CM0547 | pEPQD0CM0540 | pEPYC0CM0134 |
| | pEPQDKN0442 | GST-thrombin-AtUGT73C5-HiBiT | pEPQD0KN0245 | pEPQD0CM0541 | pEPQD0CM0540 | pEPYC0CM0134 |
| | pEPQDKN0443 | GST-TEV-AtUGT73C5-HiBiT | pEPQD0KN0245 | pEPQD0CM0543 | pEPQD0CM0540 | pEPYC0CM0134 |
| | pEPQDKN0444 | MBP-TEV-AtUGT73C5-HiBiT | pEPQD0KN0245 | pEPQD0CM0546 | pEPQD0CM0540 | pEPYC0CM0134 |
| | pEPQDKN0453 | NbUGT74T6-HiBiT | pEPQD0KN0025 | - | pEPQD0CM0552 | pEPYC0CM0134 |
| | pEPQDKN0454 | NET-NbUGT74T6-HiBiT | pEPQD0KN0024 | - | pEPQD0CM0552 | pEPYC0CM0134 |
| | pEPQDKN0455 | S-tag-TEV-NbUGT74T6-HiBiT | pEPQD0KN0245 | pEPQD0CM0281 | pEPQD0CM0552 | pEPYC0CM0134 |
| | pEPQDKN0456 | NET-SUMO-TEV-NbUGT74T6-HiBiT | pEPQD0KN0244 | pEPQD0CM0549 | pEPQD0CM0552 | pEPYC0CM0134 |
| | pEPQDKN0457 | TrxA-TEV-NbUGT74T6-HiBiT | pEPQD0KN0245 | pEPQD0CM0547 | pEPQD0CM0552 | pEPYC0CM0134 |
| | pEPQDKN0458 | GST-thrombin-NbUGT74T6-HiBiT | pEPQD0KN0245 | pEPQD0CM0541 | pEPQD0CM0552 | pEPYC0CM0134 |
| | pEPQDKN0459 | GST-TEV-NbUGT74T6-HiBiT | pEPQD0KN0245 | pEPQD0CM0543 | pEPQD0CM0552 | pEPYC0CM0134 |
| | pEPQDKN0460 | MBP-TEV-NbUGT74T6-HiBiT | pEPQD0KN0245 | pEPQD0CM0546 | pEPQD0CM0552 | pEPYC0CM0134 |
| | pEPQDKN0461 | NbUGT74N4-HiBiT | pEPQD0KN0025 | - | pEPQD0CM0553 | pEPYC0CM0134 |
| | pEPQDKN0462 | NET-NbUGT74N4-HiBiT | pEPQD0KN0024 | - | pEPQD0CM0553 | pEPYC0CM0134 |
| | pEPQDKN0463 | S-tag-TEV-NbUGT74N4-HiBiT | pEPQD0KN0245 | pEPQD0CM0281 | pEPQD0CM0553 | pEPYC0CM0134 |
| | pEPQDKN0464 | NET-SUMO-TEV-NbUGT74N4-HiBiT | pEPQD0KN0244 | pEPQD0CM0549 | pEPQD0CM0553 | pEPYC0CM0134 |
| | pEPQDKN0465 | TrxA-TEV-NbUGT74N4-HiBiT | pEPQD0KN0245 | pEPQD0CM0547 | pEPQD0CM0553 | pEPYC0CM0134 |
| | pEPQDKN0466 | GST-thrombin-NbUGT74N4-HiBiT | pEPQD0KN0245 | pEPQD0CM0541 | pEPQD0CM0553 | pEPYC0CM0134 |
| | pEPQDKN0467 | GST-TEV-NbUGT74N4-HiBiT | pEPQD0KN0245 | pEPQD0CM0543 | pEPQD0CM0553 | pEPYC0CM0134 |
| | pEPQDKN0468 | MBP-TEV-NbUGT74N4-HiBiT | pEPQD0KN0245 | pEPQD0CM0546 | pEPQD0CM0553 | pEPYC0CM0134 |
| | pEPQDKN0469 | NbUGT71AT3-HiBiT | pEPQD0KN0025 | - | pEPQD0CM0554 | pEPYC0CM0134 |
| | pEPQDKN0470 | NET-NbUGT71AT3-HiBiT | pEPQD0KN0024 | - | pEPQD0CM0554 | pEPYC0CM0134 |
| | pEPQDKN0471 | S-tag-TEV-NbUGT71AT3-HiBiT | pEPQD0KN0245 | pEPQD0CM0281 | pEPQD0CM0554 | pEPYC0CM0134 |
| | pEPQDKN0472 | NET-SUMO-TEV-NbUGT71AT3-HiBiT | pEPQD0KN0244 | pEPQD0CM0549 | pEPQD0CM0554 | pEPYC0CM0134 |
| | pEPQDKN0473 | TrxA-TEV-NbUGT71AT3-HiBiT | pEPQD0KN0245 | pEPQD0CM0547 | pEPQD0CM0554 | pEPYC0CM0134 |
| | pEPQDKN0474 | GST-thrombin-NbUGT71AT3-HiBiT | pEPQD0KN0245 | pEPQD0CM0541 | pEPQD0CM0554 | pEPYC0CM0134 |
| | pEPQDKN0475 | GST-TEV-NbUGT71AT3-HiBiT | pEPQD0KN0245 | pEPQD0CM0543 | pEPQD0CM0554 | pEPYC0CM0134 |
| | pEPQDKN0476 | MBP-TEV-NbUGT71AT3-HiBiT | pEPQD0KN0245 | pEPQD0CM0546 | pEPQD0CM0554 | pEPYC0CM0134 |
| | pEPQDKN0477 | NbUGT94E7-HiBiT | pEPQD0KN0025 | - | pEPQD0CM0555 | pEPYC0CM0134 |
| | pEPQDKN0478 | NET-NbUGT94E7-HiBiT | pEPQD0KN0024 | - | pEPQD0CM0555 | pEPYC0CM0134 |
| | pEPQDKN0479 | S-tag-TEV-NbUGT94E7-HiBiT | pEPQD0KN0245 | pEPQD0CM0281 | pEPQD0CM0555 | pEPYC0CM0134 |
| | pEPQDKN0480 | NET-SUMO-TEV-NbUGT94E7-HiBiT | pEPQD0KN0244 | pEPQD0CM0549 | pEPQD0CM0555 | pEPYC0CM0134 |
| | pEPQDKN0481 | TrxA-TEV-NbUGT94E7-HiBiT | pEPQD0KN0245 | pEPQD0CM0547 | pEPQD0CM0555 | pEPYC0CM0134 |
| | pEPQDKN0482 | GST-thrombin-NbUGT94E7-HiBiT | pEPQD0KN0245 | pEPQD0CM0541 | pEPQD0CM0555 | pEPYC0CM0134 |
| | pEPQDKN0483 | GST-TEV-NbUGT94E7-HiBiT | pEPQD0KN0245 | pEPQD0CM0543 | pEPQD0CM0555 | pEPYC0CM0134 |
| | pEPQDKN0484 | MBP-TEV-NbUGT94E7-HiBiT | pEPQD0KN0245 | pEPQD0CM0546 | pEPQD0CM0555 | pEPYC0CM0134 |
| | pEPQDKN0485 | NbUGT73A24-HiBiT | pEPQD0KN0025 | - | pEPQD0CM0556 | pEPYC0CM0134 |
| | pEPQDKN0486 | NET-NbUGT73A24-HiBiT | pEPQD0KN0024 | - | pEPQD0CM0556 | pEPYC0CM0134 |
| | pEPQDKN0487 | S-tag-TEV-NbUGT73A24-HiBiT | pEPQD0KN0245 | pEPQD0CM0281 | pEPQD0CM0556 | pEPYC0CM0134 |
| | pEPQDKN0488 | NET-SUMO-TEV-NbUGT73A24-HiBiT | pEPQD0KN0244 | pEPQD0CM0549 | pEPQD0CM0556 | pEPYC0CM0134 |
| | pEPQDKN0489 | TrxA-TEV-NbUGT73A24-HiBiT | pEPQD0KN0245 | pEPQD0CM0547 | pEPQD0CM0556 | pEPYC0CM0134 |
| | pEPQDKN0490 | GST-thrombin-NbUGT73A24-HiBiT | pEPQD0KN0245 | pEPQD0CM0541 | pEPQD0CM0556 | pEPYC0CM0134 |
| | pEPQDKN0491 | GST-TEV-NbUGT73A24-HiBiT | pEPQD0KN0245 | pEPQD0CM0543 | pEPQD0CM0556 | pEPYC0CM0134 |
| | pEPQDKN0492 | MBP-TEV-NbUGT73A24-HiBiT | pEPQD0KN0245 | pEPQD0CM0546 | pEPQD0CM0556 | pEPYC0CM0134 |
| | pEPQDKN0493 | NbUGT73A25-HiBiT | pEPQD0KN0025 | - | pEPQD0CM0557 | pEPYC0CM0134 |
| | pEPQDKN0494 | NET-NbUGT73A25-HiBiT | pEPQD0KN0024 | - | pEPQD0CM0557 | pEPYC0CM0134 |
| | pEPQDKN0495 | S-tag-TEV-NbUGT73A25-HiBiT | pEPQD0KN0245 | pEPQD0CM0281 | pEPQD0CM0557 | pEPYC0CM0134 |
| | pEPQDKN0496 | NET-SUMO-TEV-NbUGT73A25-HiBiT | pEPQD0KN0244 | pEPQD0CM0549 | pEPQD0CM0557 | pEPYC0CM0134 |
| | pEPQDKN0497 | TrxA-TEV-NbUGT73A25-HiBiT | pEPQD0KN0245 | pEPQD0CM0547 | pEPQD0CM0557 | pEPYC0CM0134 |
| | pEPQDKN0498 | GST-thrombin-NbUGT73A25-HiBiT | pEPQD0KN0245 | pEPQD0CM0541 | pEPQD0CM0557 | pEPYC0CM0134 |
| | pEPQDKN0499 | GST-TEV-NbUGT73A25-HiBiT | pEPQD0KN0245 | pEPQD0CM0543 | pEPQD0CM0557 | pEPYC0CM0134 |
| | pEPQDKN0500 | MBP-TEV-NbUGT73A25-HiBiT | pEPQD0KN0245 | pEPQD0CM0546 | pEPQD0CM0557 | pEPYC0CM0134 |

#### *Supplemental Table S1 (continued)*

| | Plasmid name | Description | Acceptor | N-term tag | CDS | C-term tag |
|---|---|---|---|---|---|---|
| **Figure3A (continued)**: All plamids contain T7 promoter and T7 terminator needed for E. coli CFPS. All UGT coding sequenes were optimized for E. coli codon usage | pEPQDKN0501 | NbUGT85A73-HiBiT | pEPQD0KN0025 | - | pEPQD0CM0558 | pEPYC0CM0134 |
| | pEPQDKN0502 | NET-NbUGT85A73-HiBiT | pEPQD0KN0024 | - | pEPQD0CM0558 | pEPYC0CM0134 |
| | pEPQDKN0503 | S-tag-TEV-NbUGT85A73-HiBiT | pEPQD0KN0245 | pEPQD0CM0281 | pEPQD0CM0558 | pEPYC0CM0134 |
| | pEPQDKN0504 | NET-SUMO-TEV-NbUGT85A73-HiBiT | pEPQD0KN0244 | pEPQD0CM0549 | pEPQD0CM0558 | pEPYC0CM0134 |
| | pEPQDKN0505 | TrxA-TEV-NbUGT85A73-HiBiT | pEPQD0KN0245 | pEPQD0CM0547 | pEPQD0CM0558 | pEPYC0CM0134 |
| | pEPQDKN0506 | GST-thrombin-NbUGT85A73-HiBiT | pEPQD0KN0245 | pEPQD0CM0541 | pEPQD0CM0558 | pEPYC0CM0134 |
| | pEPQDKN0507 | GST-TEV-NbUGT85A73-HiBiT | pEPQD0KN0245 | pEPQD0CM0543 | pEPQD0CM0558 | pEPYC0CM0134 |
| | pEPQDKN0508 | MBP-TEV-NbUGT85A73-HiBiT | pEPQD0KN0245 | pEPQD0CM0546 | pEPQD0CM0558 | pEPYC0CM0134 |
| | pEPQDKN0509 | NbUGT709J6-HiBiT | pEPQD0KN0025 | - | pEPQD0CM0559 | pEPYC0CM0134 |
| | pEPQDKN0510 | NET-NbUGT709J6-HiBiT | pEPQD0KN0024 | - | pEPQD0CM0559 | pEPYC0CM0134 |
| | pEPQDKN0511 | S-tag-TEV-NbUGT709J6-HiBiT | pEPQD0KN0245 | pEPQD0CM0281 | pEPQD0CM0559 | pEPYC0CM0134 |
| | pEPQDKN0512 | NET-SUMO-TEV-NbUGT709J6-HiBiT | pEPQD0KN0244 | pEPQD0CM0549 | pEPQD0CM0559 | pEPYC0CM0134 |
| | pEPQDKN0513 | TrxA-TEV-NbUGT709J6-HiBiT | pEPQD0KN0245 | pEPQD0CM0547 | pEPQD0CM0559 | pEPYC0CM0134 |
| | pEPQDKN0514 | GST-thrombin-NbUGT709J6-HiBiT | pEPQD0KN0245 | pEPQD0CM0541 | pEPQD0CM0559 | pEPYC0CM0134 |
| | pEPQDKN0515 | GST-TEV-NbUGT709J6-HiBiT | pEPQD0KN0245 | pEPQD0CM0543 | pEPQD0CM0559 | pEPYC0CM0134 |
| | pEPQDKN0516 | MBP-TEV-NbUGT709J6-HiBiT | pEPQD0KN0245 | pEPQD0CM0546 | pEPQD0CM0559 | pEPYC0CM0134 |
| | pEPQDKN0517 | NbUGT76A4-HiBiT | pEPQD0KN0025 | - | pEPQD0CM0560 | pEPYC0CM0134 |
| | pEPQDKN0518 | NET-NbUGT76A4-HiBiT | pEPQD0KN0024 | - | pEPQD0CM0560 | pEPYC0CM0134 |
| | pEPQDKN0519 | S-tag-TEV-NbUGT76A4-HiBiT | pEPQD0KN0245 | pEPQD0CM0281 | pEPQD0CM0560 | pEPYC0CM0134 |
| | pEPQDKN0520 | NET-SUMO-TEV-NbUGT76A4-HiBiT | pEPQD0KN0244 | pEPQD0CM0549 | pEPQD0CM0560 | pEPYC0CM0134 |
| | pEPQDKN0521 | TrxA-TEV-NbUGT76A4-HiBiT | pEPQD0KN0245 | pEPQD0CM0547 | pEPQD0CM0560 | pEPYC0CM0134 |
| | pEPQDKN0522 | GST-thrombin-NbUGT76A4-HiBiT | pEPQD0KN0245 | pEPQD0CM0541 | pEPQD0CM0560 | pEPYC0CM0134 |
| | pEPQDKN0523 | GST-TEV-NbUGT76A4-HiBiT | pEPQD0KN0245 | pEPQD0CM0543 | pEPQD0CM0560 | pEPYC0CM0134 |
| | pEPQDKN0524 | MBP-TEV-NbUGT76A4-HiBiT | pEPQD0KN0245 | pEPQD0CM0546 | pEPQD0CM0560 | pEPYC0CM0134 |
| | pEPQDKN0525 | NbUGT85A104-HiBiT | pEPQD0KN0025 | - | pEPQD0CM0561 | pEPYC0CM0134 |
| | pEPQDKN0526 | NET-NbUGT85A104-HiBiT | pEPQD0KN0024 | - | pEPQD0CM0561 | pEPYC0CM0134 |
| | pEPQDKN0527 | S-tag-TEV-NbUGT85A104-HiBiT | pEPQD0KN0245 | pEPQD0CM0281 | pEPQD0CM0561 | pEPYC0CM0134 |
| | pEPQDKN0528 | NET-SUMO-TEV-NbUGT85A104-HiBiT | pEPQD0KN0244 | pEPQD0CM0549 | pEPQD0CM0561 | pEPYC0CM0134 |
| | pEPQDKN0529 | TrxA-TEV-NbUGT85A104-HiBiT | pEPQD0KN0245 | pEPQD0CM0547 | pEPQD0CM0561 | pEPYC0CM0134 |
| | pEPQDKN0530 | GST-thrombin-NbUGT85A104-HiBiT | pEPQD0KN0245 | pEPQD0CM0541 | pEPQD0CM0561 | pEPYC0CM0134 |
| | pEPQDKN0531 | GST-TEV-NbUGT85A104-HiBiT | pEPQD0KN0245 | pEPQD0CM0543 | pEPQD0CM0561 | pEPYC0CM0134 |
| | pEPQDKN0532 | MBP-TEV-NbUGT85A104-HiBiT | pEPQD0KN0245 | pEPQD0CM0546 | pEPQD0CM0561 | pEPYC0CM0134 |
| **Figure3B**: All plamids contain T7 promoter and T7 terminator needed for E. coli CFPS. | pEPQDKN0734 | AtTGA1-HiBiT | pEPQD0KN0025 | - | pEPYC0CM0470 | pEPYC0CM0134 |
| | pEPQDKN0735 | NET-AtTGA1-HiBiT | pEPQD0KN0024 | - | pEPYC0CM0470 | pEPYC0CM0134 |
| | pEPQDKN0736 | S-tag-TEV-AtTGA1-HiBiT | pEPQD0KN0245 | pEPQD0CM0281 | pEPYC0CM0470 | pEPYC0CM0134 |
| | pEPQDKN0737 | NET-6xHis-HRV3C-AtTGA1-HiBiT | pEPQD0KN0244 | pEPMY0SP0002 | pEPYC0CM0470 | pEPYC0CM0134 |
| | pEPQDKN0738 | HiBiT-AtTGA1-strep | pEPQD0KN0245 | pEPYC0CM0258 | pEPYC0CM0470 | pEPQD0CM0029 |
| | pEPQDKN0739 | AtTGA2-HiBiT | pEPQD0KN0025 | - | pEPYC0CM0471 | pEPYC0CM0134 |
| | pEPQDKN0740 | NET-AtTGA2-HiBiT | pEPQD0KN0024 | - | pEPYC0CM0471 | pEPYC0CM0134 |
| | pEPQDKN0741 | S-tag-TEV-AtTGA2-HiBiT | pEPQD0KN0245 | pEPQD0CM0281 | pEPYC0CM0471 | pEPYC0CM0134 |
| | pEPQDKN0742 | NET-6xHis-HRV3C-AtTGA2-HiBiT | pEPQD0KN0244 | pEPMY0SP0002 | pEPYC0CM0471 | pEPYC0CM0134 |
| | pEPQDKN0743 | HiBiT-AtTGA2-strep | pEPQD0KN0245 | pEPYC0CM0258 | pEPYC0CM0471 | pEPQD0CM0029 |
| | pEPQDKN0637 | TaLUXA-HiBiT | pEPQD0KN0025 | - | pSD0KN04 | pEPYC0CM0134 |
| | pEPQDKN0638 | NET-TaLUXA-HiBiT | pEPQD0KN0024 | - | pSD0KN04 | pEPYC0CM0134 |
| | pEPQDKN0639 | S-tag-TEV-TaLUXA-HiBiT | pEPQD0KN0245 | pEPQD0CM0281 | pSD0KN04 | pEPYC0CM0134 |
| | pEPQDKN0640 | NET-6xHis-HRV3C-TaLUXA-HiBiT | pEPQD0KN0244 | pEPMY0SP0002 | pSD0KN04 | pEPYC0CM0134 |
| | pEPQDKN0351 | HiBiT-TaLUXA-strep | pEPQD0KN0245 | pEPYC0CM0258 | pSD0KN04 | pEPQD0CM0029 |
| | pEPQDKN0642 | TaLUXB-HiBiT | pEPQD0KN0025 | - | pSD0KN05 | pEPYC0CM0134 |
| | pEPQDKN0643 | NET-TaLUXB-HiBiT | pEPQD0KN0024 | - | pSD0KN05 | pEPYC0CM0134 |
| | pEPQDKN0644 | S-tag-TEV-TaLUXB-HiBiT | pEPQD0KN0245 | pEPQD0CM0281 | pSD0KN05 | pEPYC0CM0134 |
| | pEPQDKN0645 | NET-6xHis-HRV3C-TaLUXB-HiBiT | pEPQD0KN0244 | pEPMY0SP0002 | pSD0KN05 | pEPYC0CM0134 |
| | pEPQDKN0352 | HiBiT-TaLUXB-strep | pEPQD0KN0245 | pEPYC0CM0258 | pSD0KN05 | pEPQD0CM0029 |
| | pEPQDKN0647 | TaLUXD-HiBiT | pEPQD0KN0025 | - | pSD0KN06 | pEPYC0CM0134 |
| | pEPQDKN0648 | NET-TaLUXD-HiBiT | pEPQD0KN0024 | - | pSD0KN06 | pEPYC0CM0134 |
| | pEPQDKN0649 | S-tag-TEV-TaLUXD-HiBiT | pEPQD0KN0245 | pEPQD0CM0281 | pSD0KN06 | pEPYC0CM0134 |
| | pEPQDKN0650 | NET-6xHis-HRV3C-TaLUXD-HiBiT | pEPQD0KN0244 | pEPMY0SP0002 | pSD0KN06 | pEPYC0CM0134 |
| | pEPQDKN0353 | HiBiT-TaLUXD-strep | pEPQD0KN0245 | pEPYC0CM0258 | pSD0KN06 | pEPQD0CM0029 |
| **Figure 3C** | pEPQDKN0638 | NET-TaLUXA-HiBiT, for *E. coli* CFPS | pEPQD0KN0024 | - | pSD0KN04 | pEPYC0CM0134 |
| | pEPQDKN0641 | TaLUXA-HiBiT, WG acceptor 2 | pEPQD0KN0284 | - | pSD0KN06 | pEPYC0CM0134 |
| | pEPQDKN0643 | NET-TaLUXB-HiBiT, for *E. coli* CFPS | pEPQD0KN0024 | - | pSD0KN05 | pEPYC0CM0134 |
| | pEPQDKN0646 | TaLUXB-HiBiT, WG acceptor 2 | pEPQD0KN0284 | - | pSD0KN05 | pEPYC0CM0134 |
| | pEPQDKN0648 | NET-TaLUXD-HiBiT, for *E. coli* CFPS | pEPQD0KN0024 | - | pSD0KN06 | pEPYC0CM0134 |
| | pEPQDKN0651 | TaLUXC-HiBiT, WG acceptor 2 | pEPQD0KN0284 | - | pSD0KN06 | pEPYC0CM0134 |
| | pEPQDKN0846 | NET-TaLHYA-HiBiT, for *E. coli* CFPS | pEPQD0KN0024 | - | pHR0CM03 | pEPYC0CM0134 |
| | pEPQDKN0848 | TaLYHA-HiBiT, WG acceptor 2 | pEPQD0KN0284 | - | pHR0CM03 | pEPYC0CM0134 |
| | pEPQDKN0842 | NET-TaNAMA1-HiBiT, for *E. coli* CFPS | pEPQD0KN0024 | - | pHR0CM02 | pEPYC0CM0134 |
| | pEPQDKN0844 | TaNAMA1-HiBiT, WG acceptor 2 | pEPQD0KN0284 | - | pHR0CM02 | pEPYC0CM0134 |
| **Figure 4A** | pEPQDCB0093 | pOpin-His-AtUGT73C5-stop | pEPMY1CB0001 | pEPMY0SP002 | pEPQD0CM0265 | pEPQD0CM0030 |
| | pEPQDKN0248 | sfGFP-HiBiT | pEPQD0KN0025 | - | pEPQD0CM0296 | pEPYC0CM0134 |
| | pEPQDKN0437 | AtUGT73C5-HiBiT | pEPQD0KN0025 | - | pEPQD0CM0540 | pEPYC0CM0134 |
| | pEPQDKN0440 | NET-SUMO-TEV-AtUGT73C5-HiBiT | pEPQD0KN0244 | pEPQD0CM0549 | pEPQD0CM0540 | pEPYC0CM0134 |
| | pEPQDKN0092 | NET-AtUGT73C5-GFP | pEPQD0KN0024 | - | pEPQD0CM0265 | pEPQD0CM0027 |
| **Figure 4B** | pEPKK1KN0203 | CcCPPase-HiBiT | pEPQD0KN0025 | - | pEPKK0CM0195 | pEPYC0CM0134 |
| **Figure 4C** | pEPQDKN0742 | NET-6xHis-HRV3C-AtTGA2-HiBiT | pEPQD0KN0244 | pEPMY0SP0002 | pEPYC0CM0471 | pEPYC0CM0134 |
| **Supplementary Figure S1** | pEPQDKN0328 | TEVprotease-HiBiT | pEPQD0KN0025 | - | pEPQD0CM0539 | pEPYC0CM0134 |
| | pEPQDKN0329 | NET-TEVprotease-HiBiT | pEPQD0KN0024 | - | pEPQD0CM0539 | pEPYC0CM0134 |
| **Supplementary Figure S4:** All plasmids contain the SP6 promoter, an E01 translational enhancer, and a terminator suitable for the TNT SP6 High-Yield Wheat Germ Protein Expression System | pEPQDCB0334 | sfGFP-HiBiT, WG acceptor 1 | pEPQD0CB0026 | - | pEPQD0CM0296 | pEPYC0CM0134 |
| | pEPQDKN0332 | sfGFP-HiBiT, WG acceptor 2 | pEPQD0KN0284 | - | pEPQD0CM0296 | pEPYC0CM0134 |
| | pEPQDKN0330 | sfGFP-HiBiT, WG acceptor 3 | pEPQD0KN0282 | - | pEPQD0CM0296 | pEPYC0CM0134 |
| | pEPQDCB0335 | S-tag-sfGFP-HiBiT, WG acceptor 1 | pEPQD0CB0246 | pEPQD0CM0281 | pEPQD0CM0296 | pEPYC0CM0134 |
| | pEPQDKN0333 | S-tag-sfGFP-HiBiT, WG acceptor 2 | pEPQD0KN0285 | pEPQD0CM0281 | pEPQD0CM0296 | pEPYC0CM0134 |
| | pEPQDKN0331 | S-tag-sfGFP-HiBiT, WG acceptor 3 | pEPQD0KN0283 | pEPQD0CM0281 | pEPQD0CM0296 | pEPYC0CM0134 |
| | pEPQDCB0752 | HiBiT-sfGFP, WG acceptor 1 | pEPQD0CB0246 | pEPYC0CM0258 | pEPQD0CM0296 | pEPQD0CM0030 |
| | pEPQDKN0350 | HiBiT-sfGFP, WG acceptor 2 | pEPQD0KN0285 | pEPYC0CM0258 | pEPQD0CM0296 | pEPQD0CM0030 |
| | pEPQDKN0349 | HiBiT-sfGFP, WG acceptor 3 | pEPQD0KN0283 | pEPYC0CM0258 | pEPQD0CM0296 | pEPQD0CM0030 |
| | pEPQDCB0055 | sfGFP-strep, WG acceptor 1 | pEPQD0CB0026 | - | pEPQD0CM0296 | pEPQD0CM0029 |
| | pEPQDKN0759 | sfGFP-strep, WG acceptor 2 | pEPQD0KN0284 | - | pEPQD0CM0296 | pEPQD0CM0029 |

**Supplemental Table S2.** Nucleotide sequences of Level 0 DNA parts (phytobricks) encoding plant proteins

| Plasmid name | Plasmid Description / Comments | DNA sequence of "insert" of the plasmid.<br>BsaI sites are coloured red.<br>Overhangs are coloured blue. |
|---|---|---|
| pEPQD0CM0540 | CDS AtUGT73C5 (AT2G36800.1), codon optimised for *E. coli* | GGTCTCTAATGGTAAGCGAGACTACTAAGTCAAGCCCGTTGCATTTCGTCCT<br>GTTCCCGTTTATGGCCCAGGGTCATATGATCCCGATGGTCGACATTGCCCGT<br>CTGCTGGCACAACGCGGGGTAATTATTACCATCGTTACCACCCCGCATAACG<br>CGGCCCGCTTTAAGAACGTGCTGAATCGCGCGATCGAATCGGGTCTGCCGA<br>TTAATCTGGTACAGGTAAAATTCCCGTACTTGGAAGCGGGCCTGCAAGAGGG<br>CCAGGAAAACATTGACTCCCTGGATACCATGGAACGTATGATTCCGTTCTTCA<br>AGGCAGTGAATTTCTTAGAGGAGCCGGTGCAGAAACTTATCGAGGAAATGAA<br>TCCGCGCCCGAGTTGCTTAATCAGTGACTTCTGCCTGCCGTACACGTCGAAG<br>ATTGCGAAGAAATTTAACATTCCGAAAATTCTGTTTCACGGGATGGGCTGTTT<br>CTGCTTGCTCTGCATGCACGTGCTGCGTAAGAATCGCGAAATTCTCGATAAC<br>CTGAAAAGCGACAAAGAATTATTTACAGTACCAGACTTCCCGGACCGCGTGG<br>AGTTTACCCGCACTCAGGTACCAGTTGAGACTTACGTGCCTGCCGGGGATTG<br>GAAGGACATTTTCGACGGAATGGTGGAAGCCAACGAAACCAGTTACGGAGTC<br>ATTGTGAATAGTTTCCAGGAACTGGAACCGGCCTACGCGAAGGATTATAAAG<br>AAGTGCGTTCTGGAAAGGCTTGGACTATCGGGCCTGTAAGCCTGTGTAATAA<br>AGTTGGTGCGGATAAGGCCGAACGCGGTAATAAGTCTGATATCGACCAGGAC<br>GAATGTCTGAAGTGGCTGGACTCAAAGAAGCACGGAAGCGTTTTATATGTCT<br>GCCTGGGCAGCATTTGCAACTTACCACTGTCGCAGCTGAAAGAACTCGGGCT<br>GGGTCTTGAAGAGAGCCAGCGCCCATTTATCTGGGTGATTCGTGGCTGGGA<br>AAAGTATAAGGAACTGGTAGAATGGTTTAGCGAGTCGGGATTCGAGGACCGC<br>ATTCAGGACCGCGGCCTCTTGATTAAGGGCTGGAGCCCGCAGATGCTGATTC<br>TGAGCCACCCTTCGGTCGGTGGTTTTCTGACCCATTGCGGCTGGAATAGTAC<br>CCTCGAAGGTATTACGGCCGGCTTGCCTTTATTAACTTGGCCCTTATTTGCGG<br>ATCAGTTTTGTAACGAAAAGCTGGTTGTGGAAGTTTTAAAGGCGGGTGTTCG<br>CTCGGGCGTAGAGCAACCGATGAAGTGGGGCGAGGAAGAAAAGATCGGCGT<br>TTTAGTTGACAAGGAAGGTGTTAAGAAAGCGGTCGAGGAGCTCATGGGCGAA<br>TCAGACGACGCGAAGGAACGCCGCCGCCGTGCAAAGGAACTGGGCGACAG<br>CGCACATAAAGCCGTAGAGGAAGGTGGTAGCTCGCACTCAAATATTTCATTT<br>CTTCTGCAGGATATTATGGAGCTCGCCGAGCCGAACAACGGTTCGAGAGACC |
| pEPQD0CM0265 | CDS AtUGT73C5 (AT2G36800.1), not codon optimised, contains a BbsI site | GGTCTCGAATGGTTTCCGAAACAACCAAATCTTCTCCACTTCACTTTGTTCTC<br>TTCCCTTTCATGGCTCAAGGCCACATGATTCCCATGGTTGATATTGCAAGGCT<br>CTTGGCTCAGCGTGGTGTGATCATAACAATTGTCACGACGCCTCACAATGCA<br>GCGAGGTTCAAGAATGTCCTAAACCGTGCCATTGAGTCTGGCTTGCCCATCA<br>ACTTAGTGCAAGTCAAGTTTCCATATCTAGAAGCTGGTTTGCAAGAAGGACAA<br>GAGAATATCGATTCTCTTGACACAATGGAGCGGATGATACCTTTCTTTAAAGC<br>GGTTAACTTTCTCGAAGAACCAGTCCAGAAGCTCATTGAAGAGATGAACCCT<br>CGACCAAGCTGTCTAATTTCTGATTTTTGTTTGCCTTATACAAGCAAAATCGCC<br>AAGAAGTTCAATATCCCAAAGATCCTCTTCCATGGCATGGGTTGCTTTTGTCT<br>TCTGTGTATGCATGTTTTACGCAAGAACCGTGAGATCTTGGACAATTTAAAGT<br>CAGATAAGGAGCTTTTCACTGTTCCTGATTTTCCTGATAGAGTTGAATTCACA<br>AGAACGCAAGTTCCGGTAGAAACATATGTTCCAGCTGGAGACTGGAAAGATA<br>TCTTTGATGGTATGGTAGAAGCGAATGAGACATCTTATGGTGTGATCGTCAAC<br>TCATTTCAAGAGCTCGAGCCTGCTTATGCCAAAGACTACAAGGAGGTAAGGT<br>CCGGTAAAGCATGGACCATTGGACCCGTTTCCTTGTGCAACAAGGTAGGAGC<br>CGACAAAGCAGAGAGGGGAAACAAATCAGACATTGATCAAGATGAGTGCCTT<br>AAATGGCTCGATTCTAAGAAACATGGCTCGGTGCTTTACGTTTGTCTTGGAAG<br>TATCTGTAATCTTCCTTTGTCTCAACTCAAGGAGCTGGGACTAGGCCTAGAGG<br>AATCCCAAAGACCTTTCATTTGGGTCATAAGAGGTTGGGAGAAGTACAAAGA<br>GTTAGTTGAGTGGTTCTCGGAAAGCGGCTTTGAAGATAGAATCCAAGATAGA<br>GGACTTCTCATCAAAGGATGGTCCCCTCAAATGCTTATCCTTTCACATCCATC<br>AGTTGGAGGGTTCCTAACACACTGTGGTTGGAACTCGACTCTTGAGGGGATA<br>ACTGCTGGTCTACCGCTACTTACATGGCCGCTATTCGCAGACCAATTCTGCA<br>ATGAGAAATTGGTCGTTGAGGTACTAAAAGCCGGTGTAAGATCCGGGGTTGA<br>ACAGCCTATGAAATGGGGAGAAGAGGAGAAAATAGGAGTGTTGGTGGATAAA<br>GAAGGAGTGAAGAAGGCAGTGGAAGAATTAATGGGTGAGAGTGATGATGCA<br>AAAGAGAGAAGAAGAAGAGCCAAAGAGCTTGGAGATTCAGCTCACAAGGCTG |

| | | |
|---|---|---|
| | | TGGAAGAAGGAGGCTCTTCTCATTCTAACATCTCTTTCTTGCTACAAGACATA<br>ATGGAACTGGCAGAACCCAATAATGGTTCGTGAGACC<br>**generated by PCR amplifying Arabidopsis thaliana Col1 cDNA using primers AGAGGTCTCTAATGGTTTCCGAAACAACCAA and AGAGGTCTCTCGAACCATTATTGGGTTCTGCCAGTTCC; the resulting PCR fragment was assembled into into pUPD2 (Addgene # 68161) using the restriction enzyme BsmBI* (Sarrion-Perdigones *et al. Plant Physiol.* 162, 1618-1634 (2013)) |
| pEPQD0CM0552 | CDS NbUGT74T6 (GenBank ID MT945322), codon optimised for *E. coli* | GGTCTCTAATGGATCTGCTCAACAATAAGAAATACGTGGCACACATCCTGGC<br>ACTGCCGTACCCTTCTCAGGGGCATATCAATCCGATGCTGCAGTTTTGTAAG<br>CGCCTGGTCAGCAAGTCAGTCAAAACGACCCTGGCGATCACCAATTTTATCA<br>GCCACTCTGTACGCCCCATCTCTATCAATGTGTCGATTGACACTATCTCGGAC<br>GGCTTTGATAAGGGCGGTTATGCCGAGGCAGATAGTATCGTTACTTACCTGG<br>AGCGTTTTAAGAAGATTGGGTCACAGACTCTGGAAGATCTGATTAAGAAGTAT<br>GAAAAGAGCGAATTCCCGATCACCTGCGTAATCTACGACGCGTTTATGCCGT<br>GGGCACTGGATGTTGCTAAGGACCACGGTCTGATCGGCTCATGCTTCTTTAC<br>CCAGGCGTGCTCCGTAAATTACATCTATTACTACGTCCACCATGGCAAGCTG<br>ACGCTGCCGATCTCCAGCCCGCCCGTACGTATCCCTGGCCTGCCTGAACTG<br>GAGTTACGTGACATGCCATCCTTTATCTACGTTCACGGCACCTACCCGGCCT<br>ACTTCGAACTGGTCCTTAACCAATTCATTAACGTTGAAAAGGCGGACTACGTG<br>TTCGTAAATAGCTTTTATAAATTAGAAGCAGAAGTGGTTGATGCCATGAGCAA<br>GGTGATCCCCATGTCAACGATTGGACCTACCCTGCCCAGTCTGTACCTGGAT<br>AACCGCGTAGAGAACGATACCGAGTACTGCTTATCTCTGTATCAGGTTGACG<br>CGAGTACGTGCATCAGTTGGCTTAACACGAAAACCGAGGGCAGCGTAGTGTA<br>CGTCGCATTCGGCAGCATGAGCACCATGGACAACGAGCAGATGGAAGAGAT<br>TGCGTGGGGCCTGAAGGCGACTAACTATTATTTCCTGTGGGTGGTCCGCACC<br>TGCGACGAGGCCAAGATCCCGAAGAACTTTATCGAAGAGACTAGCGAAAAGG<br>GGTTGGTTATTAACTGGAGCCCGCAGTTGCAGATTCTCAGTAACAAAGCCAT<br>CGGAATGTTCTTTAGCCACGGCGGCTGGAACTCGACCACCGAGGCGCTTTC<br>CTTGGGTGTACCCATGGTGGTTATGCCGTTATGGACTGACCAGACCACCAAC<br>GCCAAGCTGGTTCAGGACGTGTGGAGCGTGGGAGTTCGTGTCTCAGTGAAC<br>GAAAAGGGCTTCGCGGGGCGCGAGGAGATCGAAAAGTGCGTGCGCATTGTG<br>ATGGAAGGCGATAAAGGCAAGGAGATGAAGAAGAACGCTTTGAAGTGGAAA<br>GACCTGGCAAAAGAGGCAGTCAACGAGGGCGGTACGTCAGACAAGAACATC<br>GAGGAGTTCGTTTCGAAGTGCACCAAGGGTTCGAGAGACC |
| pEPQD0CM0553 | CDS NbUGT74N4 (GenBank ID MT945323), codon optimised for *E. coli* | GGTCTCTAATGTCTACCACCCATAAGGCGCATTGTCTGATTTTACCGTACCCG<br>GTTCAGGGCCACATTAATCCGATGCTGCAGTTCAGCAAGCGCCTGGAGTCTA<br>AGGGAGTGAAGATTACCATTAGCCCAACCAAGTCATTTCTGAAAACAATGCAG<br>GAGCTGCCCACCTCTGTTTCGATTAAAGCGATCTCGGACGGTTATGATGACG<br>GCGGTATTGATCAGGCCGAGAGCTTTCTGGCATATATTACCCGTTTCAAGGA<br>AGTGGGGAGCGACACGCTCACACAGCTGATCAAGAAGCTGGAGTCATGCGA<br>ATATCCGGTTAACTGTATTGTATACGACCCGTTTCTCCCATGGGCAGTAGAGG<br>TCGCCAAAGACCTGGGCCTGGTGAACGCAGCCTTCTTTACTCAGAACTGCGT<br>GGTTGACAATATCTATTATCACGTGCACAAGGGCGTCTTAAAGCTGCCGCCG<br>ACGCAGGTAGACGGCCAGATTCTGATCCCGGGTTTGTCCTCAACCATCGAAA<br>GCAGCGACGTCCCGTCCTTCGAATCCAGCCCCCAGAGCGACAAGTTGGTCG<br>AGATGCTGGTCAACCAGTTTAGCAACTTAGAAAAGGTCGACTGGGTGTTGAT<br>TAATTCCTTTTACGAACTGGAAAAGGAAGTGATCGACTGGATGGCGAAATTCT<br>ACCCTATTAAAACGATCGGTCCTACCATTCCCTCGATGTATTTGGATAAACGC<br>CTGCCCAACGATAAGGAATACGGTTTGAGCTTATTTAAACCTATGGCGAAGG<br>AATGTCTCAACTGGCTGAACCCGCAGCCGATCTCCAGCGTTGTCTACGTGTC<br>GTTCGGGTCCATGGCGAAGCTGGAAGCGGAACAGATGGAAGAACTGGCGTG<br>GGGATCTAAGAACAGTAATAAGAATTTCCTGTGGGTTGTCCGTAGCACGGAA<br>GAGTCGAAGCTGCCTAAGAATTTCATTGAGGAGCTGAAGTCGGCCTCGGAAA<br>AGAAGGGCTTTGTAGTGAGCTGGTGCCCGCAGCTCCAGGTACTGGAGCACA<br>AGTCAATCGGCTGCTTCCTGACCCACAGCGGTTGGAACAGTACCCTTGAGGC<br>CATCAGCCTCGGCGTTCCTATTGTTACCATGCCGCAGTGGAGTGACCAGCCG<br>ACCAACGCCAAATTAGTTCAAGACGTCTGGGAAATGGGCGTGCGCGCGAAG<br>CAGGACGAGAAGGGGATCGTGCGCCGTGAGATCATCGAGGAGAACATTAAG<br>CTGGTCATGGAAGAGGAAAAGGGCAAGGTGATCCGCGAGAACGCCAAGAAG<br>TGGAAAGAGTTCGCCCGCAACGCCGTCGACGAGGGCGGCTCATCGGATAAG<br>AATATCGAGGAGTTTGTCTCAAGCCTGATCACGATCAGTGGTTCGAGAGACC |
| pEPQD0CM0554 | CDS NbUGT71AT3 (GenBank ID MT945324), | GGTCTCTAATGAACGAATTAATCTTTATCCCTCTGGCGGGGCTCGGCCATCT<br>GGTGAGTGCAATTGAGTTCGCGAAGTTAGTGTTAAACCGCGACGACAAGGAC<br>CTGAGTATTAGCGTCTTAATTATGAAGCTCCCTCTGGATTACGGGGTACAGAA<br>CTTTATCCAATCCCTGAACAGCCAGCCGCGCCTGAAGTTTATTGACATTAGTC |

| | | |
|---|---|---|
| | codon optimised for *E. coli* | TGGATGAAAAGACTAGCAGCACCTTTCTCAACAATCACGAGTCCTTTCTTTAC<br>GACTTTATTGACGGGCATAAGTCCAATGTTCGTGAGTACGTTCAGAACATCCC<br>ACGCCTGGCAGGATTTGTACTGGACATGTTCTGCACGTCAATGATCGATATT<br>GCGAATGAGTTCTCGGTGCCTTCTTACATCTACTTCGCGTCGAACGCTGCGT<br>TCCTGGGACTGTGCTTGCACTTCCAGGCCCTCACGAACGAACAGAACCTGGA<br>TACCTCAAAATATGTTAATACAGACAAAGAGCTCTCCATCCCCTACTTCAAGA<br>ACCCTTGCCCGACAAAGGTGCTGCCAAAACACCTGTTGAACAGCCGCCTGG<br>CGAGCACCCTGTTCTTCGACGGCATCCGCCGCTTCAAGGAAACCAAGGGCA<br>TCATCTTGAATACGTTCTTCGAACTGGAGAGTTTTAGCCTTCAAGCCCTGATG<br>GACTCGGAAATCGTGCCGACTATCTACCCCGTGGGCCCGGTCGTGTCTTTCG<br>CCAAGTCCGGCCACTTCCGCAACAACCCTTCTGAAACGGAGTCCATCATTAA<br>ATGGCTGGACGAACAGCCTGACCTTAGCGTTGTCTTTCTGTGTTTCGGATCTA<br>TGGGCAGCTTCGAAGCCGAACAGATCAAGGAGATTGCGACGGCCTTACAGC<br>ACTGTGGCCACCGCTTTCTGTGGAGCCTGCGCCGCAGCCCGCCCAAAGAGA<br>AGATTGACATTCCGTCTAACTATAATAACCTTGAGGAGATCCTGCCCAAGGAA<br>TTCCTGGAACGCACCAAGGGCATTGGCAAGGTTACGGGCTGGGCTCCGCAG<br>GTTGCGATCCTGAGTCACAAGTCTGTTGGCGGGTTCGTTTCGCACTGCGGCT<br>GGAACAGCATTCTGGAGAGCGTTTACTTCGGCGTACCCATTGCCACCTGGCC<br>GTTGTACGCGGAGCAACAGATGAACGCCTTCCTGCTGGTAAAGGAACTGGA<br>GATCGCGGAAGAGATCCGCATGGACTACTTCGTAGACTTCGTTGGACGCAAC<br>TCTAAGGTGGACATCGTGAGCGCCGAAGAGGTGGAAAGCGCCTTGCAGCGT<br>CTGATGGTTAAGAGCGAGGTCCGTGAAAAGGTAAAGAAGATGAAAGAAAAGG<br>CCCGTGTTGCCATGGAAGATGGCGGTTCGAGTTACCTGAGTCTGGCGTTACT<br>GATCAACGACATTATCAGTAACATCAGTGGTTCGAGAGACC |
| pEPQD0CM0555 | CDS NbUGT94E7 (GenBank ID MT945325), codon optimised for *E. coli* | GGTCTCTAATGGACACCCAAGTCATCGAGTGTGGCAATAGCACGTCCCTGAA<br>AGTGTTGATGTTCCCCTGGCTTGCGTACGGCCACATTAGCCCGTTCCTGAAT<br>CTCGCGAAGAAGCTGGCCGATAAAGGCTTTCTCATCTATCTTTGCTCAACGC<br>CGATTAACCTGAAGTCGGTGGTGAAGAAGATTCCGGAGAAGTATTTCGTGTC<br>TATCCAATTAATCGAGTACCACTTGGCGGAGAGCCCGGAGTTACCGCCGCAC<br>TATCACACCACTAACGGTTTACCCCCACACTTGAACCATGCTTTACGTAAGGC<br>GCTCAAGCTGTCTAAGACCAATTTTAGTAAGATTCTCAAGTCCCTTAAGCCGG<br>ACCTGGTCGTGTACGACATCCTGCAACAGTGGGCCGAGGGATTGGCAAACG<br>AGCAGAATATCCCGGCGGTCAAACTGCTCACAAGCGGCGCGGCCGTGTTCT<br>CTTACTTCTTCAATCTTGTTAAGAAGCCAGAAGTAGAGTTCCCGTTCCCGGCA<br>ATCTACCACAAGAAGATCGAACTCGTGAAGCTTGGGGAGATGATGGCTAAGA<br>GTGCGAAGGAGAAGGAGAGCGACGACGTCGGTCCCTTCACCGAGGGCAACA<br>TGCAGATTATGCTGATGTCCACAAGTCGCATCATCGAAGCCAAGTATATTGAC<br>TACAGCTCAGAGCTGAGTAACTGGAAGGTCATCCCGGTGGGGCCGCCTGTG<br>CAGGACTCGATGGCAAACGGTACCGACGATGTAGAACTGTTCGACTGGCTG<br>GGCAAGAAGGACGAAAACTCTACAGTGTTCGTTAGCTTCGGTAGCGAATACT<br>TCCTGACCAAGGAAGATCGTGAGGAGATTGCATTCGGTCTGGAATTCAGCAA<br>CGTCAACTTTATTTGGGTAGTGCGTTTCCCCAACGGAGAGGAGCAGAACCTG<br>GAGAACGCCCTGCCCCAGGGGTTCCTGGAGCGTATCGGTGAGCGTGGGCG<br>CGTACTGAATAAGTGGGCACCGCAACCGCGTATCTTAAACAATCCAAACATT<br>GGCGGTTTCATTTCGCACTGCGGCTGGAACTCGGTCATGGAGTCCGTCGACT<br>TCGGCGTGCCGATTATTGCCATGCCGATGCACCTCGACCAGCCCGTTAACGC<br>CCGCCTCATGGCCGAGCTGGGTGTGGCAGTCGAAATCGTCCGCGGTGATGA<br>CGGTAAAATCCACCGCGAGGAAATCGCCCAGGTGCTGGAGAACGTAATTGC<br>CGGTAAGATTGGTGAGAACCTGCGTACGAAGGTGAAGGACGTATCGAAGAA<br>GCTTAAGAGTGTTCGCGCCGAGGAAACCGACGTGGTGGCAGAGGAGCTGAT<br>CCATTTCTGCAAAACATCCAACAAATGCAAGGGTTCGAGAGACC |
| pEPQD0CM0556 | CDS NbUGT73A24 (GenBank ID MT945326), codon optimised for *E. coli* | GGTCTCTAATGGGGCAACTGCACATCTTCTTCTTTCCCATGATGGCCCACGG<br>ACATATGATCCCGACCTTAGATATGGCAAAACTTTTCGCATCCCGCGGCGTG<br>AAGGCAACCATTATTACCACCCCGCTGAACGAGAGTGTGTTTAGCAAGGCCA<br>TCCAGCGCAATAAACACCTCGGCATTGAGATTGAGATCCGCCTGATTAAGTTT<br>CCGGCAGTGGAGAATGATCTGCCGGAAGAGTGTGAGCGTCTGGACCAGATT<br>CCGAGCGACGAAAAGCTGCCTAACTTCTTCAAGGCGGTCGCTATGATGCAGG<br>AGCCGCTGGAAAAGTTAATCCAGGAGTGTCGTCCGAACTGCCTCGTGTCAGA<br>CATGTTTCTGCCATGGACGACCGACAGCGCTGCGAAGTTCAATATTCCTCGT<br>ATTGTGTTTCACGGAACGTCCTTCTTCGCCCTGTGTGTCGAAAACTCGGTGC<br>GTTTAAACAAACCGTTTAAGAACGTTTCGAGCGACAGCGAGACATTCGTGGT<br>GCCCAACCTTCCGCATGAGATCAAGCTTACTCGTACTCAAGTTAGTCCCTTTG<br>AACGCTCAGGCGAGGAAACGGCAATGACTCGCATGATTAAGACCGTGCGCG<br>AGAGTGACTCGAAAAGTTACGGTGTGGTCTTTAATTCATTTTACGAACTGGAG<br>ACTGACTACGTGGAGCACTACACGAAAGTCTTGGGCCGTCGCGCCTGGGCC |

| | | |
|---|---|---|
| | | ATCGGTCCACTTTCTATGTGTAATCGTGATATCGTGGACAAGGCGGAGCGCG<br>GTAAGAAGTCTAGCATCGACAAGCATGAATGTCTTAAGTGGCTGGACAGCAA<br>GAAGCCATCCAGTGTGGTCTATATCTGCTTTGGTTCAGTCGCCAACTTTACGG<br>CCAGCCAGTTGCATGAGCTGGCCATGGGCATCGAGGCCTCTGGCCAGGAGT<br>TTATCTGGGTGGTGCGTACCGAGCTTGATAATGAGGACTGGCTGCCGGAAG<br>GGTTTGAGGAGCGTACCAAGGAAAAGGGGCTGATTATTCGCGGCTGGGCGC<br>CGCAGGTGCTTATCCTCGACCATGAGTCCGTCGGCGCGTTCGTAACTCACTG<br>CGGTTGGAACTCTACGCTCGAGGGTGTGAGCGGTGGGGTGCCGATGGTTAC<br>CTGGCCGGTGTTCGCAGAACAGTTCTTTAACGAGAAACTGGTAACCGTAGTG<br>TTAAAGACGGGTGCGGGAGTGGGCTCAATCCAATGGAAACGTAGTGCATCC<br>GAGGGTGTTAAGCGTGAGGCGATTGCAAAAGCCGTTAAACGTGTCATGGTTT<br>CGGAAGAGGCTGACGGGTTTCGCAATCGCGCAAAGGCCTATAAAGAGATGG<br>CGACCAAGGCGATCGAGAAGGGCGGTTCATCGTATACCGGCCTCACAACAC<br>TGCTCGAGGACATCAGCACTTACTCATCGACGGACCACGGTTCGAGAGACC |
| pEPQD0CM0557 | CDS NbUGT73A25 (GenBank ID MT945327), codon optimised for *E. coli* | GGTCTCTAATGGGTCAACTCCATTTCTTCTTCTTCCCGATGATGGCGCAAGG<br>GCATATGATTCCAACGCTGGATATGGCCAAATTGGTGGCATCTCGTGGCGTC<br>AAAGCCACAATTATCACCACCCCATTGAACGAATCAGTGTTTTCGAAAAGCAT<br>CCAGCGTAACAAACATTTGGGGATCGAAATTGAAATTCGCTTGATTAAATTTC<br>CTGCAATTGAGAACGACCTGCCGGAAGAATGCGAGCGTTTGGATCAAATTCC<br>GTCTGACGAGAAATTACCTAATTTCTTCAAAGCCACGGCAATGATGCAGGAAC<br>CGCTGGAACAGCTGATTGAAGAATGCCGTCCAAATTGCCTCGTATCCGACAT<br>GTTCTTGCCTTGGACCACGGACAGTGCAGCCAAATTTAACATTCCGCGTATT<br>GTGTTCCACGGCACTTCGTTCTTTGCTCTCTGTGTTGAAAACAGCGTGCGTCT<br>GAATAAACCCTTTAAGAATGTGTCCAGCGATAGCGAAACCTTTGTTGTACCGA<br>ATTTGCCTCATGAGATTAAACTCACGCGCACGCAAGTTTCTCCGTTTGAACAG<br>AGCGGTGAAGAAACGACTATGGCACGCATGATCAAGACAGTATGGGAATCCG<br>ATTCTCGCTCTTACGGGGTGGTGTTTAACTCATTCTACGAGCTGGAAACGGAT<br>TACGTAGAACATTACACCAAGGTTTTAGGCCGTCGCGCCTGGGCCATTGGCC<br>CTCTTTCGATGTGCAATCGCGACATTGAAGATAAAGCAGAACGCGGGAAGAA<br>ATCAAGCATTGACAAGCACGAATGCTTGAAATGGCTGGATTCGAAGAAACCG<br>TCTAGCGTCGTGTATATTTGCTTCGGCAGCGTGGCAAACTTTACTGCAAGTCA<br>GTTGCATGAACTGGCAATGGGCATTGAAGCGTCGGGCCAAGAATTCATCTGG<br>GTTGTGCGTACTGAGTTGGATAACGAAGATTGGCTGCCCGAAGGCTTCGAGG<br>AGCGTACTCGTGAGAAAGGCCTTATTATCCGCGGGTGGGCCCCGCAGGTGG<br>TGATTTTAGATCACGAATCAGTGGGCGCGTTCGTCACTCATTGCGGATGGAA<br>CTCCACGTTAGAGGGCGTTAGCGGTGGTGTACCTATGGTTACCTGGCCGGTA<br>TTCGCTGAGCAATTCTTCAATGAAAAGTTAGTTACCGAAGTGCTGCGTACCGG<br>TGCGGATGTGGGCTCTATTCAATGGAAACGCTCAGCGTCAGAAGGCGTCAAA<br>CGCGAAGCGATCGCCAAAGCCATTCAGCGTGTCATGGTATCGGAAGAGGCA<br>GAGGGCTTCCGTAACCGTGCAAAAGCATATAAAGAAATGGCTCGCAAAGCAG<br>TAGAAGAGGGCGGCAGCAGTTACACCGGCCTGACCACGCTGCTGGAAGATA<br>TCAGCACCTACAGTAGCACCGGTCATGGTTCGAGAGACC |
| pEPQD0CM0558 | CDS NbUGT85A73 (GenBank ID MT945328), codon optimised for *E. coli* | GGTCTCTAATGGGATCGATTGGCGCAGAACTTACAAAACCTCATGCAGTGTG<br>CATTCCGTACCCGGCACAAGGTCATATCAACCCGATGTTAAAATTAGCCAAAA<br>TTCTGCACCACAAAGGCTTCCATATTACCTTTGTCAACACGGAATTTAACCAT<br>CGTCGCCTTCTGAAGTCCCGCGGCCCGGATAGCTTGAAGGGGCTGAGTAGC<br>TTTCGTTTCGAAACCATTCGTGACGGCCTGCCACCCTGTGAAGCAGATGCCA<br>CGCAGGACATCCCCTCGCTGTGTGAATCGACTACCACAACGTGCCTGGGCC<br>CATTTAAAGATCTGTTAGCCAAATTGAATGATACCAATACTAGCAATGTTCCCC<br>CGGTGAGCTGTATTGTGAGCGACGGCGTGATGTCATTTACACTGGCCGCAG<br>CACAAGAGTTAGGCGTGCCAGAAGTACTGTTTTGGACCACTTCGGCGTGTGG<br>ACTGTTAGGATACATGCACTACTACAAAGTGATTGAAAAGGGCTATGCTCCTC<br>TGAAAGATGCTACGGACTTGACCAATGGATACCTGGAGACTACCCTGGATTT<br>TATTCCAGGAATGAAAGATGTCCGTCTGCGCGATCTTCCGAGCTTTCTGCGT<br>ACCACCAATCCTGATGAGTTTATGATCAAATTTGTTCTCCAGGAAACCCAACG<br>CGCTCGCAAGGCCAGTGCCATCATTCTTAACACGTTCGAAACCTTAGAAGCC<br>GAGGTGCTGGAAACTCTTCGCAATTTGTTACCACCTGTGTATCCTATCGGCC<br>CGCTGCATTTCCTGGTTAAGCATGCAGACGATGAGAACCTGAAGGGTTTACG<br>TAGCAGTCTTTGGAAAGAAGAACCCGAATGTATCCAGTGGCTTGACACAAAG<br>GAACCTAACTCGGTTGTCTACGTGAATTTCGGGAGCATCACTGTTATGACTCC<br>GAACCAGTTAATTGAATTTGCTTGGGGCTTGGCAAACTCCCAGCAGACCTTC<br>CTGTGGATTATTCGTCCGGATATTGTTTCGGGTGATGCCAGCATCTTGCCGC<br>CAGAATTTGTGGAAGAAACCAAGAATCGTGGAATGCTGGCGTCATGGTGTAG<br>CCAGGAAGAAGTGTTGTCGCACCCGGCAATTGGCGGTTTTCTGACGCACAG<br>CGGCTGGAACAGCACCTTGGAATCTATTTCGAGCGGTGTACCGATGATTTGT<br>TGGCCCTTCTTTGCTGAACAACAAACCAATTGTTGGTTTAGCGTGACTAAGTG<br>GGATATCGGAATGGAAATTGACAGTGATGTTAAACGTGATGAGGTTGAGTCT |

| | | |
|---|---|---|
| | | CTTGTACGTGAGCTTATGGTTGGCGAGAAAGGCATTAAAATGAAGAAGAAGG<br>CTATGGAGTGGAAGGAACTGGCGGAAGAGAGCGCGAAAGAACACTCCGGAC<br>TTTCCTATGTCAATATTGAAAAGGTTGTGAACGATATTCTGCTGAGCAGCAAA<br>CATGGTTCGAGAGACC |
| pEPQD0CM0559 | CDS NbUGT709J6 (GenBank ID MT945329), codon optimised for *E. coli* | GGTCTCTAATGCTCAGCATGGACAGCAGCGCATTTGTTCCGCACGTGGCAAT<br>TTTCCCGTTTCCGGCGCAAGGACACGTTAATAGCATGCTGAAGCTGGCGCAA<br>CTCCTGTCGGTGTCTAACTTCCATGTGAGCTTCCTGGTGACGGTGGATACGC<br>ACGATCGTCTGTTAAATCATACTGACGTGCTTTCTCGCTTCGGCAGTGAGTTT<br>CATCTGCAAGAGCTGCCCGTGGGCATCAGCTTGGACGAAATGAACACCCGC<br>GACGGCGTGGCGAAACTGCATGACTCACTGAATACGATCGCCAAGCCGTTC<br>CTGCGCGAATTCCTTGCTGAAAGTCCGGTCACCTGCGTAATCGCAGATGGCA<br>TTCTTTCGATGGCCGCGGACGTGGCAGAAGAGATTAATCTGCCGATTATTTA<br>CTTTCGTACGATTAGCGCATGCGCATTTTGGTCATATTTCAGTATTCCCGAAC<br>TGCTGCAGGCGGCAGAGCTGCCGCTGAAGGAGAATGGCATGGATATCACCC<br>TGACTAAAGTTAAGGGAATGGAAGATTTTCTCCGCGGACGTGACTTACCGTC<br>GTTCTGTCGTGTAAGCGATCTGACCAGTGCCGACTTTCGCTTGCTGTCCAGC<br>GAAACACGTCAGACGCCGCGCGCCCGCGGCTTAATTTTAAATACCTTCGAGG<br>ATTTGGAAGGACCTATCTTGAGTCAGATCCGTACCGTATGCCCGAACGTATAT<br>ACCATCGGCCCTGTGCACGCGCACCTCAAAACCCGCCTGGCTACCAAGTGC<br>ACTTCATCCAACTCCCTGTGGCAGGAAGATGAGAACTGTATTAACTGGCTTGA<br>TACCCACCCGCCGAAGTCGGTGTTGTACGTGTCGTTTGGTTCAATCGCAGGT<br>GTCACACGTGAAGAATTGCTGGAATTCTGGTATGGATTGGTCAACAGCGACC<br>AGAATTTTCTGTGGGTGATGCGTGCCGATCTTATCATCGGTCAGGAAGGCAA<br>GCACGAAATCTTAGAAGAGTTAGAACAGGGCACGAAAGCGCGTGGATACATG<br>GCCGACTGGGTACCTCAGGAGAAGGTTCTGGCGCACACTGCGATCGGCGGG<br>TTTCTGACGCATTGCGGCTGGAACAGCACCTTAGAATCCATTGTCGAGGGTG<br>TGCCGATGATCTGTTGGCCGCGCTTCGCCGACCAGCAGGTTAACTCTCGCTT<br>TATCGGGGAAGTATGGAAAATGGGCTTAGATATTAAGGACACCTGCGATCGT<br>GATATCATCGGTAAGAGTATCCGCGACCTTATGGAGAAACGCCGCGGGGAG<br>TTCTTACAGCGCACGGAACAAATGGCTAGCATGGCACGCCGTACCGTGAACG<br>AGGGCGGCTCTAGCTATATCAACCTGGATCGCCTGATTCAAGATATTCGCTTA<br>ATGTGCTTACCGCTGAAGCAATTTTCAGGTTCGAGAGACC |
| pEPQD0CM0560 | CDS NbUGT76A4 (GenBank ID MT945330), codon optimised for *E. coli* | GGTCTCTAATGAAGATCCAGGAGCGTAAAATGAAGGTTGAGAAGCGCGAATC<br>AGTAGTGCTCGTACCTTACCCGTTTCAAGGTCACCTGACGCCGATGTTACAG<br>CTGGGAAGCATTCTCCACTCGCAGGGTTTCAGCGTGGTCGTAGCCCACACC<br>GAGTTTAACGCGCCTGACTACTCTAATCATCCGGAGTTTGTATTTCACTCGAT<br>GAATGATGGCCTGCAAGGCCGCGATATGTCGATGCCTAGCCTGGAGAACAT<br>GTACGACCTCAATGAAAACTGCAAAGCACCGCTGAAGGACTATCTGGCACGC<br>ATGATGGAAGATAACGGCGACGAGCTGGCCTGCATCGTGTACGATAATGTGA<br>TGTTCTTCGTCGACGACGTAGTAACACAGTTACGTATCCCTTCTATCGTGTTG<br>CGTACCTTTTCTACGACGTACTTACATTCAATGCTGACGATCTTGCAAAAGCC<br>GGATAAGTACCTTCCGTTCGAGGAAAGCCAATTATTAGACCCGCTCCCAGAA<br>CTGCACCCGTTGCGCAGCAAAGATATTCCGTTCCCGGTGATTGACAACACCG<br>TGCCGGAGCCTATCTTGGAGTTTTGCCGCGCCATGTCGGATATCGGGAGCTC<br>CGTGGCCACGATCTGGAATACCATGGAAGATCTTGAAAATAGTTTATTGCTGC<br>GTGTACAAGAGCACTATAAAGTACCGTTCTTCCCGATTGGCCCATTGCATAAG<br>ATGGCGCCGAGCACATCTTCGACGTCTTTGTTAGAGGAAGATGATTCTTGTAT<br>CGAATGGTTAGATAAGCAAGCACCGAACTCTGTGTTGTACGTGTCTTTAGGC<br>AGTCTTGTGAAAATTGACGATAAAGAACTCATCGAAACCGCGTGGGGCTTGG<br>CAAATTCTGAACAACCATTCCTGTGGGTGGTACGTCCGGGAAGCGTTTCGGG<br>ATTCCAGTGGACGGAAGCGCTGCCGGAAGGATTCGAAGAGACTATTGGTGA<br>GCGCGGCCGCATCGTAAAGTGGGCGCCCCAGAAGCAAGTACTGGCCCACCC<br>TGCAGTGGGTGGTTTCTTTACCCACTGCGGCTGGAACAGCACCTTGGAGTCG<br>ATCTGCGAGGAAGTTCCGGTTATTTGTCGTCCGATCCTCGCCGATCAGCCGG<br>TAAATGCGCGCTACCTGTCACAGATTTACATGGTGGGTTTAGAGTTAGAAGC<br>GACCGAGCGCTCAGTGATTGAAAAGACGGTTCGCAAGCTGATGCTCAGCGA<br>GGAAGGCAAGGACGTTAAGAAGCGTGTGGTTGAAATGAAGCAGAAAATTGTA<br>GCGGGCATGCAAATTGACGGCAGTAGCCACAAGAATTTAAACGACCTGGTCA<br>ATTTTATCAGCAGCCTTCCATCACGTACACCCCCGATGCCAGCAGTGGGCGG<br>CATTTTGACGAGCAATTACATTCCTTCGAAAACCATTATTGAATCGGGTTCGA<br>GAGACC |
| pEPQD0CM0561 | CDS NbUGT85A104 (GenBank ID MT945331), | GGTCTCTAATGGGTACCTTGAATCAAACCGAACAGCCGATGAGTAAACCGCA<br>CGCCGTTTGTATCCCCTTCCCGGCCCAGGGTCACATTAACCCAATGTTGAAG<br>CTGGCAAAGCTGCTGCACATTCGCGGTTTCCACATTACGTTTGTGAATACGG<br>ATTTTAATCATCGTCGCTTACTGAAGTCTCGCGGCCCGAACGCGCTGAGTGG<br>TCTGCCGAGTTTCCGCTTTGAATCGATTCCGGACGGCTTACCGCCGTCGAAT<br>GACGATGCCACTCAGGACGTCCCGTCACTGTGCGAATCTTGCACCAAACTGT |

| | | |
|---|---|---|
| | codon optimised for *E. coli* | GCCTTGCGCCCTTCCGTGAGTTGGTTACTCGTTTGAACAACAGTCTGAACTTC<br>CCGCCAGTGACGTGTATCGTAAGTGACGCGGGCATGTCTTTTACACACGAGG<br>TGTCCGAAGAGTTAGGCATCCCGAACGTCGCGTTCTGGACGGCGAGCGGCT<br>GCGCGCTTTGGGCATTTCTGCAGTACCCGAAGCTCGTCGAGGAAGGCTACT<br>GCCCGGTCAAGGACCACTCATATCTGACTAATGGTCACCTTGATACGATCATT<br>GACTGGATCCCGGGGATGGAAGGGATTCGCTTGAAGAACCTTCCCTCATTTA<br>TTCGCTCAACCGTAGATGAGCCGTCTTACATGGTTATTAAGTTCATCATGGAA<br>GAGATTCTGGATAAGATCCCGAAGGCATCGGCGTTGATCCTGAATACCTTTG<br>ACGCATTGGAAACCGACGTCCTTAAACCGATCTTAACGCTTTTCCCGACAGT<br>GTACACCCTGGGCCCGTTCCATACAAGCCTGAACAACCAGACCCAGGATGA<br>GGACCTTAAGAGCATCGGGAGCAACTTGTGGAAAGAGGACACACATTGCCTT<br>GAGTGGCTTAATACAAAGAAGCCTAACTCTGTTGTTTACGTCAACTTCGGCTC<br>GATCACCGTTCTGAGTCCTAAGCAACTTGTGGAGTTCGCTTGGGGTTTAGCA<br>AACTCGAAGCTGAACTTTCTGTGGATCATCCGCTCCGACATCGTAAAAGGCG<br>ACAGTCTGATTCTCCCGCCGGAGCTGCTTGCTGAAATCAAAGAGCGCGGACT<br>CTTGTGCGGCTGGTGTCCACAGGAGCACGTCCTGTGCCATCCGAGCGTGGG<br>CGGCTTCTTGACGCATTGCGGCTGGAACAGCACGTTCGAGAGCATTAGTTTC<br>GGCGTCCCGATGCTGTGCTGGCCGTTCTTCGCCGACCAGCAGACCAACTGC<br>TGGTTCATCTGCAACTGTCTTGGAGTGGGTATGGAGATCGACTCTCACGTTA<br>AACGTGAGGTTATCGAGGAGTTGGTGAAGGAACTCATGATCGGGGAGAAGG<br>GAATCGAACTGAAGGAGAACGCGCTCAAATGGAAACGTCTGACCGAGAAAAC<br>CATTTCATCACCCGACGGCAGCTCGTACATGAACTTCGATAATCTGGTATCTC<br>ACGTTCTTCTCCGTAAAGACTCATCCTTTCTCACTTGCGGTTCGAGAGACC |
| pEPYC0CM0470 | CDS AtTGA1 (AT5G65210.2), contains a BbsI site | GGTCTCgAATGAATTCGACATCGACACATTTTGTGCCACCGAGAAGAGTTGGT<br>ATATACGAACCTGTCCATCAATTCGGTATGTGGGGGGAGAGTTTCAAAAGCA<br>ATATTAGCAATGGGACTATGAACACACCAAACCACATAATAATACCGAATAAT<br>CAGAAACTAGACAACAACGTGTCAGAGGATACTTCCCATGGAACAGCAGGAA<br>CTCCTCACATGTTCGATCAAGAAGCTTCAACGTCTAGACATCCCGATAAGATA<br>CAAAGACGGCTTGCTCAAAACCGCGAGGCTGCTAGGAAAAGTCGCTTGCGC<br>AAGAAGGCTTATGTTCAGCAACTGGAAACAAGCAGGTTGAAGCTAATTCAATT<br>AGAGCAAGAACTCGATCGTGCTAGACAACAGGGATTCTATGTAGGAAACGGA<br>ATAGATACTAATTCTCTCGGTTTTTCGGAAACCATGAATCCAGGGATTGCTGC<br>ATTTGAAATGGAATATGGACATTGGGTTGAAGAACAGAACAGACAGATATGTG<br>AACTAAGAACAGTTTTACACGGACACATTAACGATATCGAGCTTCGTTCGCTA<br>GTCGAAAACGCCATGAAACATTACTTTGAGCTTTTCCGGATGAAATCGTCTGC<br>TGCCAAAGCCGATGTCTTCTTCGTCATGTCAGGGATGTGGAGAACTTCAGCA<br>GAACGATTCTTCTTATGGATTGGCGGATTTCGACCCTCCGATCTTCTCAAGGT<br>TCTTTTGCCACATTTTGATGTCTTGACGGATCAACAACTTCTAGATGTATGCAA<br>TCTAAAACAATCGTGTCAGCAAGCAGAAGACGCGTTGACTCAAGGTATGGAG<br>AAGCTGCAACACACCCTTGCGGACTGCGTTGCAGCGGGACAACTCGGTGAA<br>GGAAGTTACATTCCTCAGGTGAATTCTGCTATGGATAGATTAGAAGCTTTGGT<br>CAGTTTCGTAAATCAGGCTGATCACTTGAGACATGAAACATTGCAACAAATGT<br>ATCGGATATTGACAACGCGACAAGCGGCTCGAGGATTATTAGCTCTTGGTGA<br>GTATTTTCAACGGCTTAGAGCCTTGAGCTCAAGTTGGGCAACTCGACATCGT<br>GAACCAACGggTTCGTGAGACC |
| pEPYC0CM0471 | CDS AtTGA2 (AT5G06950.1), domesticated to remove BsaI and BbsI sites | GGTCTCAAATGGCTGATACCAGTCCGAGAACTGATGTCTCAACAGATGACGA<br>CACAGATCATCCTGATCTTGGGTCGGAGGGAGCACTAGTGAATACTGCTGCT<br>TCTGATTCGAGTGACCGATCGAAGGGAAAGATGGATCAAAAGACTCTTCGTA<br>GGCTTGCTCAAAACCGTGAGGCAGCAAGGAAAAGCAGATTGAGGAAGAAGG<br>CTTATGTTCAGCAGCTAGAGAACAGCCGCTTGAAACTAACCCAGCTTGAGCA<br>GGAGCTGCAAAGAGCAAGACAGCAGGGCGTATTCATTTCAGGCACAGGTGA<br>CCAGGCCCATTCTACTGGTGGAAATGGTGCTTTGGCGTTTGATGCTGAACAT<br>TCACGGTGGTTGGAAGAAAAGAACAAGCAAATGAACGAGCTGAGGTCTGCTC<br>TGAATGCGCATGCAGGTGATTCTGAGCTTCGAATAATAGTCGATGGTGTGAT<br>GGCTCACTATGAGGAGCTTTTCAGGATAAAGAGCAATGCAGCTAAGAATGAT<br>GTCTTTCACTTGCTATCTGGCATGTGGAAAACACCAGCTGAGAGATGTTTCTT<br>GTGGCTCGGTGGATTTCGTTCATCCGAACTTCTAAAGCTTCTGGCGAATCAG<br>TTGGAGCCAATGACAGAGAGACAGTTGATGGGCATAAATAACCTGCAACAGA<br>CATCGCAGCAGGCTGAAGATGCTTTGTCTCAAGGGATGGAGAGCTTACAACA<br>GTCACTAGCTGATACTTTATCGAGCGGGACTCTTGGTTCAAGTTCATCAGGG<br>AATGTCGCAAGCTACATGGGTCAGATGGCCATGGCAATGGGAAAGTTAGGTA<br>CACTCGAAGGATTTATCCGCCAGGCTGATAATTTGAGACTACAAACATTGCAA<br>CAGATGATAAGAGTATTAACAACGAGACAGTCAGCACGTGCTCTACTTGCAAT<br>ACACGATTACTTCTCACGGCTACGAGCTCTAAGCTCCTTATGGCTTGCTCGAC<br>CCAGAGAGggTTCGCGAGACC |

| | | |
|---|---|---|
| pSD0KN04 | CDS TaLUXA,<br>codon optimised<br>for *E. coli*<br><br>TRIAE_CS42_3AL_TGACv1_194142_AA0627410.1 | GGTCTCTAATGGGGGAAGAAGCAGGCGGATATGGCTTTGATTTTGGCGGCG<br>GCGGCGGTGGATACGGTGGCTATGATGGCCGTGTGACTGAATGGGAAACCG<br>GGTTGCCTGGCTGTGATGAATTAACGCCTCTGTCCCAACCCTTGGTCCCGCC<br>TGGATTGGCGGCCGCCTTTCGTATTCCACCTGAACCCGGCCGTACGCTGTTG<br>GATGTACATCGTGCTAGTAGCGCGACGGTTAGCCGTCTGCGTAGCACATCAT<br>CCAGTCCTAGCTCAGGTAATGGGCATGCTGGAACTCATGCGAATGGTGGAAG<br>TTTTCCTTCGTTTCCGGGGAAAGGTGCCGCTGCTGGCGATGATTCTGGTAAT<br>CGTGATAATAATAGCGCAGAATCCGCGGGCGAGAAAGCTGCCGCAACAAAA<br>CGGGCACGTTTGGTCTGGACACCCCAACTCCATAAACGTTTTGTTGAAGTAG<br>TAGCTCATCTCGGAATTAAAAGTGCTGTTCCTAAAACGATTATGCAACTCATG<br>AATGTCGAAGGTTTAACGCGTGAAAATGTTGCGTCACATTTACAAAAGTATCG<br>TCTGTATGTAAAACGAATGCAAGGTCTGTCTAATGAAGGTCCTTCGGCGAGC<br>GATCATATTTTCGCCAGTACCCCTGTTCCCCCGTCGCTTCGTGAACCGCAAG<br>TTCCTCATGCTGCAGCAATGGCACCTGCGATGTATCATCATCATCCAGCACC<br>GATGGGCGGTGTAGCGGCAGGACATGGCGGCTATTATCAACAACAACATTCA<br>GGGCATGCTGTATATAACGGTTATGGTGGACATGGACATGGTGGCGGGGTTT<br>CATCATATCCACATTATCATCATGGAGATCAAGGTTCGGGAGACC |
| pSD0KN05 | CDS TaLUXB,<br>codon optimised<br>for *E. coli*<br><br>TRIAE_CS42_3B_TGACv1_220755_AA0718150.1 | GGTCTCTAATGGCGGGAACAACAGGTGGTTGGCGTAGCGGTCGTCGTGCAT<br>GTCGTGGAGCCACCAGTCTGCCTGGTTGTGATGAATTGACGCCCCTCTCTCA<br>ACCCCTCGTTCCGCCCGGTTTGGCGGCTGCCTTTCGTATTCCGCCAGAACCT<br>GGCCGGACCCTTCTTGATGTCCATCGTGCATCGTCTGCGACTGTTTCGCGTC<br>TGCGTTCGGCATCATCTAGTCCTTCAAGCGGGAATGGGCATGCGACCGGTG<br>GTGGTAGCTTTCCATCATTTCCGGGTAAAGCAGCCGCCGCTGCTGAAGCTGG<br>TGAAGATAGTGGTAATCGCGATAATAATTCAGCAGAAAGTGGTGGCGATAAA<br>AGTGCTGCTGCCGCAACCAAACGAGCTCGCCTCGTTTGGACTCCACAATTGC<br>ATAAACGTTTTGTCGAAGTTGTAGCCCATCTTGGCATTAAATCAGCAGTCCCG<br>AAAACGATTATGCAATTGATGAATGTAGAAGGTCTTACGCGTGAAAATGTGGC<br>AAGTCATCTGCAAAAGTATCGTCTGTATGTGAAACGTATGCAAGGTCTTAGCA<br>ATGAAGGCCCTAGCGCGTCAGATCATATTTTCGCATCAACTCCAGTACCGCC<br>TTCACTGCGTGAACCGCAAGTTCCAGTTCCCCATGCAGCTGCGATGGCACCA<br>GCAATGTATCATCATCATCCAGCTCCTATGGGTGGGGTTGCGGCAGGTCATG<br>GTGGTTATTATCAACAACAACATTCCGGTCATGCTGTGTATAACGGCTATGGT<br>GGCCATGGTCATGGTGGTGGTGTTTCGTCTTACCCACATTATCATCATGGAG<br>ATCAAGGTTCGGGAGACC |
| pSD0KN06 | CDS TaLUXD,<br>codon optimised<br>for *E. coli*<br><br>TRIAE_CS42_3DL_TGACv1_250132_AA0862670.1 | GGTCTCTAATGGGGGAAGAAGCAGGCGGTTATGATTTCGATTTTGGCGGCG<br>GCGGTGGCGGCTACGGTGGCTATGATGGCCGGGTTACCGAATGGGAAACTG<br>GTTTGCCGGGCTGTGATGAATTAACGCCTCTGAGTCAACCTCTCGTTCCGCC<br>CGGTTTAGCAGCTGCCTTTCGTATTCCACCCGAACCTGGTCGTACCCTGTTA<br>GATGTTCATCATGCATCAAGCGCGACGGTTAGCCGTCTGCGTTCAGCGAGCA<br>GCGGTAATGGGCATGCGGGAACTGGTGCAAATGGAGCGTCATTTCCTTCATT<br>TCCTGGTAAAGGTGCAGCTGCAGGCGACGATTCTGGTAATCGTGATAATAAT<br>AGCGCAGAATCTGCAGGTGAGAAAAGCGCTGCCGCCAAACGCGCACGATTG<br>GTCTGGACCCCACAATTGCATAAACGTTTTGTCGAAGTTGTCGCACATTTGGG<br>TATTAAATCAGCAGTTCCGAAAACGATTATGCAACTCATGAATGTTGAAGGTT<br>TAACACGTGAAAATGTTGCAAGTCATTTGCAGAAATATCGTTTGTATGTGAAA<br>CGTATGCAAGGTCTGAGTAATGAAGGTCCTTCTGCTAGTGATCATATCTTTGC<br>GTCAACCCCTGTTCCCCCATCCTTGCGTGAACCACAAGTTCCGCATGCTGCA<br>GCAATGGCGCCGGCTATGTATCATCATCATCCTGCACCTATGGGCGGCGTG<br>GCGGCGGGGCATGGCGGTTATTATCAACAACAACATAGTGGTCATGCAGTTT<br>ACAATGGATACGGTGGTCATGGTCATGGTGGCGGGGTATCATCGTATCCTCA<br>TTATCATCATGGAGATCAAGGTTCGGGAGACC |
| pHR0CM03 | CDS TaLHYA,<br>codon optimised<br>for *E. coli*<br><br>TraesCS7A02G299400.2 | GGTCTCAAATGGAAATCAACAGTAGCGGGGAAGAGACTGTCATTAAAGTCCG<br>TAAACCATATACCATTACCAAACAACGTGAACGTTGGACGGAAGCAGAACATA<br>AGCGCTTTCTGGAAGCGCTGAAGTTGTACGGTCGTGCATGGCAACGTATCGA<br>GGAACACGTGGGAACTAAAACCGCGGTACAGATTCGTTCGCACGCACAGAAA<br>TTCTTTACGAAACTGGAGAAAGAGGCCATTAACAACGGCACATCCCCAGGTC<br>AGGCACACGACATCGATATTCCGCCGCCTCGCCCGAAGCGCAAGCCTAATT<br>GCCCGTACCCCCGTAAGGGCTGCTTAAGCAGCGAAACGCCAACGCGCGAGG<br>TCCCTAAGAGCAGCGTTTCCCTGTCGAACAGTAACGCCGAGATGGCATCCAA<br>CGGTACCTTACAATTGACATGTATCCGCAAGCTGCAGCGTAAAGAACTGAGC<br>GAGAATGGGAGCTGTAGCGAGGTGATCAACATTTTCCGCGAGGCCCCGAGT<br>GCAAGCTTCAGCTCATCGAACAAATCTAGCAGTAACCACGGCGTGTCAGGCG<br>GCATCGAGCCTACCAAGACTGAGAATAAGGATATCGCGACTATGGAGCGTAA<br>GTCGACGAGTATCGACGTCGGCAAAGACGTCAAGGACATCAACGATCAAGA<br>GATGGAACGCAATAATCGTGTACATATTAGCTCAAACTACGATGGATCCCACG<br>AGGACTGCTTAGACAATTCCATGAAGCATATGCAGCTGAAACCCAACACCGT<br>TGAAACGACTTATACCGGGCAGCACGCCGCAAGCGCTCCCTTATATCAGATG |

| | | |
|---|---|---|
| | | AACAAAACGGGAGCCACGGGTGCACCGGATCCAGGGACGGAAGGCTCTCAC<br>CCAGACCAGACCTCTGACCGCGTTGGCGGCGCAAACGGCTCAATGGATTGT<br>ATTCACCCGACCCTGCCTGTAGACCCGAAGATCGGTTCATCTAGCACTGCGC<br>AAAGCTTCCCACATAATTACGCCGGGTTCGCCCCTACCATGCAGTGTCACTG<br>TAATCAGGACGCATATCGTTCGAGCCTGAACATGAGCAGTACATTTTCTAATA<br>TGCTGGTGTCTACTCTGCTCAGCAATCCTACCGTGCACGCTGCGGCGCGTCT<br>GGCCGCTAGCTATTGGCCGGCCGCAGATAGTAATATCCCAGTGGGCCCGAA<br>CCAGGAAGTGTTCGCCGAAAATGCCCAAGGCCGTCACATCGGAAGTCCGCC<br>CTCAATGGCGAGTGTAGTGGCGGCTACCGTGGCGGCAGCTTCAGCCTGGTG<br>GGCCACGCAAGGATTGCTTCCCCTGTTTGCGCCGCCTATGGCATTTCCCTTC<br>GTTCCTGTCCCGACCGCCTCATTCCCTACGACAGACGTGCAACGCGCCACC<br>GAAAATTGTCCCGTAGATAATGCGCCAAAAGAGTGTCAGGTCGCCCAAGAAC<br>AGGGACAGCCGGAAGCAATGATTGTGGTGGCAAGCAGCGGGAGTGATAAAA<br>GCGGTAAGGGCGAAGTTAGTCCGCATACAGAACTTAACATTAGCCCTGCGGA<br>CAAGGTAGAAACCACGCCCCCGACGGGTGCGGAAACCTCGGACGCCTTTGG<br>TAATAAGAAGAAACAAGACCGTAGTTCCTGCGGCTCGAATACCCCGAGCTCC<br>TCTGACGTCGAAGCCGAGCACGTGCCGGAAAATCAGGACCAGGCGAATGAT<br>AAAACCCAACAGGCTTGCTGTAGCAACAGCAGCGCAGGCGATATGAATCATC<br>GTCGCTTCCGCAATATCAGTTCTACGAACGACAGCTGGAAAGAGGTATCAGA<br>GGAAGGGCGCATGGCGTTTGACAAGTTGTTTTCCCGTGGTAAACTGCCACAG<br>TCCTTCTCACCGCCGCAGGCCGAGGGTTTAAAAGTTGTTCCGCGCGGCGAG<br>CAGGATGAGGCAACCACCGTAACCGTTGATCTGAATAAATCTGCAGCTGTAA<br>TGGATCACGAGCTGGACACCCTTGTAGGCCCGCGTGCCACGTTCCCGATCG<br>AGCTGAGCCATCTTAACATGAAAAGCCGTCGTACCGGTTTTAAGCCGTATAAA<br>CGCTGTAGTGTAGAGGCCAAAGAAAACCGCGTCCCTGCCGCAGATGAAGTC<br>GGCACGAAACGCATCCGTTTGGATTCGGAGCCGTCAACCGGTTCGTGAGAC<br>C |
| pHR0CM02 | CDS TaNAMA1, codon optimised for *E. coli*<br><br>TraesCS6A02G 108300.2 | GGTCTCAAATGCGTAGTATGGGTTCGAGTGATAGTTCCAGCGGGTCAGCCCA<br>GAAGGCGGCCCGTCACCAACACGAACCTCCTCCGCCACGTCAACGCGGGAG<br>CGCACCAGAATTGCCGCCTGGTTTTCGCTTTCATCCTACCGATGAAGAACTT<br>GTTGTCCATTATCTTAAGAAGAAAGCAGCGAAAGTTCCTCTTCCGGTAACGAT<br>TATTGCGGAAGTCGACCTGTATAAATTTGATCCCTGGGAACTGCCTGAAAAAG<br>CTACGTTTGGAGAACAAGAATGGTATTTCTTTTCTCCCCGTGATCGTAAATAT<br>CCGAATGGTGCCCGACCTAATCGCGCCGCAACCAGCGGTTATTGGAAAGCA<br>ACTGGTACTGATAAGCCGATTTTGGCGTCAGGTACTGGCTGTGGACTTGTAC<br>GTGAAAAGCTGGGTGTAAAGAAAGCCTTGGTCTTTTATCGTGGTAAACCACC<br>AAAAGGTTTAAAGACGAATTGGATTATGCATGAATATCGTTTAACGGATGCGA<br>GTGGTAGTACGACTACGTCGCGCCCGCCTCCTCCGGTCACGGGCGGTTCCC<br>GTGCAGCCGCGAGTCTGCGTGTTCGCACTCGCGTTGATCGTACCCTGGATG<br>ATTGGGTCCTTTGTCGTATTTATAAGAAGATTAATAAAGCAGCAGCCGGGGAC<br>CAACAACGGTCTACTGAATGTGAAGATTCAGTAGAAGATGCCGTAACTGCATA<br>TCCACTGTACGCAACTGCAGGTATGGCAGGCGCTGGAGCACACGGTTCTAAT<br>TATGCCTCGCCCTCTTTATTGCATCACCAAGATTCTCACTTTCTTGAAGGTTTG<br>TTTACGGCCGATGATGCGGGACTTTCTGCCGGGGCGACAAGCTTATCGCATC<br>TTGCAGCAGCAGCACGTGCCTCACCCGCGCCAACGAAGCAATTCCTGGCAC<br>CAAGCAGCTCGACACCCTTTAATTGGCTGGACGCATCTCCTGCAGGGATTCT<br>TCCTCAAGCCCGCAACTTTCCAGGATTCAATCGTTCGCGAAATGTGGGGAAC<br>ATGTCACTTAGCAGCACAGCGGATATGGCGGGTGCCGCAGGTAACGCTGTT<br>AATGCAATGTCAGCGTTCATGAACCCGCTGCCAGTACAGGATGGCACATATC<br>ACCAGCATCACGTGATCTTAGGTGCGCCTCTCGCCCCGGAAGCCACGACCG<br>GCGGAGCAACTTCCGGCTTCCAACACCCGGTTCAGGTGAGCGGAGTAAATT<br>GGAACCCTGGTTCGTGAGACC |
| pEPKK0CM0195 | chrysanthemyl diphosphate synthase (CcCPPase) from *Chrysanthemum cinerariaefolium* P0C565.2, codon optimised for *E. coli* | GGTCTCTAATGACCACCACTTTATCGTCCAACCTCGATAGCCAGTTTATGCAG<br>GTCTACGAAACCCTTAAGTCAGAACTGATCCACGATCCTAGCTTCGAATTCGA<br>CGATGACAGCCGCCAATGGGTTGAACGCATGATCGATTATAACGTGCCAGGT<br>GGGAAAATGGTTCGCGGTTACAGCGTGGTCGATAGTTATCAGCTCTTGAAGG<br>GCGAGGAGCTGACAGAGGACGAGGCCTTCCTTGCATGCGCCTTAGGCTGGT<br>GTACCGAGTGGCTGCAGGCGTTCATTCTGGTGCTCGACGATATTATGGACGG<br>TAGCCATACTCGCCGTGGGCAGCCGTGCTGGTTCCGCCTTCCAGAAGTGGG<br>CGTGGTGGCGATCAACGACGGCGTACTCTTACGTAATCACGTTCACCGCATT<br>TTAAAGAAGTACTTTCAGGGCAAACCGTACTATGTTCATCTGCTCGATTTATTT<br>AACGAGACTGAGTTCCAGACTATTTCAGGCCAGATGATCGACACTATCTGCC<br>GCCTGGCGGGTCAGAAGGACTTGTCGAAATACACCATGACCTTAAATCGCCG<br>TATCGTGCAATATAAGGGCAGCTATTATTCCTGCTATCTGCCTATCGCATGCG<br>CGTTATTAATGTTCGGTGAAAACTTGGAAGATCACGTGCAGGTTAAGGATATT<br>CTGGTGGAGCTGGGCATGTACTACCAGATCCAAAACGACTACCTGGATACCT<br>TCGGCGACCCGGACGTGTTCGGTAAAACTGGTACGGACATCGAGGAGTGTA |

| | | |
|---|---|---|
| | | AATGCTCTTGGCTTATCGCTAAGGCCTTGGAGCTGGCAAATGAGGAGCAGAAGAAGATCCTGAGTGAGAATTACGGTATTAATGACCCTAGCAAAGTTGCGAAGGTTAAAGAGCTGTATCACGCGCTGGACCTTAAAGGAGCATACGAGGACTACGAAACCAACTTGTACGAAACCAGTATGACGAGTATCAAGGCACACCCCAATATCGCGGTTCAGGCAGTCCTCAAGAGCTGCTTAGAGAAAATGTACAAAGGCCACAAAGGTTCGAGAGACC |
